# A Robust Bayesian Approach to Bulk Gene Expression Deconvolution with Noisy Reference Signatures

**DOI:** 10.1101/2022.10.25.513800

**Authors:** Saba Ghaffari, Kelly J. Bouchonville, Ehsan Saleh, Remington E. Schmidt, Steven M. Offer, Saurabh Sinha

## Abstract

**Background:** Differential gene expression in bulk transcriptomics data can reflect change of transcript abundance within a cell type and/or change in the proportion of cell types within the sample. Expression deconvolution methods can help differentiate these scenarios and enable more accurate inference of gene regulation by estimating the contributions of individual cell types to bulk transcriptomic profiles. However, the accuracy of these methods is sensitive to technical and biological differences between bulk profiles and the cell type-signatures required by them as references.

**Results:** We present BEDwARS, a Bayesian deconvolution method specifically designed to address differences between reference signatures and the unknown true signatures underlying bulk transcriptomic profiles. Through extensive benchmarking utilizing eight different datasets derived from pancreas and brain, we demonstrate that BEDwARS outperforms leading in-class methods for estimating cell type proportions and signatures. Furthermore, we systematically show that BEDwARS is more robust to noisy reference signatures than all compared methods. Finally, we apply BEDwARS to newly generated RNA-seq and scRNA-seq data on over 100 induced pluripotent stem cell-derived neural organoids to study mechanisms underlying a rare pediatric condition (<u>D</u>ihydro<u>p</u>yridine <u>D</u>ehydrogenase deficiency), identifying the possible involvement of ciliopathy and impaired translational control in the etiology of the disorder.

**Conclusion:** We propose a new approach to bulk gene expression deconvolution which estimates the cell type proportions and cell type signatures simultaneously and is robust to commonly seen mismatches between reference and true cell type signatures. Application of our method lead to novel findings about mechanisms of a rare pediatric condition.

## Background

RNA sequencing (RNA-seq) is the cornerstone of regulatory genomics studies. It provides information on changes in gene expression accompanying a biological process and allows for the reconstruction of gene regulatory networks. The widespread adoption and utility notwithstanding, traditional “bulk” RNA-seq technologies offer an incomplete and potentially biased view of expression changes, especially for heterogeneous tissues, since they report gene expression levels aggregated across multiple cell types present in unknown proportions. In data from heterogeneous samples, differential gene expression can reflect a regulated change of transcript abundance within a cell type, a change in the proportion of cell types within the sample, and/or a combination of both phenomena. Differentiating these scenarios is important for inferring mechanisms surrounding biological processes.

Single-cell transcriptomics techniques, such as single-cell RNA-seq (scRNA-seq), can address this issue directly, but are considerably more expensive and time consuming, limiting the use of these high-resolution assays to fewer samples per biological condition than is afforded by bulk technologies. This practical consideration underlies the need to develop reliable computational methods to deconvolve bulk transcriptomics profiles. Deconvolution methods (1–3) have the potential to reveal cell type-resolution transcriptomes at the low cost and large scale afforded by bulk RNA-seq, allowing greater statistical power in detecting transcriptomic and compositional changes in a biological process.

Deconvolution methods assume that a bulk RNA-seq profile is a weighted mixture of cell type-specific profiles, known as “signatures”, and use statistical techniques to estimate the weights and/or cell type signatures that comprise the bulk profile. While it may be possible to estimate both simultaneously, a more practical approach is to rely on reference signatures (cell type-specific expression profiles from similar biological conditions) to estimate cell type proportions. CIBERSORT (4) and FARDEEP (5) adopt this approach. A similar approach is used by MuSIC (6), SCDC (7), and BISQUE (8), which utilize entire existing scRNA-seq data sets as reference. While these methods demonstrate the potential of this approach, they also highlight challenges that arise due to technical and biological differences between reference signatures and bulk transcriptomic profiles. For instance, reference cell type signatures obtained from scRNA-seq data may be unsuitable for deconvolving bulk RNA-seq data due to a difference in technologies, even if they profile the same biological conditions. Similarly, reference signatures derived from healthy subjects used for deconvolving patient transcriptomics profiles or those from untreated biospecimens used for experimentally perturbed samples may introduce unknown biological biases, leading to errors in deconvolution. Tissue-specific differences in cell type transcriptomes may also lead to such errors.

Recently, comprehensive benchmarking studies have detailed the extent to which biologically and/or technologically mismatched reference signatures can affect the accuracy of deconvolution methods. For instance, Sutton et al. (9) observed a negative impact of biological differences on deconvolution accuracy across all methods tested. Newman et al. (10) propose the use of batch correction to bridge the gap between transcriptomics technologies, while Jew et al. (8) suggest learning a transformation between synthetic bulk profiles generated from a reference scRNA-seq data set and the target bulk data set, which can be used for deconvolution. Sutton et al. propose using reference signatures that are aggregated from multiple sources and technologies, while Wang et al. (6) and Dong et al. (7) utilize heterogeneity across multiple single cell data sets to improve the accuracy of deconvolution. Despite these efforts, deconvolution in the face of mismatched references signatures remains an unsolved problem.

In this work, we describe a rigorous Bayesian probabilistic method for bulk expression deconvolution, called BEDwARS (Bayesian Expression Deconvolution with Approximate Reference Signatures), which tackles the problem of signature mismatch from a complementary angle. It does not assume availability of multiple reference signatures, nor does it rely solely on transformations of data prior to deconvolution. Instead, it incorporates the possibility of reference signature mismatch directly into the statistical model used for deconvolution, using the reference to estimate the true cell type signatures underlying the given bulk profiles while simultaneously learning cell type proportions. It assumes that each bulk expression profile is a weighted mixture of cell type-specific profiles (“true signatures”) that are unknown but not very different from given reference signatures. Thus, the reference signatures are used as priors in a Bayesian estimation of true signatures, along with the cell type proportions. Our strategy of jointly inferring both proportions and signatures is a notable departure from the two-step strategy of current methods whereby reference signatures are first “corrected” and then used for deconvolution. It has parallels to so-called “full deconvolution” methods (11,12) but its ability to utilize given reference signatures distinguishes it from this genre of methods. Moreover, our technique works with reference cell type signatures from any source and is not limited to scRNA-seq references.

We demonstrate the advantages of BEDwARS through extensive tests on semi-synthetic data sets mimicking human pancreatic islet and brain gene expression data, under varying levels of misalignment between reference and true signatures. We evaluate its ability to recover cell type-specific expression signatures as well as sample-specific cell type compositions in comparison to state-of-the-art reference signature-based deconvolution methods (1). In these tests, BEDwARS outperforms leading methods such as CIBERSORT, CIBERSORTx and FARDEEP in the estimation of cell type proportions. Furthermore, it provides more accurate estimates of true cell type signatures compared to RODEO, a state-of-the-art expression deconvolution method that estimates such signatures based on cell type proportions inferred by methods such as CIBERSORTx or FARDEEP. Our evaluations demonstrate the advantage of jointly inferring cell type proportions and cell type-specific signatures while allowing the latter to deviate from pre-determined reference signatures that may not be accurate for the bulk data being studied. Finally, we apply BEDwARS to study the mechanisms underlying pediatric <u>D</u>ihydro<u>p</u>yridine <u>D</u>ehydrogenase (DPD) deficiency, based on new data from induced pluripotent stem (iPS) cell-derived neural organoids.

## Results

### Overview of BEDwARS

BEDwARS is a Bayesian approach to deconvolving bulk expression profiles using reference expression profiles (“signatures”) of the constituent cell types. It is designed to be robust to “noise” in provided reference signatures that may arise due to biological and/or technical differences from the bulk expression profiles. The underlying model assumes, like other deconvolution models, that the bulk expression profile, say ***X***, of a biological sample is a weighted mixture of cell type-specific signatures, say ***S***_***c***_ (for each cell type *c*). Loosely speaking, ***X*** = ∑_*c*_ *w*_*c*_***S***_***c***_, where ***X*** and ***S***_***c***_ are *G*-dimensional expression profiles (*G* is the number of genes) and *w*_*c*_ is the proportion of cell type *c* in the sample (**Figure 1**). Importantly, the BEDwARS model assumes that a cell type’s signature ***S***_***c***_, henceforth called the “true signature” of cell type *c*, is similar to but not identical to the available reference signature, say 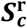, of that cell type, and must be estimated as part of the deconvolution process. This is the fundamental conceptual difference of BEDwARS from existing approaches. In other words, given a collection of bulk expression profiles {***X***} and a set of reference signatures 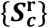, BEDwARS simultaneously infers the proportions ***w*** = {*w*_*c*_} of all cell types *c* in each sample, and the unknown “true signatures” {***S***_***c***_} of all cell types while maintaining similarity between reference signatures and inferred true signatures.

**Figure 1.**
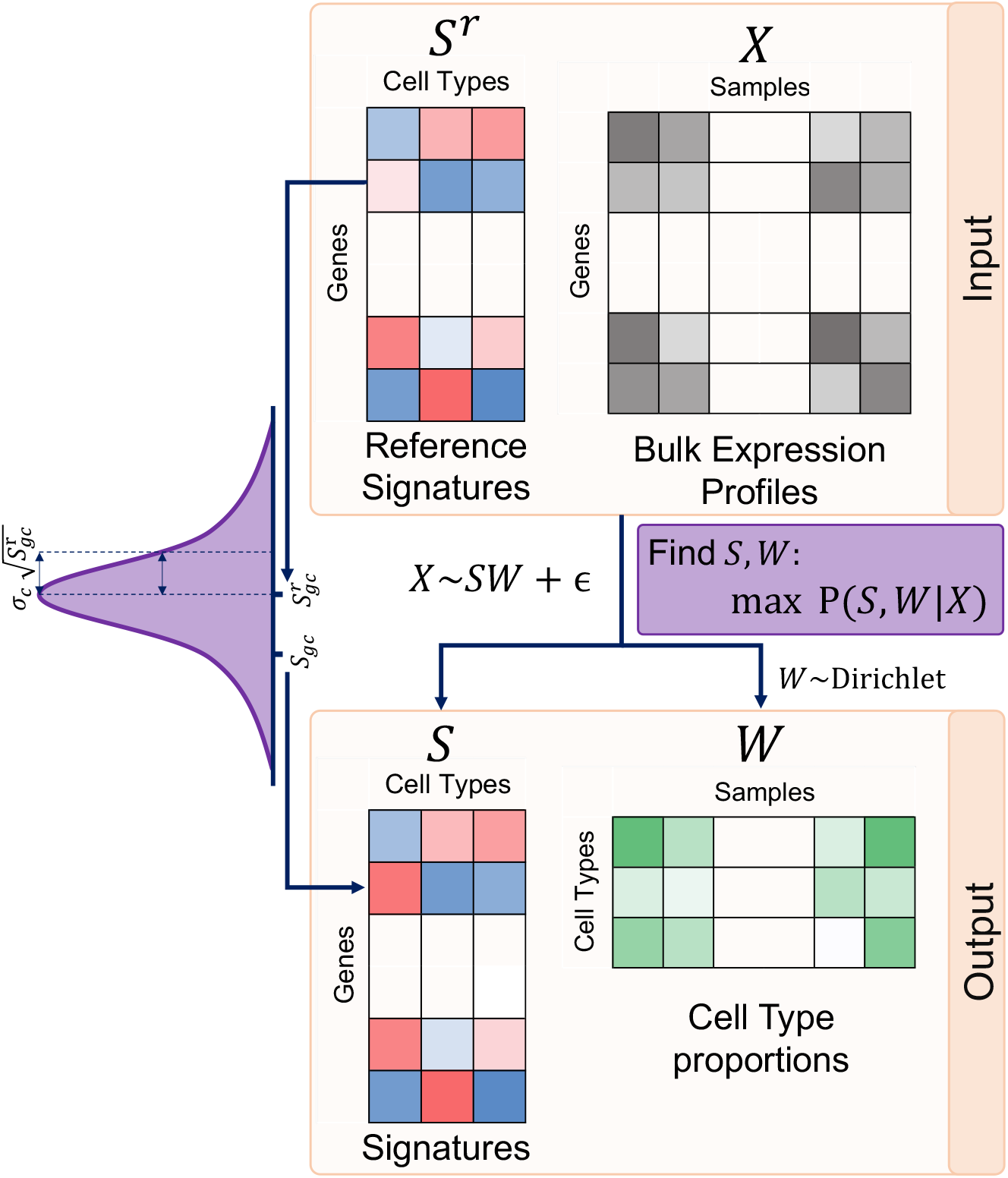
Model outline. BEDwARS takes as input the bulk expression profiles (*X*) as well as the reference signatures of individual cell types (*S^r^*). BEDwARS models bulk profiles (*X*) as combinations of “true” but unknown signatures (*S*) of cell types mixed in unknown proportions (*W*) and estimates both *S* and *W* from data. The true signatures are assumed to be similar to the reference signatures and differences between them are assumed to be normally distributed with mean zero and variance proportional to the reference gene expression 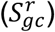. The constant of proportionality is a cell-type specific parameter (*σ_c_*) that allows for the degree of differences to vary across the cell types. The unknown cell type proportions (*W*) are assumed to follow a Dirichlet distribution. Maximum *a posteriori* estimation is used to find the cell type proportions and signatures based on the data.

To understand the model-prescribed relationship between the provided reference signatures and unknown true signatures (**Figure 1**), consider the expression of gene *g* in cell type *c*, as per the true signature (*S*_*gc*_) and reference signature 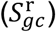. The model assumes that *S*_*gc*_ differs from 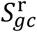 by an amount that is Gaussian distributed with mean 0 and a gene- and cell type-dependent variance. In particular, the variance of this “noise” term reflecting the difference between the true and reference signature values is proportional to gene’s reference expression 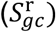 and the constant of proportionality is cell type-dependent. (The noise is modeled on log-transformed expression, see Methods.) Thus, the model prefers the true signature value *S*_*gc*_ to be similar to the reference 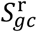 but also allows them to diverge, and the divergence can be larger for more highly expressed genes, following a conventional assumption of gene expression models (13). Furthermore, since biological differences in expression profiles may manifest to different extents in different cell types, the model allows the divergence to be greater for some cell types than for others. The extent of divergence, reflected in the variance term, is learnt from the data.

The above design principles underlie how BEDwARS assigns a probability Pr(***X***| {***S***_***c***_}, ***w***) to the bulk expression profile ***X*** conditional on a specific estimation of true signatures {***S***_***c***_} and cell type proportions ***w***, and how it assigns prior probabilities to ***S***_***c***_ based on reference signature 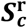 (see Methods). The tool estimates the true signatures and cell type proportions with greatest posterior probability Pr({***S***_***c***_}, ***w***|***X***), using Metropolis Hastings sampling. The calculations include *C* + 2 additional parameters (where *C* is the number of cell types), which are simultaneously optimized. Importantly from a usability perspective, no parameters require hand-tuning and the preset parameters used in the model were the same for all experiments performed in this study (see Methods for a more precise description of the model and optimizations.)

### BEDwARS deconvolution of human pancreatic islet transcriptomic profiles is robust to mismatched and noisy reference signatures

We assessed the accuracy of bulk expression deconvolution by BEDwARS following benchmarking practices established by recent publications (1,9) and making use of eight different transcriptomics data sets (**Table 1)**. The overall approach to these evaluations is the following: (1) begin with a single-cell transcriptomics data set with labeled cell types and aggregate the transcriptomes of heterotypic cells to create a “pseudo-bulk” transcriptomic profile, keeping track of the relative proportions of different cell types in the aggregate; repeats of this process results in multiple pseudo-bulk profiles that form the “target” data set to be deconvolved, (2) select a suitable transcriptomics data set representing a biological condition similar to the target data set and wherefrom we can derive an expression profile for each cell type; this is the collection of “reference signatures”, (3) deconvolve the target data set using the reference signatures and a method of choice, and compare the recovered proportions of different cell types to their true values from step 1. We compared the performance of BEDwARS to leading available tools – FARDEEP (5), CIBERSORT (4) and CIBERSORTx (10). FARDEEP and CIBERSORT were the best performing deconvolution tools from the benchmarking study of Cobos et al. (1). CIBERSORTx was also included because it improves upon CIBERSORT by performing batch correction to account for technical differences between the target data set and reference signatures.

**Table 1.** Summary of datasets used for benchmarking.

| Dataset | Tissue Type | Data Type | Sequencing Protocol | Species |
| --- | --- | --- | --- | --- |
| Baron | pancreatic islet | scRNA-seq | inDrop/CEL-Seq (Illumina Hi-Seq 2500) | Human |
| Segerstolpe | pancreatic islet | scRNA-seq | FACS/Smart-Seq2 | Human |
| Enge | pancreatic tissue | scRNA-seq | FACS/Smart-Seq2 | Human |
| Darmanis | middle temporal gyrus | scRNA-seq | Smart-Seq | Human |
| IP | temporal lobe cortex | Bulk RNA-seq | Illumina Next-Seq | Human |
| CA | middle temporal gyrus | snRNA-seq | Smart-Seq2 | Human |
| NG | prefrontal cortex | snRNA-seq | 10X Chromium | Human |
| MM | cerebral cortex | Bulk RNA-seq | Illumina Hi-Seq 2000 | Mouse |

An important aspect of our evaluations was to record how different methods perform with mismatched reference signatures, i.e., those derived from conditions or transcriptomic assays that do not perfectly match those of the target transcriptomic data. Additionally, we tested the impact of artificially “noised” versions of these reference signatures, which we simulated while respecting the general trend observed between these signatures and their corresponding true signatures (see Methods; also, we refer below to figures illustrating noisy signatures).

For our first evaluations, we used single cell RNA-seq data on human pancreatic islets from healthy subjects (14) to generate 100 pseudo-bulk profiles from weighted mixtures of cells of six pre-labeled types – alpha, beta, gamma, delta, acinar, and ductal. We adopted the procedures of Cobos et al. (1) for data processing and generation of mixtures. Deconvolution of this target data set (called “Segerstolpe-H”) was set to be performed using reference signatures constructed from the inDrop scRNA-seq data of pancreatic islet samples from Baron et al. (15). (Single cell transcriptomic profiles of cells of a type were averaged to obtain the reference signature of that cell type.) Note that the reference signatures and the target data set (pseudo-bulk profiles) are derived from different sequencing platforms (**Table 1**); this is one way of mimicking the technical differences between transcriptomics profiles that are often encountered in real-world deconvolution problems. **Additional File 1: Figure S1** shows the relationship between reference and true signatures, suggesting a high level of concordance despite the technical differences. Each of the four evaluated methods – BEDwARS, FARDEEP, CIBERSORT and CIBERSORTx – was used to infer cell type proportions in each pseudo-bulk profile and we calculated, for each cell type, the Pearson Correlation Coefficient (PCC) between true and predicted proportions across the 100 pseudo-bulk profiles. **Figure 2A** (group “Baron”) shows that BEDwARS makes more accurate estimates, indicated by average PCC over the six cell types, though all four methods proved highly accurate in this evaluation. **Figure 2H** shows this comparison for each cell type separately, for BEDwARS versus FARDEEP, revealing that the difference in performance is mainly for the ductal cell type (PCC 0.96 vs 0.88), as seen more clearly in **Figure 2G**. (See **Additional File 1: Figure S2** for similar comparisons for all cell types and methods.) The improved accuracy of BEDwARS estimates over the other three methods is also borne out when using alternative metrics of comparison -- mean absolute error (MAE) or root mean squared error (RMSE), rather than PCC – between true and estimated proportions (**Additional File 1: Figure S3A,B**).

**Figure 2.**
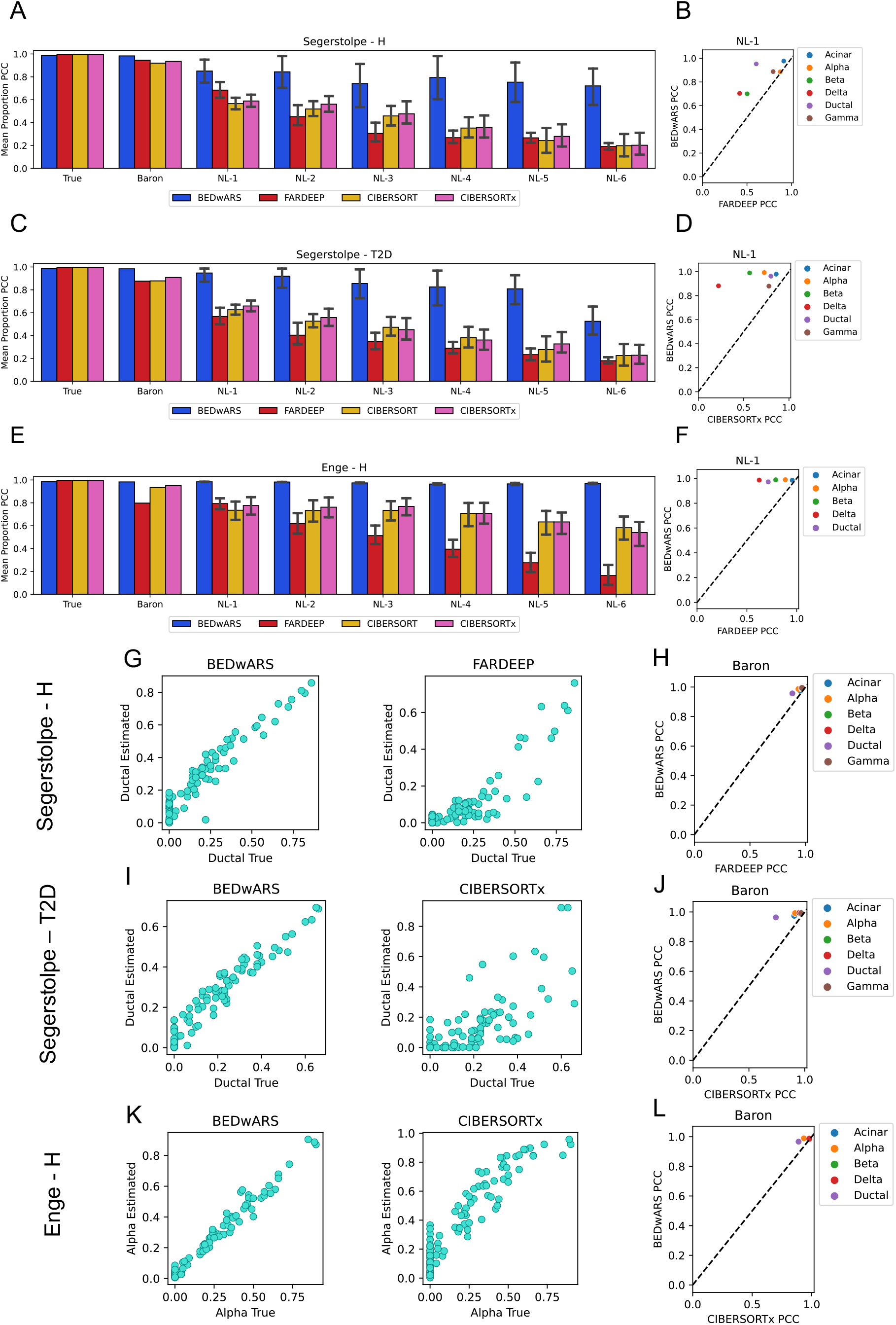
Evaluation of cell type proportion estimation in pancreatic transcriptomic profiles. (**A, C, E**) Pearson correlation coefficient (PCC) between true and estimated cell type proportions in 100 pseudo-bulk samples, averaged over cell types, is shown for different deconvolution methods. Results are shown for the Segerstolpe-H (**A**), Segerstolpe-T2D (**C**), and Enge-H (**E**) datasets. Category labels of bar charts indicate the reference signature, with “True” indicating the true underlying signature that is normally not available during deconvolution, “Baron” indicating the Baron signatures, and “NL-x” indicating Baron signatures with noise added at level x. For “NL-x”, results shown are mean with 95% confidence interval from evaluations using 11 variants of the Baron signature with noise added at level x. (**B**,**D**,**F**,**H**,**J**,**L)** PCC for each cell type separately is compared between the two best methods for respective datasets, when using NL-1 signatures (**B**,**D**,**F**) or Baron signatures (**H**,**J**,**L**). (**G,I,K**): Estimated and true proportions in the 100 pseudo-bulk profiles are directly compared, for a single cell type from each dataset, and for the two best methods for that dataset. BEDwARS performance is more robust to noise than the other methods in all datasets. All methods have comparable performance when the true signature is used. For the Baron signature, performance of BEDwARS is similar to other methods, with a noticeable improvement for Segerstolpe-T2D dataset. BEDwARS provides better estimates for all cell types in the NL-1 evaluations and for at least one cell type with the Baron signatures.

To test the effect of noisy reference signatures, we created six sets of perturbed versions of the above-mentioned Baron reference signature, representing different levels of noise and named “NL-1”, “NL-2”, etc., using randomization to create 11 such signatures in each set. Noise was introduced as described in Methods, maintaining the overall trend with the true signature and with a greater noise level inducing greater variance among genes with similar expression values in the true signature. The extent of injected noise is illustrated, for the lowest and highest levels NL-1 and NL-6, in **Additional File 1: Figure S1**; as seen in **Additional File 1: Figure S4A**, increasing noise levels result in progressively decreasing correlation coefficient between the reference and true signatures. As shown in **Figure 2A**, accuracy of cell type proportion estimation degrades with increasing noise levels (groups NL-1, … NL-6), but to a far lesser extent for BEDwARS than the other three methods. **Figure 2B** shows that the advantage of BEDwARS over FARDEEP, the next best method at the lowest noise level (NL-1), arises due to substantially greater accuracy for inferring proportions of delta, beta, and ductal cell types. **Additional File 1: Figure S5** shows BEDwARS has greater or equal accuracy compared to CIBERSORTx for every cell type and noise level.

Next, we created a new target data set (called “Segerstople-T2D”) comprising pseudo-bulk mixtures derived from the same study as in **Figure 2A** (human pancreatic islets (14)) but representing patients with Type II Diabetes (T2D). (See **Additional File 1: Figures S4B** and **S6**.) The trends noted above were even more pronounced now (**Figure 2C**) and the gap between BEDwARS and other methods was larger at all noise levels, including the lowest (NL-1), where cell types beta and delta were most poorly handled by the second-best method (CIBERSORTx, **Figure 2D**). (See **Additional File 1: Figure S7** for more complete comparisons.) In fact, the performance gap was substantial even in the absence of noise (**Figure 2C**, group “Baron”), with the greatest difference seen for the ductal cell type (**Figures 2J,I**). **Additional File 1: Figure S8** shows a detailed comparison with all methods for all cell types, revealing for instance that CIBERSORT and CIBERSORTx (CIBERSORT(x)) underestimate acinar cell type proportions by nearly an order of magnitude while BEDwARS estimates are close to the true values.)

We repeated the above evaluations with a different target data set – pseudo-bulk mixtures generated using scRNA-seq data from Enge et al. (16), representing human pancreatic tissue from healthy subjects (“Enge-H data set”). This is similar to the above tests in that the reference signatures (from Baron et al. (15)) and target profiles represent different technologies (**Table 1**). (Also see **Additional File 1: Figures S4C** and **S9**.) Performance comparisons yielded similar trends (**Figure 2E**) – BEDwARS showed marginal improvement over CIBERSORT(x) with the Baron signatures (**Figure 2L**), but now the gap with FARDEEP was larger (average PCC of 0.98 vs 0.8). Closer inspection showed that CIBERSORTx estimates, while highly correlated with true proportions (average PCC of 0.95), were often significantly inaccurate in absolute value (**Figure 2K**, **Additional File 1: Figure S10**); BEDwARS estimates were clearly more accurate in terms of MAE and RMSE (**Additional File 1: Figure S3E,F**). BEDwARS also showed a remarkable robustness to increasing noise levels, with progressively greater improvements over the other methods. A direct comparison with the second-best performing method (FARDEEP) at the lowest noise level (**Figure 2F)** shows that the largest performance gap is for beta, delta and ductal, the same cell types noted in **Figure 2B**. (Also see **Additional File 1: Figure S11** for comparisons to CIBERSORTx at varying noise levels.)

In summary, BEDwARS was found to provide more accurate estimates of cell type proportion compared to three leading methods, across a range of benchmarking conditions representing varying levels and sources of divergence between the true cell type signatures underlying the target data set and the provided reference signatures. (This was observed not only with the PCC but also alternative evaluation metrics such as MAE and RMSE.) Notably, all methods yielded near-perfect estimates of proportions when provided the true signatures, in all settings, indicating that the challenge in accurate deconvolution arises mainly from signature mismatch and noise.

### BEDwARS accurately estimates cell type signatures from noisy references

The principle underlying robust deconvolution by BEDwARS is to jointly estimate proportions as well as cell type signatures, allowing the latter to diverge from the reference. It is natural to ask, then, if the estimated signatures are indeed accurate. This can be assessed by comparing the BEDwARS-estimated cell type signature to the corresponding true signature from the target data set, using correlation coefficients. An alternative strategy to reconstructing the true signatures is to estimate cell type proportions in the target data set (as above) and use this information to re-estimate the true signatures. For this last step, we chose RODEO, a leading deconvolution method based on robust linear regression that infers cell type-specific signatures underlying a given bulk transcriptomics data set, given their cell type proportions in each sample. RODEO has been shown to be more accurate compared to other existing methods and to be robust to noise in the cell type proportions provided to it. We thus compared signature estimation accuracy of BEDwARS and RODEO, with the latter using cell type proportions estimated using FARDEEP, CIBERSORT or CIBERSORTx (in three separate runs). We also examined the accuracy of RODEO when provided cell type proportions from BEDwARS deconvolution. The correlation between the true signatures and reference signatures was used as a baseline.

The above evaluations were performed on each of the three benchmarks with human pancreatic data and revealed a few clear trends (**Figure 3**). First, BEDwARS estimates significantly more accurate signatures when provided with noisy reference signatures, with the accuracy gap increasing with noise levels. For instance, on the Segerstolpe-T2D target data set, even at the lowest noise level (NL-1) BEDwARS recovers the true underlying signatures with a correlation of 0.97 (on average), while the second-best alternative method (RODEO-CIBERSORTx) achieves an average correlation of 0.69 (**Figure 3C**). This gap is most pronounced for the ductal cell type (**Figure 3D**), a trend also seen in the other two benchmarks -- Segerstolpe-H and Enge-H (**Figures 3B,F**). In fact, the advantage of BEDwARS over RODEO-CIBERSORTx manifests to varying degrees for five of the six cell types, across all noise levels (**Additional File 1: Figures S12-S14**). A second trend we noted was that all methods inferred accurate signatures when provided with the Baron signatures without noise (**Figures 3A,C,E**, group Baron). Thirdly, we realized that poor signature estimation was tied mainly to errors in the proportion estimation (the first deconvolution step), as evidenced by the fact that RODEO recovers equally accurate signatures as BEDwARS if provided with the more accurate cell type proportions from BEDwARS; this is true regardless of the noise levels. Trends seen here were confirmed when using RMSE instead of correlation as the metric for signature comparison (**Additional File 1: Figure S15**).

**Figure 3.**
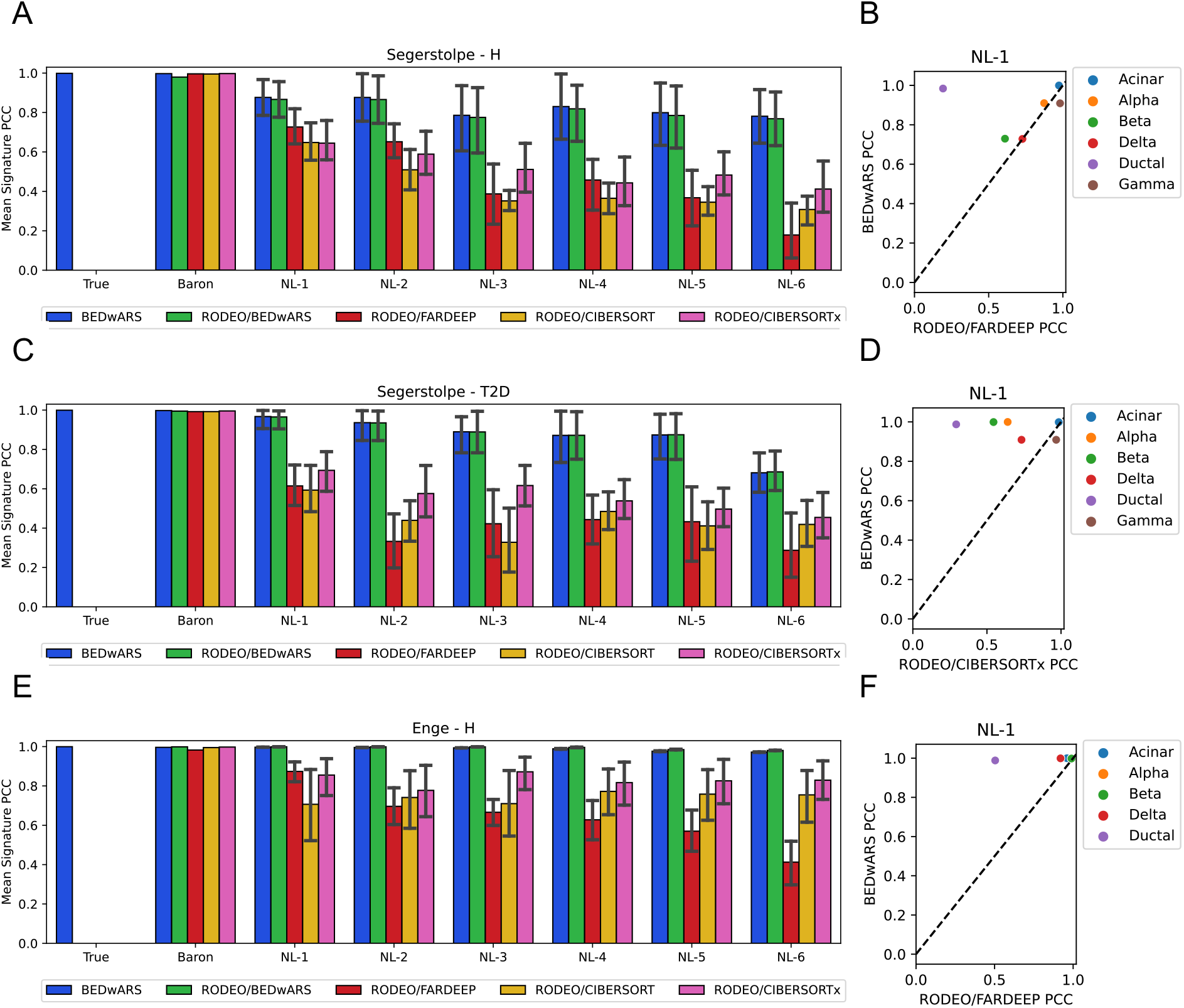
Evaluation of cell type signature estimation using pancreatic transcriptomic profiles. (**A**, **C**, **E**) Pearson correlation coefficient (PCC) between true and estimated cell type signatures, averaged over cell types, is shown for different deconvolution methods. Signatures were estimated by deconvolving 100 pseudo-bulk samples generated from Segerstolpe-H (**A**), Segerstolpe-T2D (**C**) and Enge (**E**) datasets. Performance of BEDwARS is compared with RODEO provided with cell type proportion estimates obtained using CIBERSORT, CIBERSORTx, FARDEEP or BEDwARS. Category labels of bar charts indicate the reference signature used. Category label “True” indicates that the true signatures were provided as reference to BEDwARS; no comparisons are made to RODEO in this case, rather this setting was used to assess if BEDwARS, which allows the estimated signature to deviate from the given reference, reports back an estimated signature similar to the true signature. BEDwARS is more robust to the increasing noise levels in recovering the true signatures in all datasets. RODEO provided with BEDwARS-estimated cell type proportions performs as well as BEDwARS, suggesting that accurate proportion prediction by BEDwARS is key to accurate signature estimation. (**B**, **D**, **F**) PCC for each cell type separately is compared between BEDwARS and its closest competitor (excluding RODEO/BEDwARS) for respective datasets, when using NL-1 signatures. In this setting, the PCC is substantially higher for BEDwARS-estimated signatures of ductal cell type than other methods, for all datasets.

The results of this and the previous section together demonstrate the value of jointly estimating cell type proportions and signatures when deconvolving bulk profiles using noisy or mismatched reference signatures. Indeed, when true signatures underlying the target data set are known accurately, all methods recover proportions accurately (**Figure 2A,C,E**; group True), and when proportions are estimated accurately using BEDwARS, RODEO can also provide equally accurate signatures.

### Robust deconvolution of brain transcriptomic profiles with BEDwARS

The next set of evaluations were performed following the recent benchmarking study of Sutton et al. (9), where scRNA-seq data from middle temporal gyrus in human brain of (17) were used to generate the target data set. Using their methodology, we generated 100 pseudo-bulk profiles as weighted mixtures of the three most frequent cell types in the scRNA-seq data – neurons, astrocytes and oligodendrocytes. As above, the proportions and true signatures used here were recorded as ground truths of the benchmark. In our first evaluations on these data, the reference signatures used were bulk RNA-seq profiles of <u>i</u>mmuno<u>p</u>urified (IP) cells from human brain (18) (see Methods); this is the “IP” signature. Noisy versions of this signature were also tested. (See **Additional File 1: Figures S4D** and **S16** for illustrations of how the IP signature and its noisy versions relate to the true signature.)

As shown in **Figure 4A**, BEDwARS provides more accurate estimates of cell type proportions in the brain data set, as compared to the three other methods. This is true even with the no-noise reference signatures (group ‘IP’, BEDwARS correlation 0.91 vs. CIBERSORTx correlation 0.84). BEDwARS deconvolution accuracy remains stable (between 0.91 and 0.85) at the wide range of noise levels, while other methods see their accuracy drop from ~0.83 (at no noise) to ~0.32 at the highest noise level; this trend is also seen with alternative metrics such as MAE and RMSE (**Additional File 1: Figure S17A,B**). A closer examination (**Figure 4E**) reveals that with the IP signature the oligodendrocytes cell type is the primary reason for performance deterioration in CIBERSORTx (BEDwARS correlation 0.94 vs. CIBERSORTx correlation 0.81), which underestimates the true proportions by nearly four-fold. (Also see **Additional File 1: Figures S18** and **S19**.) Evaluation of signature estimation accuracy by PCC criterion (**Figure 4B**) suggests that all methods are capable of recovering the true underlying signatures for this data set, although at higher noise levels RODEO using BEDwARS-estimated cell type proportions (RODEO-BEDwARS) clearly outperforms others, including BEDwARS. (Also see **Additional File 1: Figure S20**.) This is reaffirmed by evaluations with the RMSE criterion **(Additional File 1: Figure S15D**) and provides further support for the advantage of BEDwARS in terms of proportion estimation.

**Figure 4.**
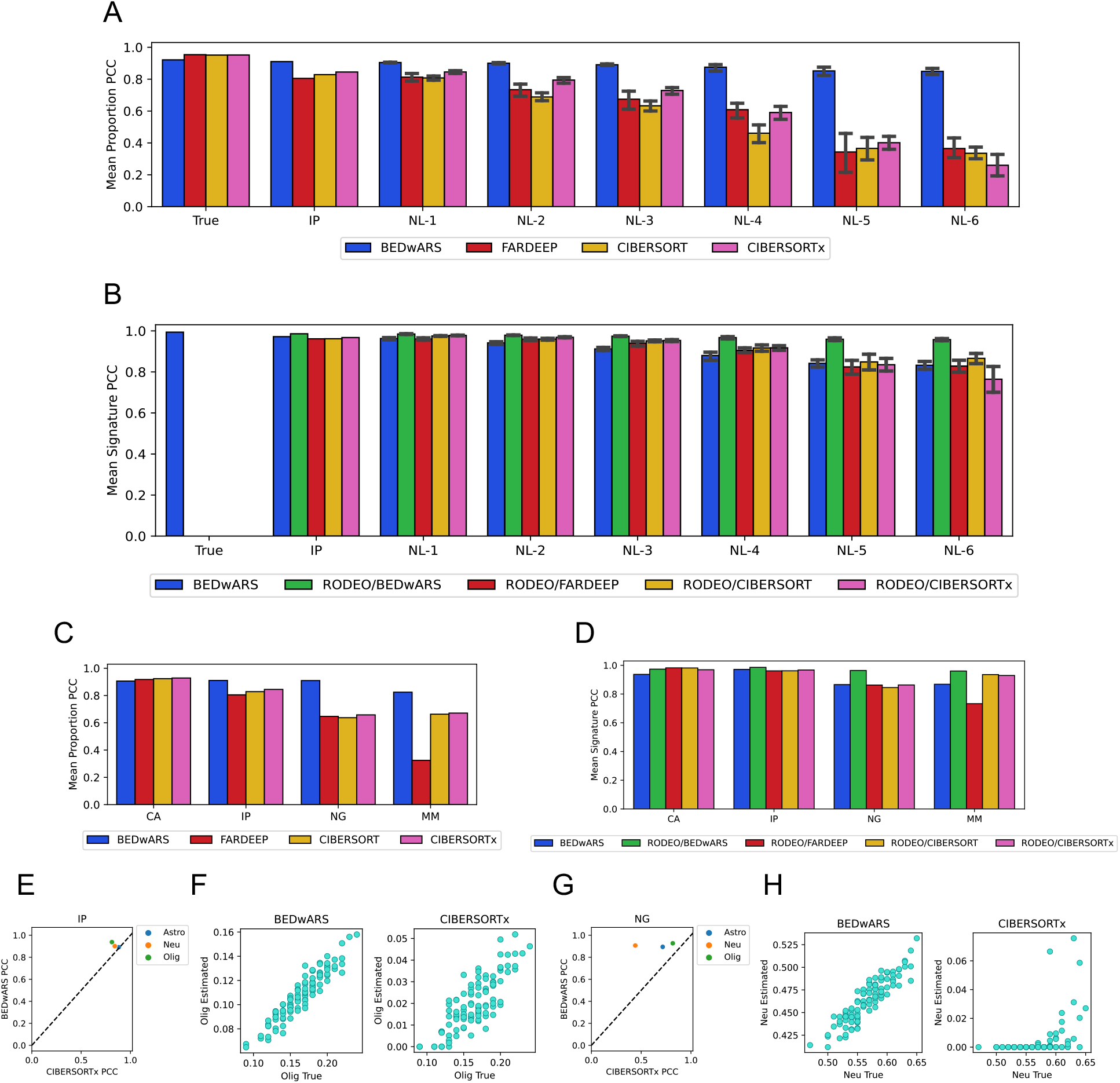
Evaluation of cell type proportion and signature estimation from brain transcriptomic profiles. (**A**, **B**) Pearson correlation coefficient (PCC) computed between the estimated and true cell type proportions (**A**) or cell type signatures (**B**), averaged over cell types, when deconvolving 100 pseudo-bulk samples generated from Darmanis dataset. Category labels of bar charts indicate the reference signature used. BEDwARS has higher PCC than the other methods in the estimation of cell type proportions for the IP signature and its noisy versions (NL-x), with the performance gap increasing as the noise level increases. For estimation of cell type signatures, RODEO provided with BEDwARS-estimated proportions (RODEO/BEDwARS) outperforms other methods including BEDwARS. (**C**, **D**) Average PCC between estimated and true cell type proportions (C) or signatures (D), using the IP signatures (same as in A,B) as well as the CA, NG, MM signatures. All methods perform comparably for proportion estimation when using the CA signature but BEDwARS exhibits better performance when the reference signature is more diverged from the true signature, such as NG (different region of human brain) and MM (mouse brain). All methods show comparable performance in signature estimation when provided the CA and IP references signatures, but RODEO provided with BEDwARS-estimated proportions exhibits superior performance for the more diverged reference signatures (NG and MM). (**E, G)** PCC for each cell type separately is compared between the two best methods (BEDwARS and CIBERSORTx) when using IP signatures (E) or NG signatures (G). In each case there is at least one cell type for which BEDwARS estimates have a higher correlation with true proportions – oligodendrocytes (Olig) in (E) and neurons (Neu) in (G). (**F, H**) Estimated and true proportions in the 100 pseudo-bulk profiles are directly compared, for a single cell type, for the two best methods when using the IP signatures (F) or NG signatures (H). In (F), BEDwARS estimates are not only better correlated (with true proportions) than CIBERSORTx estimates, they are also considerably more accurate in magnitude. In (G) BEDwARS estimates appear far better correlated with and also far more similar in magnitude to the true proportions than CIBERSORTx estimates. OLIG, ASTRO, and NEU stand for oligodendrocytes, astrocytes, and neurons respectively.

Next, we evaluated brain transcriptome deconvolution with three additional reference signatures that were considered by Sutton et al. (9). These include a signature obtained from bulk RNA-seq profiles of immune-purified cells from mouse brain tissue (19) (“MM”-Mus Musculus), one obtained from Human Cell Atlas containing single-nucleus RNA-seq of adult human middle temporal gyrus (20) (“CA”-Cell Atlas), and one signature from single-nucleus expression profiles of adult human prefrontal cortex in control samples (21) (“NG”-Nagy et al.). **Additional File 1: Figures S21** and **S22** show that the MM and NG signatures are more diverged from the true signatures (than are IP signatures), while the CA signatures are more similar to the true signatures. As shown in **Figure 4C**, with the NG and MM reference signatures (more mismatch), the BEDwARS-estimated proportions (average correlation of 0.91 and 0.82 with true proportions) are clearly more accurate than CIBERSORTx-estimated proportions (correlation of 0.65 and 0.67) and FARDEEP (correlation of 0.65 and 0.32). The improvement over CIBERSORTx is most pronounced for the neuron cell type (**Figure 4F**) and oligodendrocytes (**Additional File 1: Figure S23**). Evaluations with MAE and RMSE metrics **(Additional File 1: Figure S17C,D**) confirm the advantage of BEDwARS in this benchmark, but only for the NG signature; the MM signature yields comparable accuracy across methods by these alternative metrics.

On the other hand, evaluation of signature estimation (**Figure 4D**) suggests that for the CA and IP signatures (more matched with true signatures) all methods perform equally well, while for the NG and MM signatures (more mismatched), RODEO using BEDwARS-estimated proportions has the best performance. (Also see **Additional File 1: Figure S15E** and **Figure S24B**.) For deconvolution with MM signatures, we observed (**Additional File 1: Figure S25**) that oligodendrocyte proportions are poorly estimated by all methods but BEDwARS estimates are well correlated with true proportions. This suggests that a cell type signature estimation method such as RODEO can benefit from proportion estimates that are accurate in relative if not absolute terms. Evaluations using the CA reference signature revealed similar performance by all evaluated methods, both for proportion estimation (**Additional File 1: Figure S26)** and for signature estimation, with correlation values of ~0.9 or greater (**Figures 4C,D**), highlighting the importance of matched reference signatures for the deconvolution task.

We also compared BEDwARS with the recently reported tool SCADIE (22), which, like RODEO, relies on initial estimates of cell type proportions provided by any existing deconvolution method and iteratively refines the cell type signatures and proportions. Our evaluations demonstrate that the performance of SCADIE in both proportion and signature estimation is strongly dependent on the quality of cell type proportions used in the initialization. In particular, we found that when using reference signatures that are more diverged from true signatures SCADIE initialized with CIBERSORTx- or FARDEEP-estimated proportions performs significantly worse than when initialized with BEDwARS-estimated and comparably to BEDwARS (for proportion estimation) and RODEO/BEDwARS (for signature estimation). (See **Additional File 1: Figures S27-S29**).

In summary, extensive comparative evaluations on a published set of benchmarks involving brain transcriptomics data reaffirmed the conclusions drawn from pancreatic islet benchmarks, that BEDwARS is capable of robust proportion estimation in the face of noisy and mismatched signatures and such proportions can then be the basis of more accurate signature estimation as well.

### Application of BEDwARS to characterize the cell type-specific regulomes of DPD-deficient patients

In this section, we present a case study in the use of single cell and bulk transcriptomics to characterize molecular mechanisms underlying a rare disorder. Dihydropyridine dehydrogenase (DPD) deficiency is caused by deleterious germline variants within the *DPYD* gene and typically presents as a pharmacogenomic condition, in which patients are at significantly higher risk of severe adverse events when treated with the commonly used chemotherapeutic 5-fluorouracil (5-FU) (23). DPD deficiency has also been linked to rare inborn error of metabolism that is accompanied by neurological disorders of varying degrees of severity in children (24,25). The penetrance of the pediatric condition within individuals with DPD deficiency is very low. For the purposes of this manuscript, we will refer to this condition as “pediatric DPD deficiency” to distinguish it from the pharmacogenomic disorder or the generalized reduction in DPD function. While the biochemistry surrounding DPD is well characterized, there is extremely limited information pertaining to how DPD deficiency could contribute to the clinical presentation of neurologic and metabolic conditions in affected children.

The analyses presented in this manuscript represent a subset of a larger clinical study designed to characterize the developmental and biochemical pathways that are altered in pediatric DPD deficiency with the goals of gaining a better understanding of the disease etiology as well as identifying potential therapeutic approaches to improve quality of life for affected patients. For the overall study, fibroblasts were obtained from affected individuals, non-affected family members, and unrelated controls. Fibroblasts were reprogrammed into induced pluripotent stem cells (iPS cells), which were subsequently used to derive neural organoids. At least 3 independent iPS clones were generated from each subject.

For the present study, RNA-seq was performed on 72 brain organoids from three patients with pediatric DPD deficiency (referred to as DPD1, DPD3, and DPD6) and on 48 organoids from two non-affected subjects (DPD2 and DPD4). ScRNA-seq profiling was also performed for three organoids from patient DPD1 and for three organoids from the non-affected subject DPD4. For purposes of cross-technology calibration, we ensured that eight of the organoids profiled using bulk RNA-seq in each group (patient or non-affected) were generated and cultured in parallel with the three organoids used for scRNA-seq. We will refer to these bulk-profiled organoids as “semi-matched” bulk samples below.

To deconvolve the bulk RNA-seq profiles into cell type-specific components, we first generated reference signatures using the single-cell data. Single-cell profiles from the affected and non-affected subjects were processed together to obtain ~29,707 cells that segregated into 17 clusters that potentially represented different cell types and states (**Figure 5A**). To identify cell types represented by these clusters, we utilized a multi-pronged strategy based on work by Tanaka et al. (26). The average expression of neuronal markers (STMN2, GAP43, DCX) and early neurogenesis genes (VIM, HES1, SOX2) was used to discriminate neuronal from non-neuronal clusters (**Figure 5B**, **Additional File 1: Figure S30**). Further resolution was achieved through the consideration of additional known cell type markers, as well as statistically identified marker genes, enrichment of cell type-related Gene Ontology terms in these markers and overlaps with similarly obtained marker sets from Tanaka et al. (see Methods) (**Additional File 1: Figure S31**). Using this approach, we were able to assign cell types to 15 of the 17 clusters (**Additional File 1: Table S1**). Notably, cortical neurons and astrocytes were the only cell types with representation in the affected and non-affected samples. Reference signatures were then obtained as average gene expression profile of each cell type found in the non-affected individual (astrocytes (AS), cortical neurons (CN), progenitor cells (PGC), cilia-bearing cells (CBC), intermediate (INTER), BMP-related cells (BRC)), as well as three cell types in the affected individual (neurons (NEU), neuroepithelial cells (NEC) and cluster-11); in some cases, multiple clusters were mapped to the same cell type in this step (see Methods).

**Figure 5.**
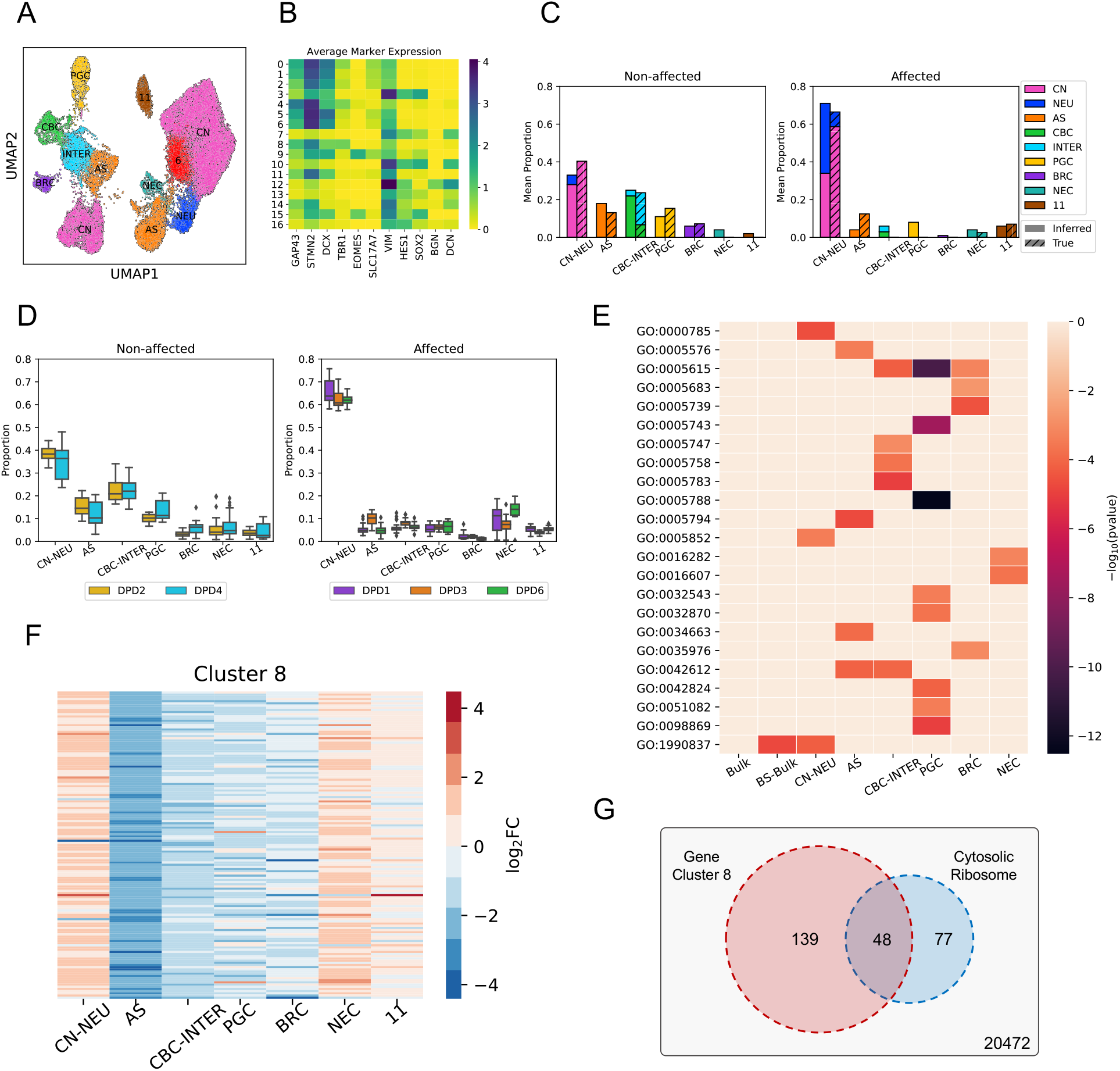
Cell type-specific characterization of transcriptomic differences between organoids from DPD deficiency affected and non-affected subjects using BEDwARS deconvolution of bulk RNA-seq data. **A**. UMAP plot of processed cells clustered into 17 groups. Cells on the right represent the affected patient and cells on the left represented the non-affected subject. **B**. Average expression of 11 marker genes in cells of each cluster indexed by numbers. These markers were used for cell type assignment. **C**. Comparison between the average inferred proportions (plain) from eight bulk samples of an affected and a non-affected subject and the average of “true” proportions (diagonal striped) derived from the semi-matched single cell data on three organoids for the same subject. **D**. Inferred proportions of different cell types obtained by BEDwARS deconvolution of bulk RNA-seq data from organoids derived from two non-affected subjects and three affected patients. CN-NEU: sum of inferred proportions of CN and NEU; CBC-INTER: sum of inferred proportions of CBC and INTER. **E**. Negative logarithm (base 10) of the p-value (−log_10_(pvalue)) of Hypergeometric tests of Gene Ontology (GO) term enrichments in the top 200 differentially expressed genes (DEGs) from bulk samples (72 affected vs 48 non-affected, “Bulk”), bootstrapped bulk profiles derived from the single cell data (100 affected vs 100 non-affected, “BS-Bulk”) and the cell type-specific bulk expression derived from BEDwARS deconvolution (72 affected vs 48 non-affected, “CN-NEU”, “AS”, “CBC-INTER”, “PGC”, “BRC”, “NEC”). None of the GO terms enriched in the top 200 DEGs of cell type-specific profiles are enriched in the top 200 DEGs derived from the bulk expression profiles. See **Additional File 1: Table S3** for information on GO terms. **F**. Logarithm (base 2) of the fold-change (log_2_FC) of expression of the 139 genes grouped into cluster 8 based on their pattern of differential expression in different cell types. **G**. The 139 genes of cluster 8 are highly enriched in the GO term “cytosolic ribosome” (Hypergeometric test p-value 2 × 10^−85^).

We next performed deconvolution of bulk RNA-seq profiles in each group (i.e., affected and non-affected) separately, using BEDwARS with the above-mentioned references signatures of nine cell types. (Bulk profiles for each group were first batch corrected to match the pseudo-bulk profiles generated from single cell data for the respective group, see Methods and **Additional File 1: Figure S32**.) As an internal control, we first compared estimated cell type proportions in the eight semi-matched bulk samples in each group to those in the scRNA-seq samples of the same individual and found the deconvolution to successfully recover the proportions of dominant cell types (**Figure 5C**). In both groups, the sum of inferred proportions of cortical neurons (CN) and neurons (NEU) matched the corresponding sum in the single cell data. However, the proportions of these two individual cell types could not be accurately resolved due to the similarity in the signatures (see **Additional File 1: Figure S33**). Similar observations were made for the cilia-bearing cells (CBC) and intermediate (INTER) cell type proportions, with their sum matching between bulk-deconvolved and single cell data. Apart from these four cell types, any other cell type with either the inferred or true proportion above 5% (AS, PGC, BRC in non-affected and PGC, cluster-11 in affected) was deconvolved accurately. The only exception to this trend was the AS cell type in the affected samples, where the true proportion from single cell data, roughly 10%, was underestimated at ~4%, at the expense of an over-prediction of PGC proportion. Overall, this exercise confirmed our ability to deconvolve cell type proportions in bulk RNA-seq from organoids, leading us to apply the same procedure to the entire data set.

We next deconvolved the 48 and 72 bulk profiles from two non-affected and three affected subjects respectively, using the same signatures and procedure as above. Estimated proportions of almost every cell type (and in two cases, sums over cell type pairs – CN+NEU, CBC+INTER) were consistent among subjects within the same group (**Figure 5D**). The CN+NEU proportion is noticeably higher in the affected subjects, consistent with the limited single cell data (**Figure 5C**), while CBC+INTER proportion is higher in the non-affected subjects. Ciliated cells (i.e., CBCs) are involved in extracellular signal transduction that is critical for patterning and morphogenesis during neural development (27,28). Disruptions to the processes facilitated by ciliated cells has been shown to contribute to both neurodevelopmental and degenerative diseases (29). The reduced number of CBC populations within organoids derived from affected individuals is suggestive that ciliopathies might be linked to DPD deficiency and contribute to the observed clinical presentation of pediatric DPD deficiency, warranting further study of this novel observation.

Deconvolution with BEDwARS also allowed us to examine cell type-resolved components of each bulk transcriptomic profile, obtained by multiplying the cell type’s inferred proportion with the respective inferred signature. We could thus compare gene expression between the two groups of samples (affected versus non-affected) in a cell type-specific manner, with the large numbers of bulk RNA-seq samples (72 and 48 in the two groups) providing high statistical power and the multiplicity of subjects in each group offering a more diverse representation than possible with the limited single cell data. We derived the genes most differentially expressed (DE) between groups, for each cell type separately (**Additional File 2: Table S2**), and performed Gene Ontology (GO) enrichment tests for the top 200 genes to characterize their biological functions (**Figure 5E, Additional File 1: Table S3** and **Additional File 3: Table S4**), see Methods. None of the significant GO terms (FDR < 0.05), except for one, obtained from this cell type-specific analysis were significantly enriched in the top 200 DE genes derived from bulk profiles or from bootstrapped samples of the single cell data. This demonstrates that the deconvolution approach likely helped reveal latent patterns of gene expression changes within specific cell populations that could not be observed within bulk data or in a limited sampling strategy (i.e., small number of organoids from fewer subjects) common to scRNA-seq analyses.

For example, chronic fatigue and metabolic dysfunction, consistent with mitochondrial disorders, have been previously reported in subjects with DPD deficiency (30). However, it is unclear if mitochondrial disorder is a shared feature of pediatric DPD deficiency (31). Mitochondria-related GO terms were identified in the deconvolved expression data for PGC, CBC-INTER, and BRC cell types (**Figure 5E and Additional File 1: Table S3** and **Additional File 3: Table S4**, suggesting that changes in the expression of mitochondrial genes within these compartments might be relevant to the disease etiology. As another example of latent features potentially identified by this analysis, dysfunctions in protein synthesis and folding (e.g., endoplasmic reticulum, ER, dysfunction) have been suggested to contribute to numerous neurological disorders; however, the etiology and/or pathogenesis have not been fully elucidated (32). Terms related to ER and translation initiation were significantly enriched in CBC-INTER, PGC, CN-NEU, NEC, and AS clusters (**Figure 5E**), suggesting that changes in protein translation and folding might contribute to clinical presentation of pediatric DPD deficiency.

In a complementary analysis, we clustered genes based on their patterns of differential expression across all nine cell types (**Additional File 1: Figure S34**), obtaining 10 major clusters of 39-5107 genes (**Additional File 4: Table S5**), with similar GO term associations as above (**Additional File 5: Table S6**). One of these clusters (cluster 8, **Figure 5F**) shows a pattern in which genes are up-regulated in CN+NEU and NEC but downregulated in AS, PGC, and CBC+INTER cell types, in affected subjects compared to non-affected subjects. Genes showing this pattern of expression were enriched for those associated with the cytosolic ribosome (i.e., “free” ribosomes, **Figure 5G**; FDR=5×10^−83^). Cells of the nervous system, in particular neuronal cells, rely on localized translation of gene products via cytosolic ribosomes (33). This pseudo-compartmentalized translation within neural cells has been shown to create spatial variability in protein expression that is critical for neural development, function, and plasticity (34). Disruptions to localized translation have been linked to various neurodevelopmental and neurodegenerative disorders (35,36). Combined with the results linking ER/translation GO terms in these same cells, these findings indicate that dysregulation of translation and protein folding, whether at the ER and/or at distal sites, might contribute to the clinical presentation of pediatric DPD deficiency.

## Discussion

We present here a Bayesian approach to the deconvolution of bulk expression profiles especially designed to address potential differences between reference cell type signatures and the true (but unknown) cell type signatures underlying the bulk profiles, a common challenge in deconvolution. The novelty of our method lies in adjusting for such unknown differences by simultaneously estimating the true signatures (as perturbed versions of reference signatures) and optimizing for the cell type proportions in each bulk profile. Through extensive benchmarking utilizing eight different datasets and a procedure for generating noisy reference signatures in a controlled manner, we demonstrate that not only does our method outperform leading in-class methods in the estimation of cell type proportions, but it also is more robust to the noise added to the reference signatures. Furthermore, with a few exceptions our method achieves a better estimation of true cell type signatures than the state-of-the-art method. Given the well-recorded challenges arising from signature differences between bulk datasets and corresponding reference signatures, we believe BEDwARS robustness to noise, demonstrated on semi-synthetic data, will confer on it a significant advantage in real world deconvolution tasks.

One might expect that a Bayesian deconvolution approach that allows for noisy signatures as part of its model will not perform best if the reference signatures are in fact very similar to the true signatures underlying the bulk data, i.e., when the anticipated noise is not there. However, we noted that our model’s performance is better than or competitive with the other approaches in most of the benchmarks in this study, including those where the signature differences were the smallest. This suggests that in such cases the model learns to perturb the reference signature to a lesser extent based on the data.

The BEDwARS model estimates the true cell type signatures using reference signatures as prior information. This means that *G* × *C* parameters are learnt from the entire bulk data, where *G* and *C* are the numbers of genes and cell types respectively. Among other things, this implies that one needs to exercise care when using BEDwARS with many cell types and few bulk samples. While it is difficult to make precise recommendations about these numbers, since they depend on the additional data characteristics, we note that our evaluations have been successful with ~40 bulk profiles and ~10 cell types. Scenarios with more bulk profiles and fewer cell types should be safe for BEDwARS application, and further tests are needed for other scenarios. These considerations further imply that cell type proportion estimation cannot be done for one bulk profile at a time and all bulk samples should be provided at once to BEDwARS, even though the estimated proportions are different for each sample.

Applying BEDwARS to a new dataset, we were able to gain new insight into a rare pediatric inborn error of metabolism linked to DPD deficiency. Using a limited set of scRNAseq data to generate reference signatures, we deconvolved bulk RNAseq data from complex patient-derived neural organoids to identify novel expression changes in specific neural cell types. These findings suggest that multiple changes likely contribute to the pathology of the disorder, including disruptions to ciliated cell function, mitochondrial dysfunction, and alterations to translational machinery at the ER and associated with free ribosomes. While pediatric DPD deficiency has previously been suspected of having a mitochondrial component (31), to our knowledge this is the first reported evidence for possibly involvement of ciliopathy and impaired translational control in the etiology of the disorder. Further study of these pathways as potential targets for the development of new treatments for ameliorate the symptoms associated with pediatric DPD deficiency is warranted.

Based on our empirical experience as well as on theoretical grounds, we believe BEDwARS may not be able to accurately tease apart the contributions/proportions of highly correlated cell types (4,37), a common problem with deconvolution methods. We also believe that BEDwARS performance can be further improved by modifying it to be more robust to outliers, e.g., by changing the optimization objective or through outlier detection and removal. We leave these important engineering challenges for future iterations of the tool.

## Methods

### Preprocessing of datasets used for benchmarking

#### Pancreas data

“Baron”: scRNA-seq data obtained from Baron et al. (15) were used in generating signatures. Count-level data on cells from all four human subjects (3 healthy and 1 T2D) were utilized. “Segerstolpe”: scRNA-seq datasets from Segerstolpe et al. (14) were used, with count-level data on cells from six healthy individuals forming the “Segerstolpe-H” dataset and those from four T2D individuals forming the “Segerstolpe-T2D” dataset. Cells with “not applicable”, “unclassified”, and “co-expression” tags were removed in this step. “Enge”: Count-level scRNA-seq data on cells from eight healthy individuals, reported in (16), formed the “Enge-H” dataset. Except for the Enge-H dataset, six cell types – alpha, beta, gamma, delta, acinar and ductal – were analyzed. The Enge-H dataset did not contain gamma cell type therefore only the remaining 5 cell types were considered.

Following the quality control procedure of Cobos et al. (1), for each pancreatic dataset, we removed cells with library size, ribosomal or mitochondrial content more than three median absolute deviations away from the median. Then, only the genes with nonzero counts in at least 5% of all cells were kept. Finally, RPKM-normalization was done using hg19 human genome assembly.

#### Brain data

scRNA-seq data from the middle temporal gyrus in human brain, reported by Darmanis et al. (17), were RPKM-normalized using hg19 human genome assembly, following the preprocessing pipeline of Sutton et al. (9), to form the “Darmanis” dataset. The three most abundant cell types – neurons, astrocytes and oligodendrocytes – were analyzed. “IP” signatures were obtained from FPKM-normalized RNAseq data on immunopurified cells from human adult brain (temporal lobe cortex) (18). “MM” signatures were formed from FPMK-normalized RNA-seq data on immunopurified mouse brain tissue, reported by Zhang et al. (19). “CA” signatures represent RPKM-normalized count-level single-nucleus expression in middle temporal gyrus (20), while “NG” signatures were single-nucleus expression of human prefrontal cortex (21), obtained from Sutton et al. (9) without repeating their preprocessing pipeline.

### Generation of cell type signatures and their variants for benchmarking

#### Signature generation

Cell type signatures were generated by averaging the gene expression profiles of all cells of the same type in a dataset. Following Sutton et al. (9), for the signatures generated from Darmanis, IP, MM, and CA datasets, only genes with more than one RPKM/FPKM expression in at least one cell type were retained. NG signature was taken directly from Sutton et al. (9)Signatures used in brain gene expression deconvolution were restricted to neurons, astrocytes, and oligodendrocytes, matching a similar restriction imposed on the Darmanis data set (see above).

#### Perturbation of signatures

This procedure is performed separately for each cell type, starting with a reference signature and a true signature of that cell type. (The true signature represents the target dataset to be deconvolved and the reference signature reflects the related dataset used by the deconvolution method.) The procedure, described next, modifies the reference signature by adding random noise to it while maintaining a statistical relationship between the reference and true signatures. It is parameterized by a single parameter *σ*. First, the signatures are log-transformed. (Genes with zero expression in any cell type were excluded from the reference and true signatures before the transformation.) Next, genes are partitioned into equal-frequency bins based on their expression values in the true signature. Since a reference signature generally exhibits high positive correlation with the true signature (e.g., see **Additional File 1: Figures S1, S6, S9**, and **S16**, genes in a bin that represents high (or low) expression level in the true signature have a high (resp., low) mean expression in the reference signature as well. This is the statistical relationship that the perturbation procedure maintains, as noted next. In the next step, for each bin, the mean expression of genes in that bin is calculated and each gene’s deviation from the mean is scaled by the same constant; this constant is set so that the resulting expression values (of genes in that bin) have a standard deviation of *σ*. This step ensures that the average expression of genes in a bin remains unchanged, so the above-mentioned statistical relationship is maintained; at the same time the variance of reference gene expression in each bin is increased to the pre-set level *σ*, thereby adding noise to the signature overall. Some examples of the result of such perturbation are shown in **Additional File 1: Figures S1, S6, S9** and **S16**.

In our benchmarking, we set *σ* to values in the range [1, 2.25] with increments of 0.25, to define six noise levels called “NL-1” (*σ* = 1), “NL-2”, … “NL-6” (*σ* = 2.25), with higher noise-levels resulting in lower correlation coefficients between reference and true signatures. The partitioning of genes was done so that each bin has 300 genes when benchmarking the Darmanis dataset with IP signatures and 100 genes when adding noise to the Baron signatures for deconvolution of Segerstolpe-H, Segerstolpe-T2D, and Enge-H datasets. The exception to this was in the benchmarks where the Baron signatures were used with the Segerstolpe-H and Segerstolpe-T2D target datasets, no perturbation was applied to the signatures of acinar and ductal cell types, as the reference signatures were already relatively poorly correlated with the true signatures for these cell types. A similar exception was made for the ductal cell type when adding noise to the Baron signatures for use with the Enge-H dataset

The above deterministic procedure for signature perturbation was followed by a second procedure that introduces additional noise to the reference signatures. All genes in the same bin (defined above, representing a small range of values of true signature) were further partitioned into bins of four genes each based on their reference expression values; then the reference expression levels of the four genes in each such bin were shuffled. The entire procedure was repeated 10 times to get 10 variants of the deterministically perturbed reference signature from the first procedure (previous paragraph). Thus, for each noise level, we obtained 11 different randomly generated variants of the reference signature, perturbing it similarity to the true signature in a controlled manner. Note: The IP signature (Zhang et al. (18)) and its noisy variants were restricted to include genes with at least two-fold higher expression in one cell type compared to the others.

### Generation of pseudo-bulk mixtures

#### Pancreas datasets

We followed the pseudo-bulk mixture generation pipeline by Cobos et al. (1) with minor modifications. First, we randomly selected the number of cell types to be present in a mixture, uniformly from the range [2, K] where K is the total number of cell types. Second, the selected number of cell type identities were randomly sampled without replacement. Next, the “true” proportions associated with the selected cell types were uniformly sampled from [0.05, 1], followed by scaling to ensure that they sum to one. Finally, 100 cells were sampled so that each cell type was represented with its respective proportion and the expression profiles of the sampled cells were averaged to create a pseudo-bulk profile. By repetitions of this process, 100 mixtures with known cell type proportions and a pseudo-bulk expression profile were generated for each of the data sets Segerstolpe-H, Segerstolpe-T2D and Enge-H.

#### Brain datasets

One hundred mixtures and corresponding pseudo-bulk profiles were generated by sampling (without replacement) 100 cells at a time from the Darmanis dataset (17) and averaging their expression profiles. The same process was used by Sutton et al. (9) to generate the bulk mixtures for this dataset.

### Induced pluripotent stem cells (iPSCs) and cerebral organoids for the study of DPD-deficiency

iPSCs were reprogrammed from skin fibroblasts that were obtained from skin biopsies. Biopsies were collected following written informed consent/assent from the donor and/or guardian and approved by the Mayo Clinic Institutional Review Board (IRB protocol 14-005685). iPSCs were maintained on 60-mm plates coated with hESC-qualified Matrigel (Corning Life Sciences, Corning, NY) in mTeSR Plus medium (StemCell Technologies, Vancouver, Canada) containing 100 units/mL penicillin and 100 mg/mL streptomycin. Cells were grown at 37°C in humidified air containing 5% CO_2_. Differentiated cells were removed and medium was exchanged every 1–2 days. Cells were passed using ReLeSR (StemCell Technologies).

Cerebral organoids were generated from iPSCs using the StemCell Technologies STEMdiff Cerebral Organoid kit according to the manufacturer’s instructions. For bulk RNAseq analyses, organoids were harvested on day 46, lysed in TRIzol (Invitrogen, Waltham, MA), and stored at – 80°C until RNA extraction. RNA was extracted using the Zymo Research Direct-zol RNA miniprep kit (Zymo Research, Irvine, CA) according to the manufacturer’s instructions. RNAseq libraries were prepared using TruSeq Stranded mRNA reagents (Illumina, San Diego, CA). For scRNAseq, single cells were isolated using the Neural Tissue Dissociation Kit P (Miltenyi Biotec, Gaithersburg, MD) with gentle trituration. Single cell partitioning and scRNAseq library preparation performed using Single Cell Gene Expression reagents on a Chromium Controller (10x Genomics, Pleasanton, CA) in the Mayo Clinic Medical Genome Facility Genome Analysis Core. RNAseq and scRNAseq libraries were sequenced using 2×150 PE chemistry on a NovaSeq 6000 (Illumina) at the University of Minnesota Genomics Center.

For RNAseq, adapter sequences were removed using the TrimGalore wrapper around Cutadapt (38), and reads were aligned to the human genome (hg19) using two-pass mapping and genes expression quantified as gene counts using STAR (39). scRNAseq data was processed, mapped to hg19, and quantified using the CellRanger pipeline version 6.1.2 implemented on the 10x Genomics cloud analysis platform.

### Preprocessing of the scRNA-seq and bulk RNA-seq data for the study of DPD-deficiency

Quality control, clustering, and marker detection: Scanpy (40) was used to process the combined scRNA-seq data of the non-affected (DPD4) and affected (DPD1) individuals. In the quality control step, cells with less than 1000 genes expressed and genes that were detected in less than 500 cells were removed. Furthermore, cells with more than 5% mitochondrial gene percentages were removed. To cluster the cells, the top 2000 highly variables genes were selected based on the highest standardized variance approach of Stuart et al. (41) implemented as “seurat_v3” in Scanpy. Principal component analysis (PCA) was performed on these highly variable genes and the top 20 PCs were used to build the neighborhood graph of cells. Leiden graph clustering method with resolution 0.8 was used to detect 17 clusters of cells. Most cells for clusters (0,1,2,3,5,6,9,11,15) were from the affected individual (“affected clusters”) and the rest of clusters mostly contained non-affected cells (“non-affected clusters”). Markers for each cluster were identified using t-test and Benjamini-Hochberg method was used for multiple hypothesis testing correction. The markers were then filtered by their adjusted t-test p-value less than 0.05 and logFC greater than 0.25.

Cell type assignment details: A cell type was assigned to a cluster if at least two (out of three) criteria were met. The first criterion is based on the average expression of marker genes of the cell type, following Tanaka et al. (26), that had detectable expression in our dataset. Clusters with high average expression of GAP43, STMN2, and DCX were tagged as neuronal clusters. Among these clusters the expression of either TBR1 or SLC17A7 is indicative of cortical neurons (CN) whereas expression of EOMES is indicative of neurons (NEU). Based on average expression of neuronal marker genes, clusters 0, 1, 2, 4, 5, 6, 8 and 9 had supporting evidence of being cortical neurons and neurons, respectively. Non-neuronal clusters were identified by the high expression of VIM, HES1 and SOX2. Expression of two other markers, BGN and DCN, was detected for cluster 12, supporting its assignment to the progenitor cells (PGC) cell type. The cellular level expression of neuronal and non-neuronal marker genes is visualized in **Additional File 1: Figures S35** and **S30**.

The second criterion was the enrichment of certain Gene Ontology (GO) terms in the computationally derived markers of each cluster following the pipeline of Tanaka et al. (26). Top 200 markers of each cluster (**Additional File 6: Table S7**) were tested for their enrichment in specific GO terms using the KnowEnG platform (42). Clusters 3 and 10 were enriched in astrocyte differentiation, cluster 15 in mitosis-related terms, cluster 13 in motile cilium and epithelial cilium movement, and cluster 16 was also enriched in cilium-related terms (see **Additional File 1: Table S8** and **Additional File 7: Table S9**). Following Figure S1.B of Tanaka et al. (26), we interpreted enrichment in astrocyte differentiation (clusters 3,10), mitosis-related terms (cluster 15) and cilium-related terms (clusters 13, 16) as indicators of astrocytes, neuroepithelial cells (NEC), and cilia-bearing cells (CBC), respectively. (See **Additional File 1: Table S1**.)

The third criterion used in cell type assignment was based on the overlap of the top 100 computationally derived markers of a cluster (**Additional File 6: Table S7**) with the corresponding markers from an annotated cluster in Tanaka et al. (26). The cell type annotation of the annotated cluster of (26) with the largest overlap is used for labeling our clusters. In cases where multiple annotated clusters of (26) were assigned subtypes of the same cell type, the overlap was averaged over all subtypes. Based on this criterion, clusters 0,1,2,4,5, and 8 should be designated as cortical neurons, cluster 9 as neurons, clusters 3 and10 as astrocytes, clusters 13 and 16 as cilia-bearing cells (CBC), clusters 6,7,11 and 14 as intermediate (INTER), and cluster 15 should be tagged as neuroepithelial cells (NEC).

The final cell type assignment was based on presence of at least two of the above three types of supporting evidence. Cluster 6 had conflicting evidence in support of cortical neurons (CN) and intermediate cells (INTER) based on the first and third criteria respectively. Cluster 11 was only supported by the third criterion to be assigned to intermediate cells (INTER). Therefore, clusters 6 and 11 were left unassigned and their indices were used as their “cell types”. Clusters 7 and 14 were identified as non-neuronal clusters and were supported by the overlap criterion only. However, we noted that they are well separated in the UMAP plot, and both can be assigned to BRC or INTER. Through a closer examination of **Additional File 1: Figure S31** we decided to tag cluster 7 and 14 with INTER and BRC cell types. The rest of the clusters were supported by two out of three criteria and were thus reliably annotated with cell types.

#### Cell type Signature Generation

The preprocessed combined scRNA-seq data from non-affected and affected individuals was further filtered for cells with library size, ribosomal or mitochondrial content more than three median absolute deviations away from median. Also, only the genes with nonzero counts in at least 5% of all cells were kept. The signature was generated from all non-affected annotated clusters as well as three affected clusters. So, the final signature contained cell types CN, AS, CBC, INTER, BRC and PGC from the non-affected individual as well as NEC, Neuron, and cluster 11 from the affected individuals. The difference between the number of cell types used in the reference signature (9) and the total number of clusters (17) was due to the existence of multiple clusters being assigned to the same cell type and clusters representing the same cell type being present in both non-affected and affected samples.

The count-level expression of all the cells in the clusters annotated with a cell type were summed, then RPKM-normalized (using hg19 assembly) to generate the final cell type signature. Affected cell types/clusters were used in the signature if they were not found in data from the non-affected individual (NEC and Neuron) or if we were not certain about their annotation (cluster 11). Cluster 6 from the affected individual was excluded in the signature generation as it had conflicting cell type assignment evidence to INTER and CN, both of which had representatives via cluster 11 (having weak evidence of being INTER) or non-affected clusters.

#### Preprocessing the bulk RNA-seq data of affected and non-affected groups

Pseudo-bulk mixtures were generated from the scRNA-seq data of the non-affected (DPD4) and affected (DPD1) individuals separately. One hundred pseudo-bulk mixtures were generated per individual by randomly sampling 100 cells without replacement and summing their count-level expression followed by RPKM-normalization. Furthermore, genes with zero expression in more than 20% of the samples were removed. Similarly, bulk RNA-seq profiles of non-affected and affected organoids (from 48 and 72 individuals respectively) were RPKM-normalized and filtered separately for the genes with zero expression in more than 20% of the samples. The bulk RNA-seq of affected and non-affected samples were batch corrected using ComBat (43) to affected and non-affected pseudo-bulk mixtures, respectively. The deconvolution was performed using the batch-corrected bulk RNA-seq data for each group separately.

#### Differential gene expression analysis, clustering of genes, and gene set characterization

After deconvolving the batch-corrected bulk RNA-seq samples, for each cell type, differential gene expression (DGE) analysis was performed with log2-transformed expression values for affected vs non-affected group using Limma package (44). For cell type pairs (CBC-INTER and CN-Neuron) whose inferred proportion sum matched their true proportion sum-derived from the single cell data-DGE analysis was performed for the sum of their deconvolved bulk profiles. After DGE analysis, genes were clustered into 10 groups by k-means algorithm using their discretized expression log fold-change (logFC) for the nine cell types. The discretization was performed as follows: first, genes with absolute expression logFC greater than 5 in any cell type were excluded. Then the logFC of the remaining genes was discretized to values ±2, ±0.2, and 0 according to the following assignment rule,

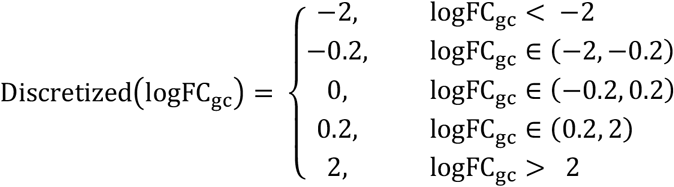

where g and c are gene and cell type indices respectively.

Gene set characterization was performed using the David tool (45,46) for each set of DE genes and cluster of genes. For each annotation cluster, the significant GO-term (FDR < 0.05) with the least FDR was only considered.

#### Model

BEDwARS is a Bayesian probabilistic model for cell type proportion deconvolution specifically designed to adjust for deviations between the reference cell type signatures and the true cell type signatures. The deconvolution is formulated as

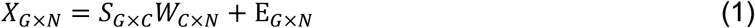

where *G*, *C*, and *N* represent the number of genes, cell types and samples with bulk expression profiles, respectively. *X* is the bulk expression matrix to be deconvolved, with each column being a *G*-dimensional vector and each dimension representing the bulk expression of a gene in a sample. *S* is the “true” (but unknown) signature matrix, with each column being the *G*-dimensional expression signature of a cell type. *W* is the (unknown) proportions matrix, with each column being a *C*-dimensional vector and each dimension representing the proportion of a cell type in a sample. *E* contains the unmodelled bulk gene expression noise which has N(0, *σ*) distribution for all genes and bulk samples. Prior distributions for *S* and *W* are defined as follows:

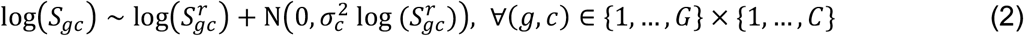

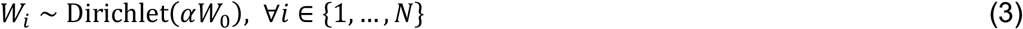

where 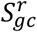 is the reference expression of gene *g* in cell type *c*. Equation (2) is the key modeling assumption addressing the deviations between the known reference signature and the unknown true signature underlying the bulk profiles *X*. It states that the (log transformed) expression of a gene in a cell type deviates from the corresponding value in the reference signature by an amount that is Normally distributed with zero mean and a variance that is gene- as well as cell type-dependent. This variance term, 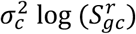 in Equation (2), is proportional to the (log transformed) reference signature value, thus allowing greater deviations for more abundant genes, and the constant of proportionality 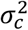 is cell type-dependent, allowing different cell types to exhibit globally more or less deviations. *W*_0_, a *C*-dimensional probability vector is the mean of a Dirichlet distribution and can be set by user based on prior knowledge of cell type proportions. However, as such information is not commonly available, the value of *W*_0_ is set to 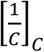 by default. *α* controls the variance of the Dirichlet distribution and its high values are associated with low variation. We also defined priors for the parameters of the distributions above,

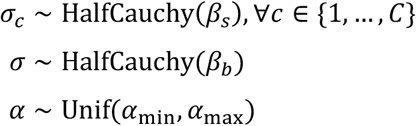

where *β_s_*, *β_b_*, *α*_min_ and *α*_max_ were set to 1, 5, 0 and 30 for all the tests performed. These values can be set by user. For example, *β_s_* can be set to smaller values to reduce the amount of perturbation added to the reference signature. The Half Cauchy prior was used for the standard deviation as suggested by Gelman (47).

Inference of parameters was done by maximizing the posterior probability of all parameters given the bulk expression profiles *X*,

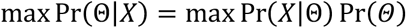

where 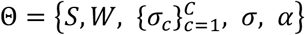. Metropolis-Hastings (MH) sampling was used for maximum *a posteriori* estimation. In implementing MH algorithm, multiple chains were run in parallel to sample from the posterior distribution. In all chains *S* was initialized with the reference signature, *α* and *σ* were initialized by sampling from their corresponding priors and columns of *W* were initialized with *W*_0_. For 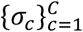, equal number of chains were initialized with 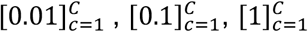 and by sampling from the prior. The “best” chain was selected to estimate the parameters. The criterion for selecting the best chain was the mean squared error between *X* and 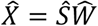, restricted to marker genes identified using the reference signature. For all tests reported in this work, the set of genes with at least four-fold higher expression in one cell type compared to the others were chosen as markers. The inference of parameters was done by averaging over the samples drawn by the best chain after its burn-in period. BEDwARS runs multiple chains in parallel on GPU and is implemented in PyTorch. The number of chains used for benchmarking and DPD-deficiency deconvolution experiments were set to 150 and 100, respectively.

## Declarations

### Ethics approval and consent to participate

This study was approved and was overseen by the institutional review board of Mayo Clinic (IRB study ID: 14-005685). Written informed consent was obtained from all study subjects. In the case of study subjects under the age of 18, written consent was obtained from parents and assent was provided by the subject for study participation.

### Consent for publication

Not Applicable

### Availability of data and materials

The datasets used in this study were downloaded from the following links,

Baron (15): https://www.ncbi.nlm.nih.gov/geo/query/acc.cgi?acc=GSE84133

Segrestolpe (14): https://www.ebi.ac.uk/arrayexpress/experiments/E-MTAB-5061/

Enge (16): https://www.ncbi.nlm.nih.gov/geo/query/acc.cgi?acc=GSE81547

Darmanis (17): https://github.com/VCCRI/CIDR-comparisons/tree/master/Brain/Data

IP (18): https://www.ncbi.nlm.nih.gov/geo/query/acc.cgi?acc=GSE73721

CA (20): https://portal.brain-map.org/atlases-and-data/rnaseq/human-mtg-smart-seq

NG (21): Sutton et al. Supplementary Data 5

MM (19): Data was shared by Steven Sloan contacted by email (https://www.brainrnaseq.org)

Newly generated sequence data used for this study are deposited in the NIH Sequence Read Archive (SRA) under BioProject PRJNA888085.

For more details on implementation and usage of BEDwARS, please visit https://github.com/sabagh1994/BEDwARS.

### Author’s contributions

SG, SS, and SMO designed the study and wrote the manuscript. SG designed and implemented the algorithm and performed the data analyses. ES contributed to the algorithm implementation. RES and KJB performed laboratory experiments. The authors read and approved the final manuscript.

## Acknowledgements

Not Applicable

## Additional Files Information

**Additional File 1. Supplementary.pdf. Supplementary Figures and Tables.**

**Additional File 2. Table_S2.xlsx. Differential gene expression analysis for DPD deficiency**. The summary of differential gene expression analysis using Limma package for bulk, pseudo-bulk, and deconvolved cell type or pairs of cell types (CN-NEU, CBC-INTER) expression profiles (**“DGE”**). Top 200 DE genes per cell type or pairs of cell types are listed in **“Top 200 DE genes per cell type”** sheet.

**Additional File 3. Table_S4.xlsx. Summary of David gene set characterization performed on top 200 DE genes identified by DGE analysis for bulk, bootstrapped pseudo-bulk and deconvolved cell type expression profiles for DPD deficiency**. In each annotation cluster the first GO with significant FDR (FDR < 0.05), highlighted with yellow, was considered.

**Additional File 4. Table_S5.xlsx. Cluster of genes identified by Kmeans clustering based on the pattern of genes’ differential expression across the cell types for DPD deficiency**. Each column represents the genes belonging to the same cluster.

**Additional File 5. Table_S6.xlsx. Summary of David gene set characterization for cluster of genes identified by Kmeans algorithm based on the pattern of genes’ differential expression across the cell types for DPD deficiency**. The David results for each cluster are included in a sheet named with the cluster name (Cluster X). Clusters zero and four were excluded as they had more than 2000 genes. David results for top 200 DE genes identified by DGE analysis on bulk and bootstrapped pseudo-bulk are included for convenient comparison.

**Additional File 6. Table_S7.xlsx. Top 200 markers per cluster of non-affected and affected cells for DPD deficiency**. Each column contains the top 200 filtered marker genes for a cluster of cells. These markers were used for assigning cell types to the clusters.

**Additional File 7. Table_S9.xlsx. Table S9. KnowEng gene set characterization performed on the markers of a subset of cell clusters for DPD deficiency**. These results were used to assign cell types to cluster of non-affected and affected cells. The GO terms that were enriched and used for cell type assignment are summarized in **Table S8** for easier lookup.

