## Additional File 1 for "A Robust Bayesian Approach to Bulk Gene Expression Deconvolution with Noisy Reference Signatures"

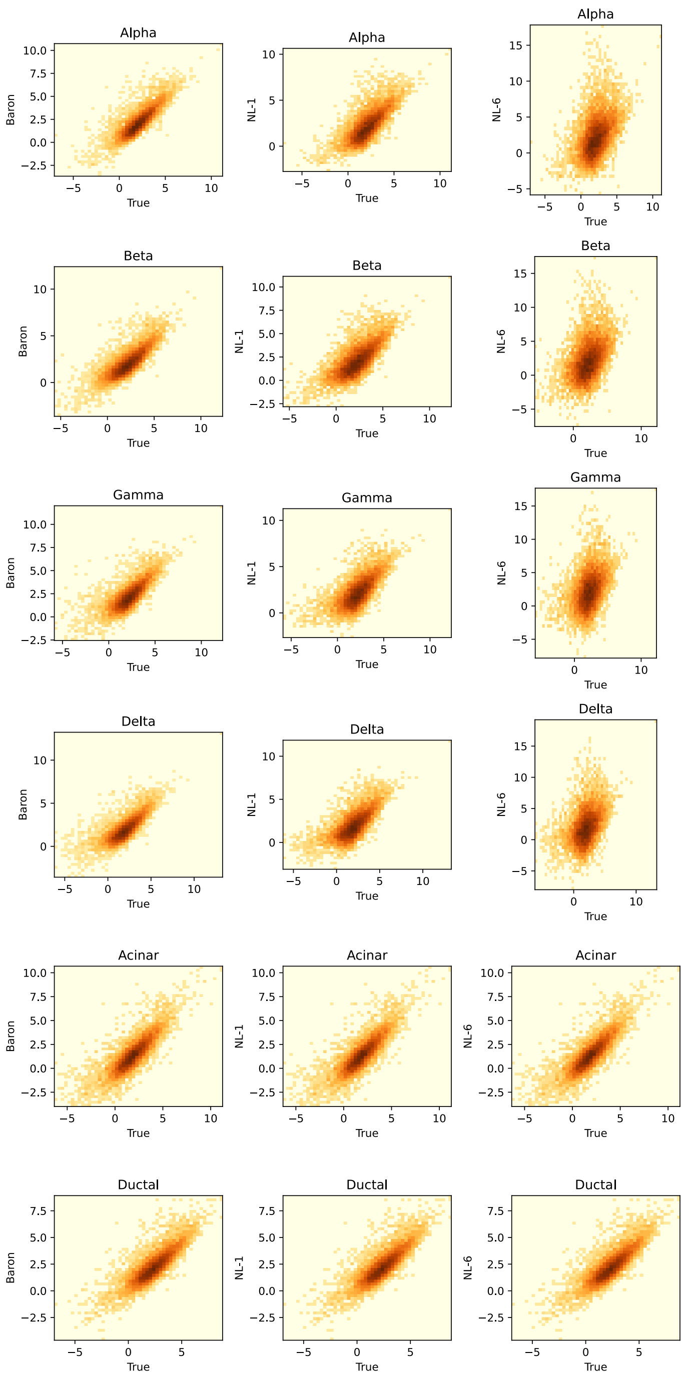


**Figure S1. Relationship of Baron and its two noisy variants with the true Segerstolpe-H signature.** For each reference signature ("Baron" and its two variants – "NL-1" and "NL-6", representing the lowest and highest levels of added noise respectively), each gene's expression value in that signature is shown on the y-axis and its expression in the true signature (Segerstolpe-H) is shown on the x-axis; both are in log scale. As the noise increases the signatures are less similar to the true signature. For acinar and ductal cell types, noise is not added to the Baron signature, so the noisy signature (NL-1, NL-6) relationship with the true signature is the same as that of the Baron signature.

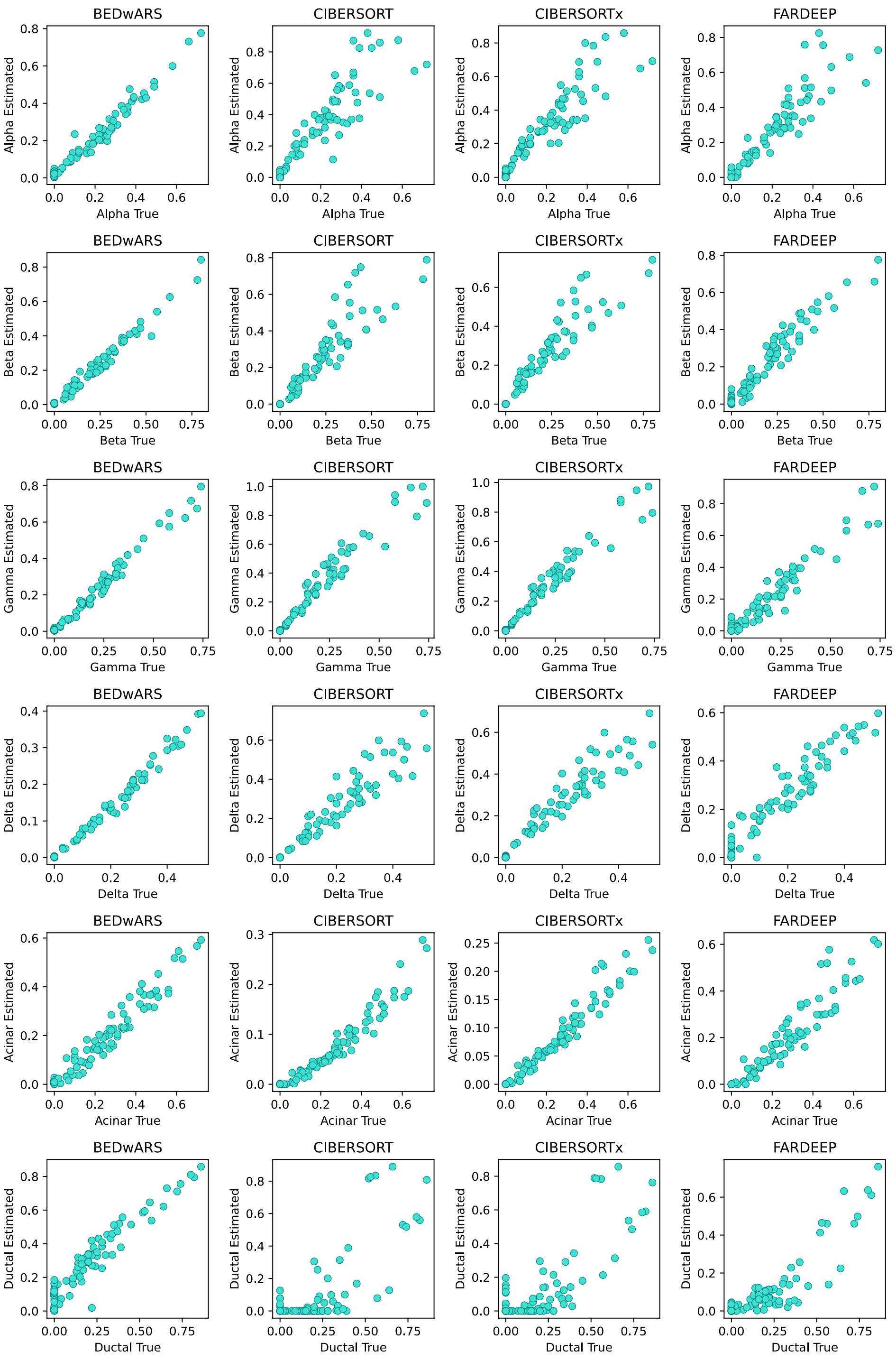


**Figure S2. Quality of cell type proportion inference using Baron signature in deconvolving pseudo-bulk samples from Segerstolpe-H.** The estimated and true proportions for each of 100 pseudo-bulk samples are compared, for all cell types and all methods (BEDwARS, CIBERSORT, CIBERSORTx, FARDEEP). BEDwARS estimations match the true values better than the other methods for all cell types. CIBERSORT(x) underestimates the acinar proportions by a factor of 2 even though the estimated and true proportions are highly correlated. There is a severe underestimation of ductal proportions by all methods except BEDwARS.

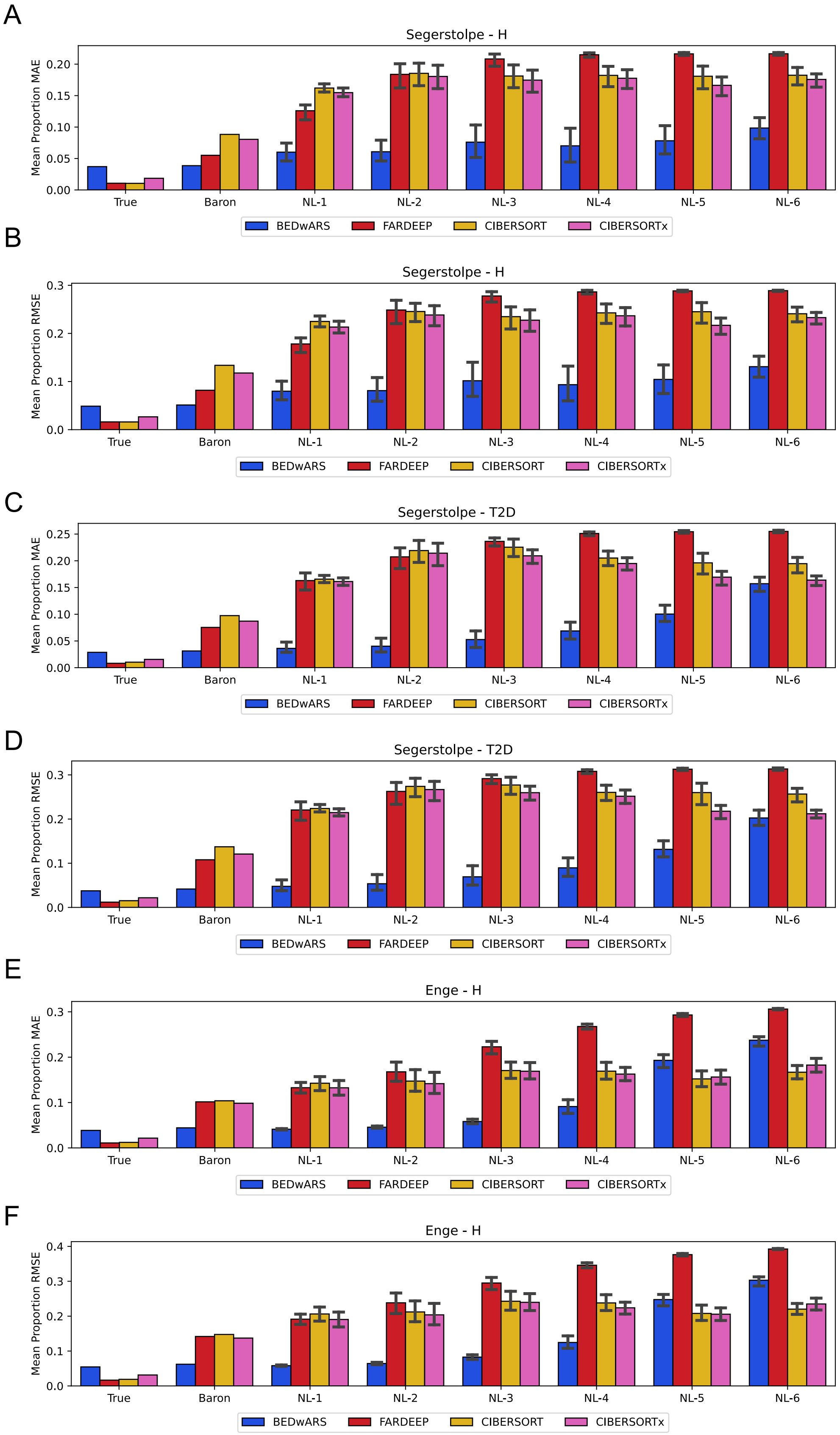
A
Segerstolpe - H
Mean Proportion MAE
B
Segerstolpe - H
Mean Proportion RMSE
C
Segerstolpe - T2D
Mean Proportion MAE
D
Segerstolpe - T2D
Mean Proportion RMSE
E
Enge - H
Mean Proportion MAE
F
Enge - H
Mean Proportion RMSE
True
Baron
NL-1
NL-2
NL-3
NL-4
NL-5
NL-6
BEDwARS
FARDEEP
CIBERSORT
CIBERSORTx

**Figure S3. Performance evaluation, by RMSE and MAE criteria, of methods for estimation of cell type proportions by deconvolving pancreatic pseudo-bulk profiles.** Shown are MAE (mean absolute error) (**A, C, E**) and RMSE (root mean squared error) (**B, D, F**) between estimated and true proportions, averaged over the cell types, for all four methods – BEDwARS (this work), FARDEEP [CITE], CIBERSORT [CITE], CIBERSORTx [CITE] – tested by us. RMSE and MAE are computed between the inferred and true cell type proportions of 100 pseudo-bulk samples derived from Segerstolpe-H (**A, B**), Segerstolpe-T2D (**C, D**), and Enge-H (**E, F**) data sets. Evaluations are shown with the Baron reference signature as well as its noisy variants (NL-1, NL-2, … NL-6). BEDwARS has the least RMSE and MAE in Baron group and all noise levels except for the largest two noise levels in (**E, F**). Evaluations are also shown for the hypothetical case where the true underlying signature was available during deconvolution ("True" category in each panel), though this is not a common situation.

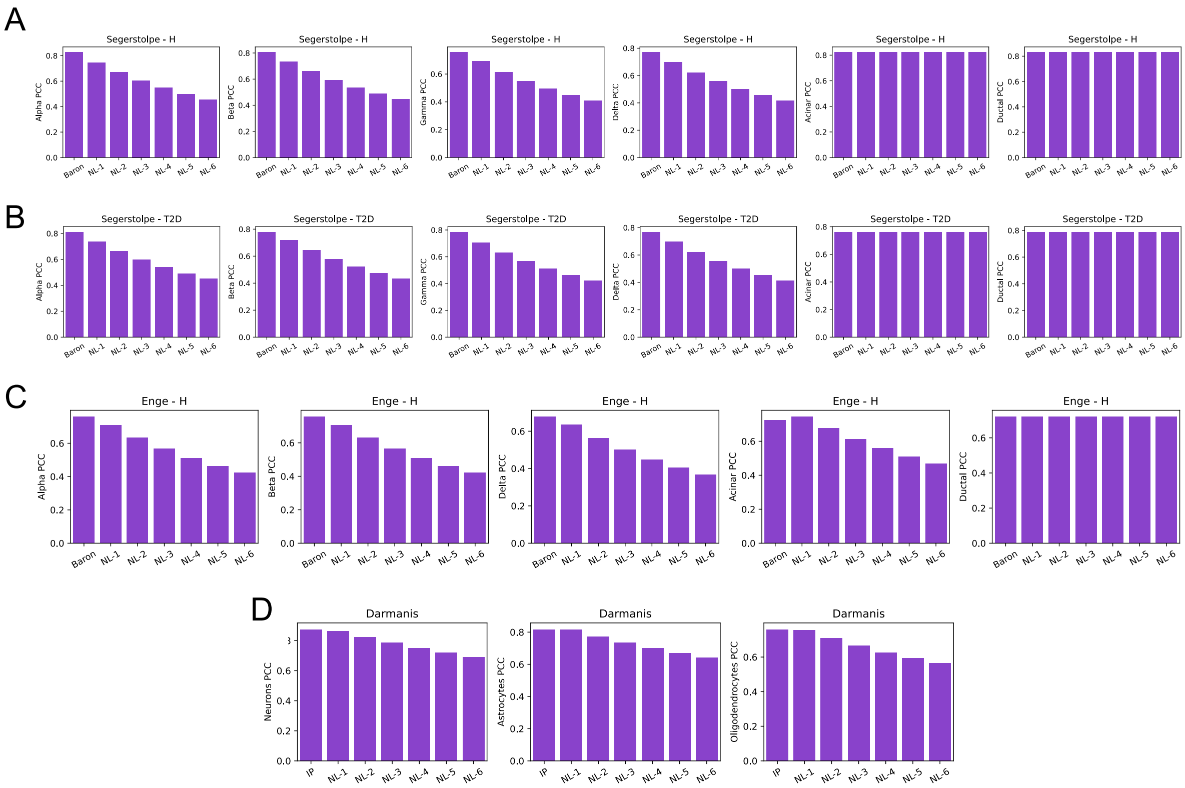


**Figure S4. Correlation between reference and true signature at different noise levels, for three pancreatin data sets (A,B,C) and one brain data set (D).** Each panel corresponds to a cell type, showing median Pearson correlation coefficient (PCC) between log-transformed reference and true signatures of that cell type, at varying noise levels. In each panel, the first category ("Baron" for A-C and "IP" for D) represents the reference signature without added noise and the remaining categories ("NL-X") represent noisy versions of this signature, with higher values of "X" indicating greater added noise. As the noise level increases, the noisy reference and true signatures become more dissimilar. To avoid large deviations, the signatures of cell types acinar and ductal are not perturbed when creating noisy reference signatures for Segerstolpe-H (A**)** and Segerstolpe-T2D (B) data sets. For the same reason, ductal signature is not perturbed in the Enge-H (C) data set.

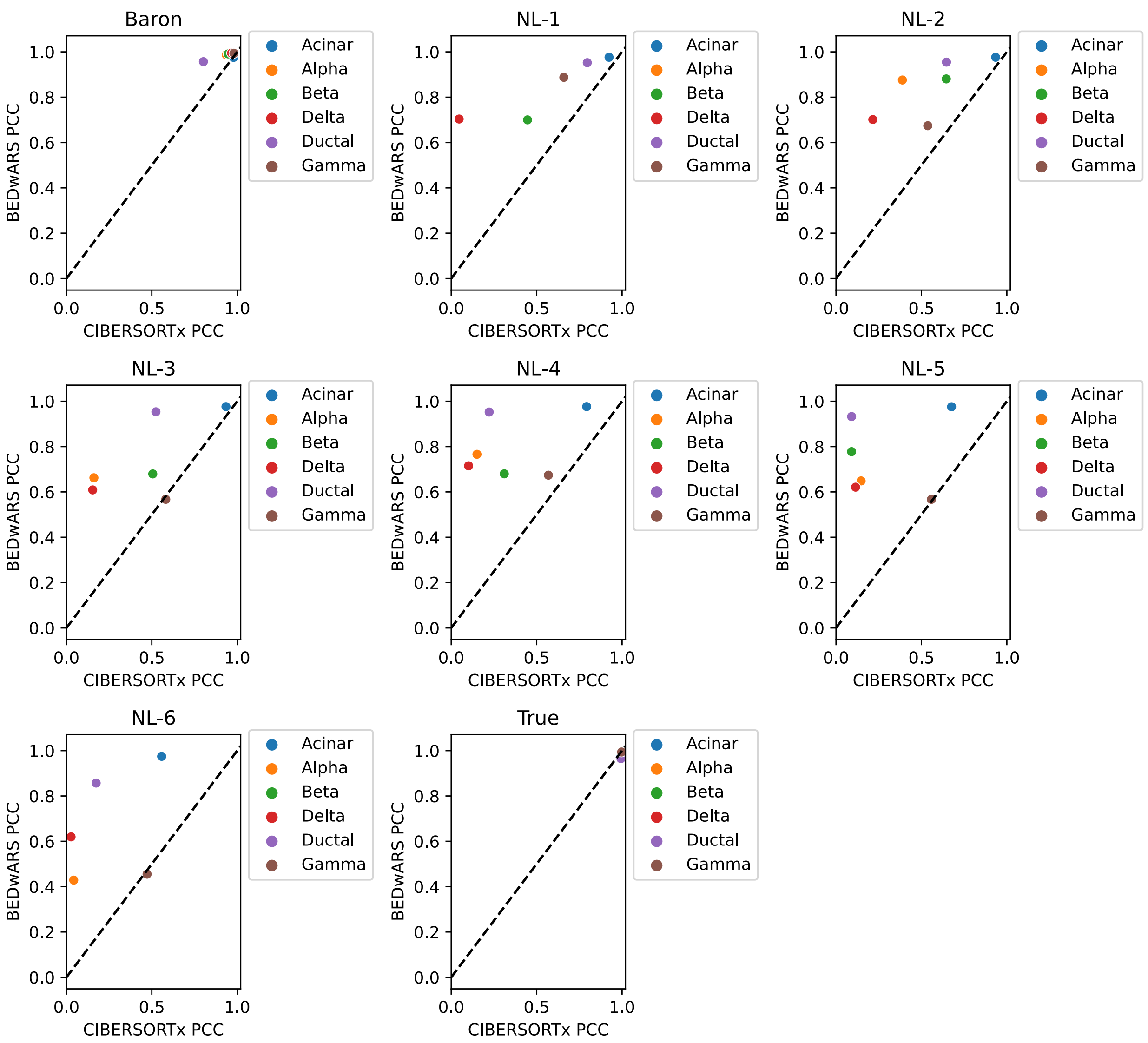


**Figure S5. Cell type level comparison of BEDwARS and CIBERSORTx for the task of proportion estimation by deconvolution of Segerstolpe-H pseudo-bulk samples.** Different panels correspond to different reference signatures used during deconvolution -- the Baron signature and its noisy variants NL-1, NL-2, … NL-6, as well as the true signature. Performance is measured by the Pearson Correlation Coefficient (PCC) between estimated and true proportions of a cell type, across the 100 pseudo-bulk samples. For evaluations with noisy signatures, the average over 11 noisy signatures at the same noise level is shown. The performance gap between BEDwARS and CIBERSORTx is large at all noise levels for most of the cell types. The performance gap is small in the case of the Baron signature, and absent when the true signature is used.

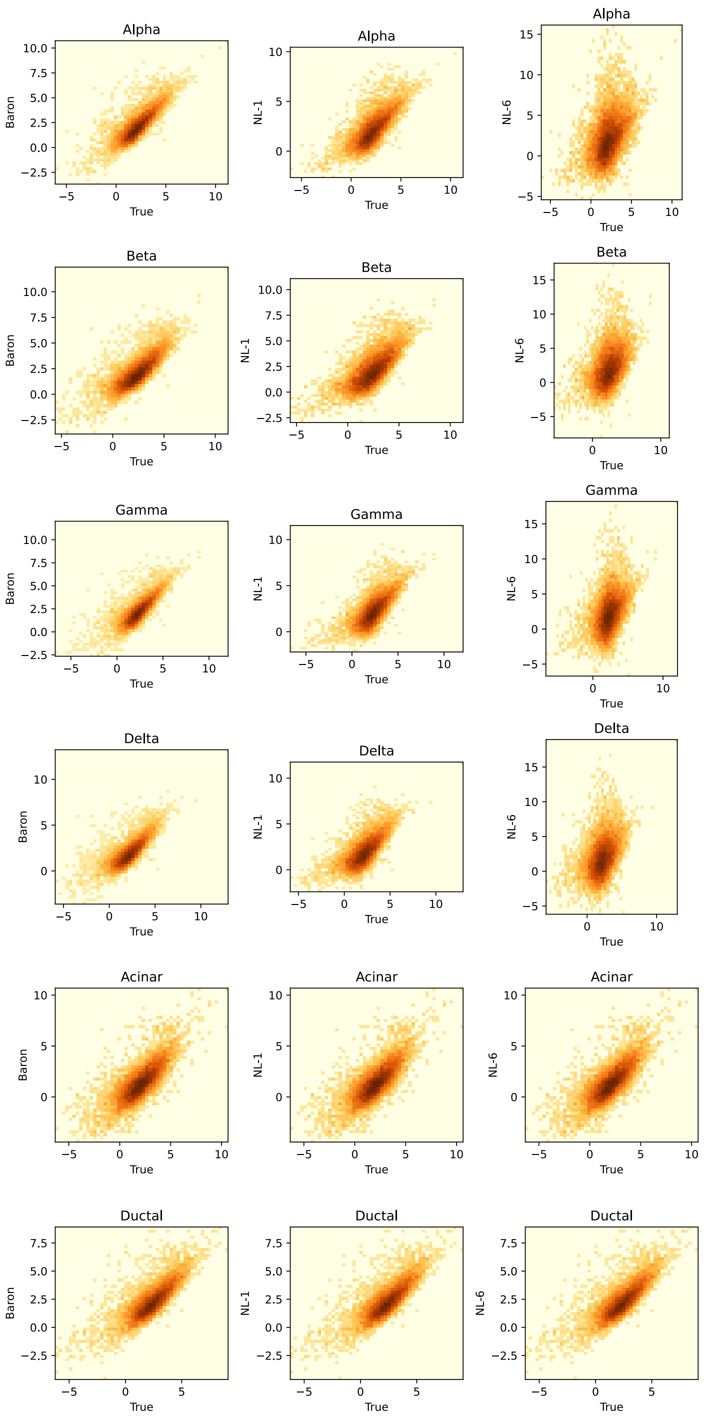


**Figure S6. Relationship of Baron and its two noisy variants with the true Segerstolpe-T2D signature.** For each reference signature ("Baron" and its two variants – "NL-1" and "NL-6", representing the lowest and highest levels of added noise respectively), each gene's expression value in that signature is shown on the y-axis and its expression in the true signature (Segerstolpe-

T2D) is shown on the x-axis; both are in log scale. As the noise increases the signatures are less similar to the true signature. For acinar and ductal cell types, noise is not added to the Baron signature, so the noisy signature (NL-1, NL-6) relationship with the true signature is the same as that of the Baron signature.

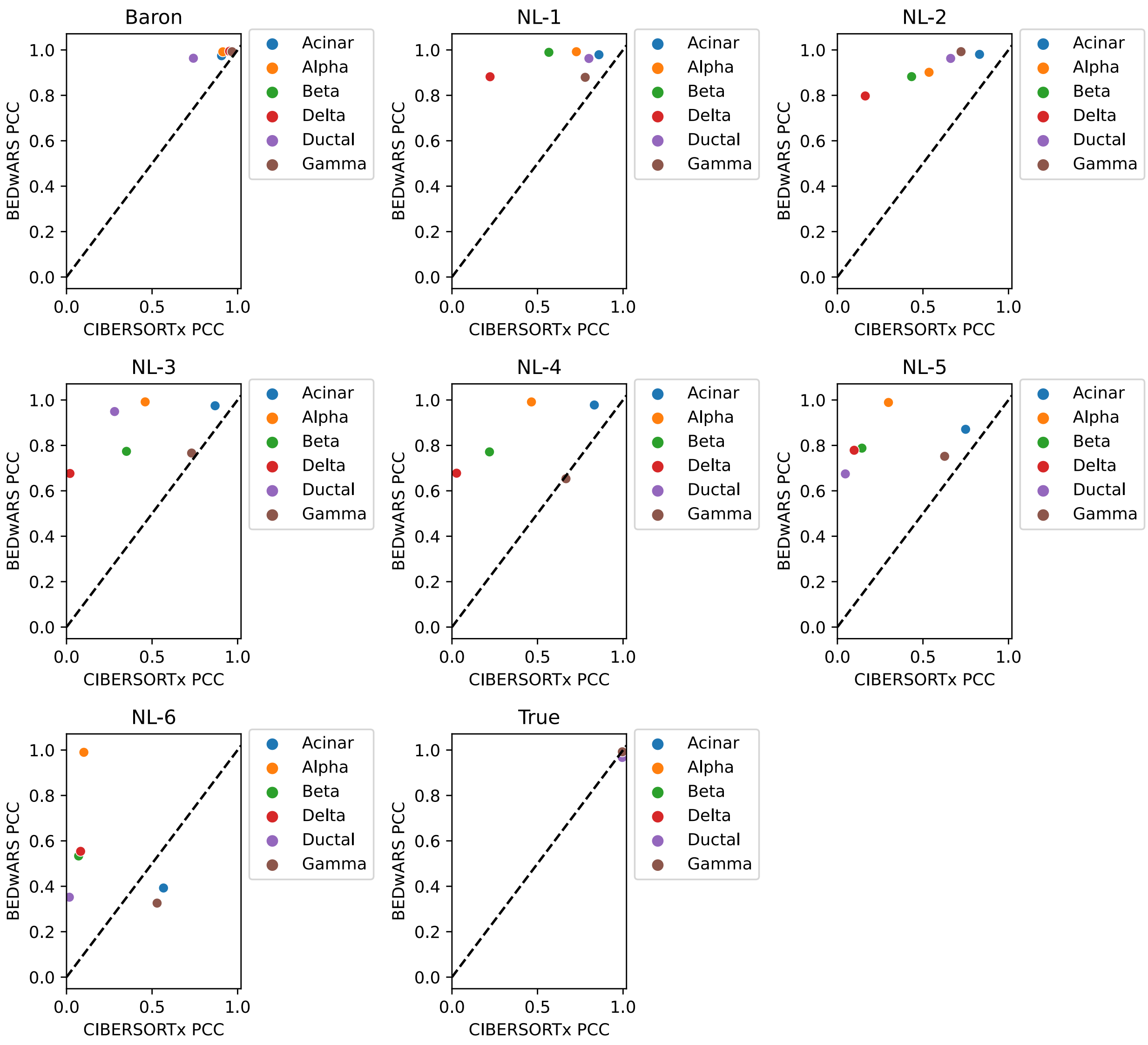


**Figure S7. Cell type level comparison of BEDwARS and CIBERSORTx for the task of proportion estimation by deconvolution of Segerstolpe-T2D pseudo-bulk samples.** Different panels correspond to different reference signatures used during deconvolution -- the Baron signature and its noisy variants NL-1, NL-2, … NL-6, as well as the true signature. Performance is measured by the Pearson Correlation Coefficient (PCC) between estimated and true proportions of a cell type, across the 100 pseudo-bulk samples. For evaluations with noisy signatures, the average over 11 noisy signatures at the same noise level is shown. The performance gap between BEDwARS and CIBERSORTx is large at all noise levels for most of the cell types. The performance gap is small in the case of the Baron signature, and absent when the true signature is used.

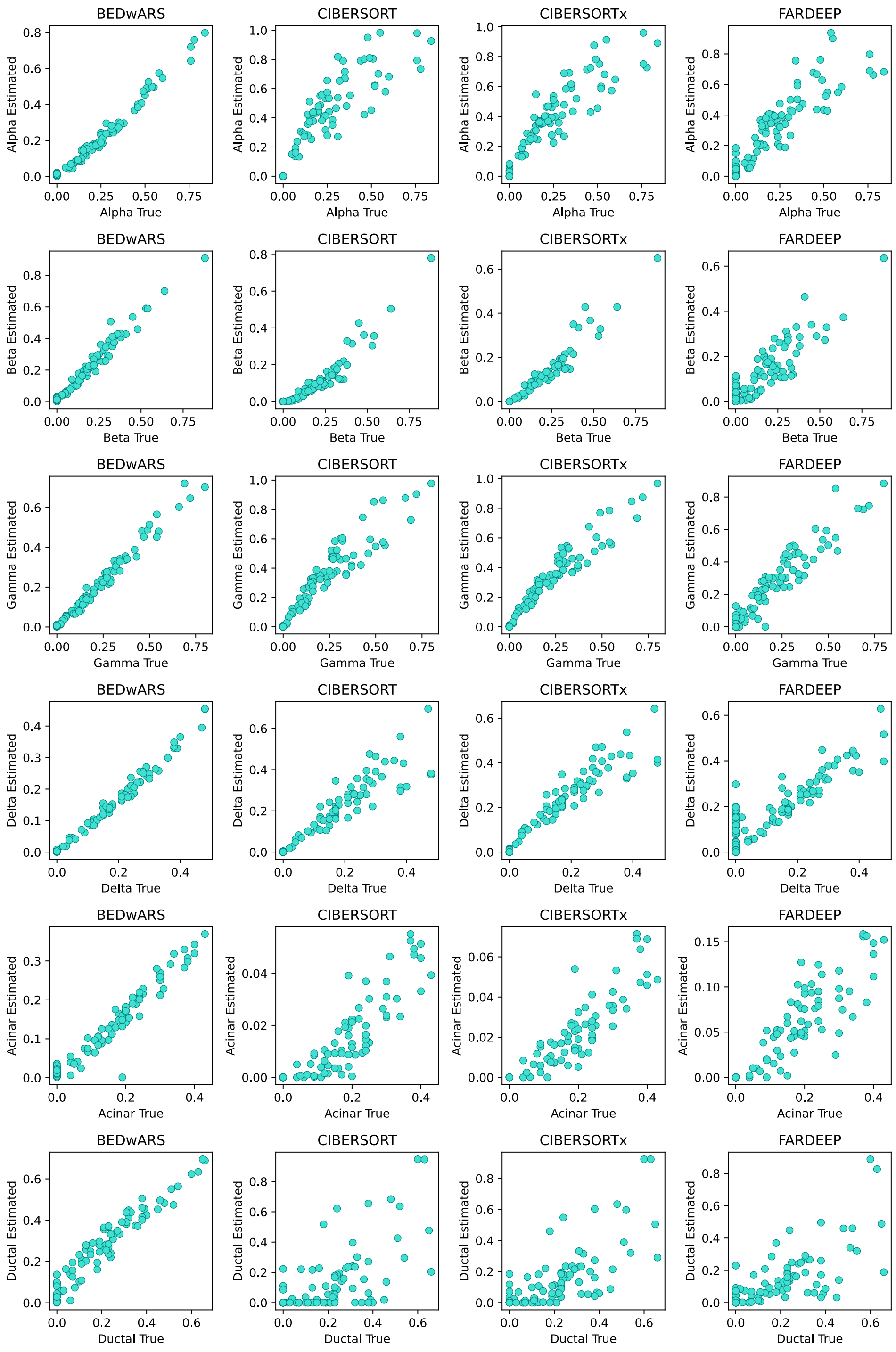


**Figure S8. Quality of cell type proportion inference using Baron signature in deconvolving pseudo-bulk samples from Segerstolpe-T2D.** The estimated and true proportions for each of 100 pseudo-bulk samples are compared, for all cell types and all methods (BEDwARS, CIBERSORT, CIBERSORTx, FARDEEP). BEDwARS estimations match the true values better than the other methods for all cell types. CIBERSORT(x) overestimates proportions of alpha, gamma, and delta cell types and underestimate acinar and ductal proportions. FARDEEP also frequently overestimates the small proportions and severely underestimates proportions of acinar and ductal cell types.

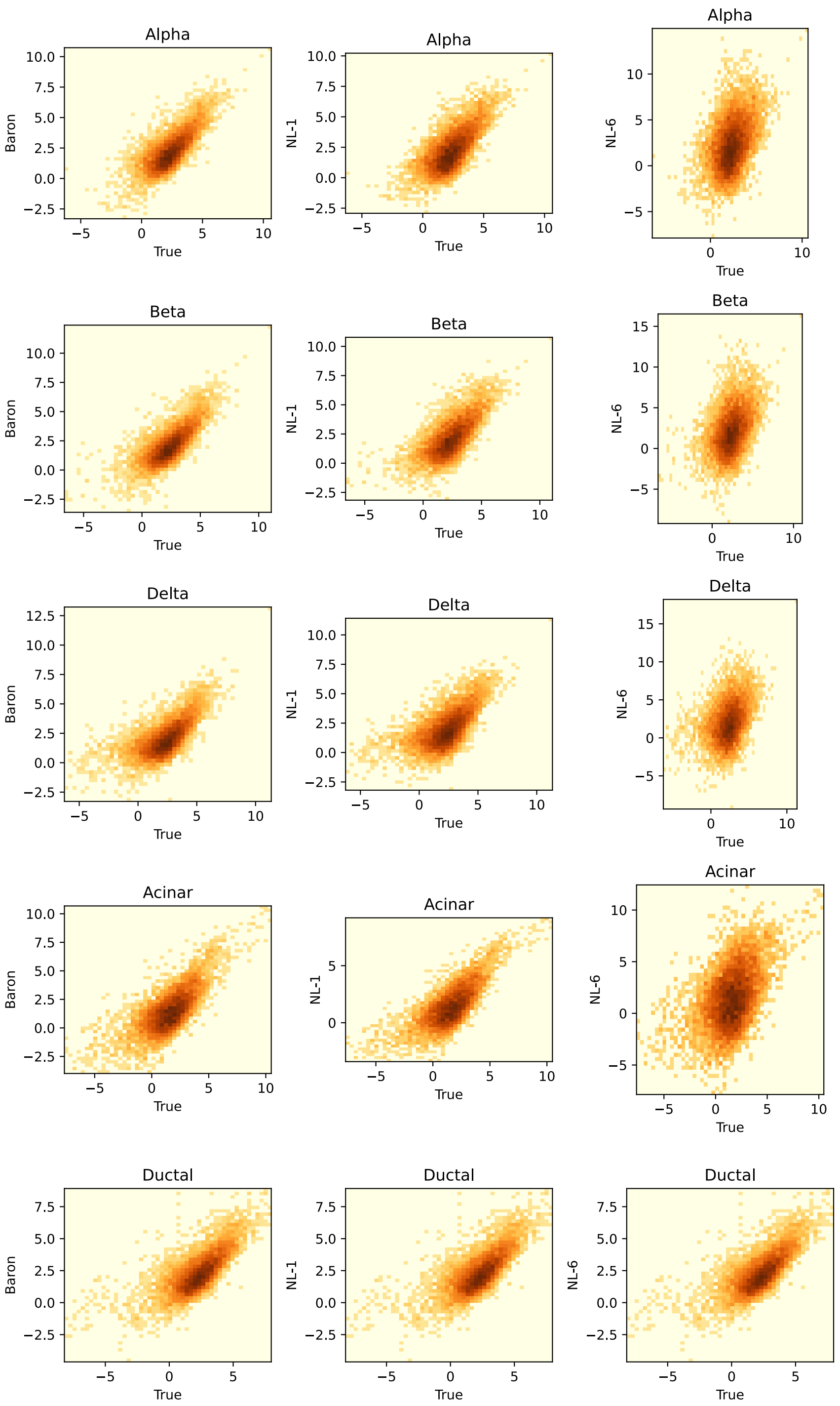


**Figure S9. Relationship of Baron and its two noisy variants with the true Enge-H signature.** For each reference signature ("Baron" and its two variants – "NL-1" and "NL-6", representing the lowest and highest levels of added noise respectively), each gene's expression value in that signature is shown on the y-axis and its expression in the true signature (Enge-H) is shown on the x-axis; both are in log scale. As the noise increases the signatures are less similar to the true signature. For ductal cell type, noise is not added to the Baron signature, so the noisy signature (NL-1, NL-6) relationship with the true signature is the same as that of the Baron signature.

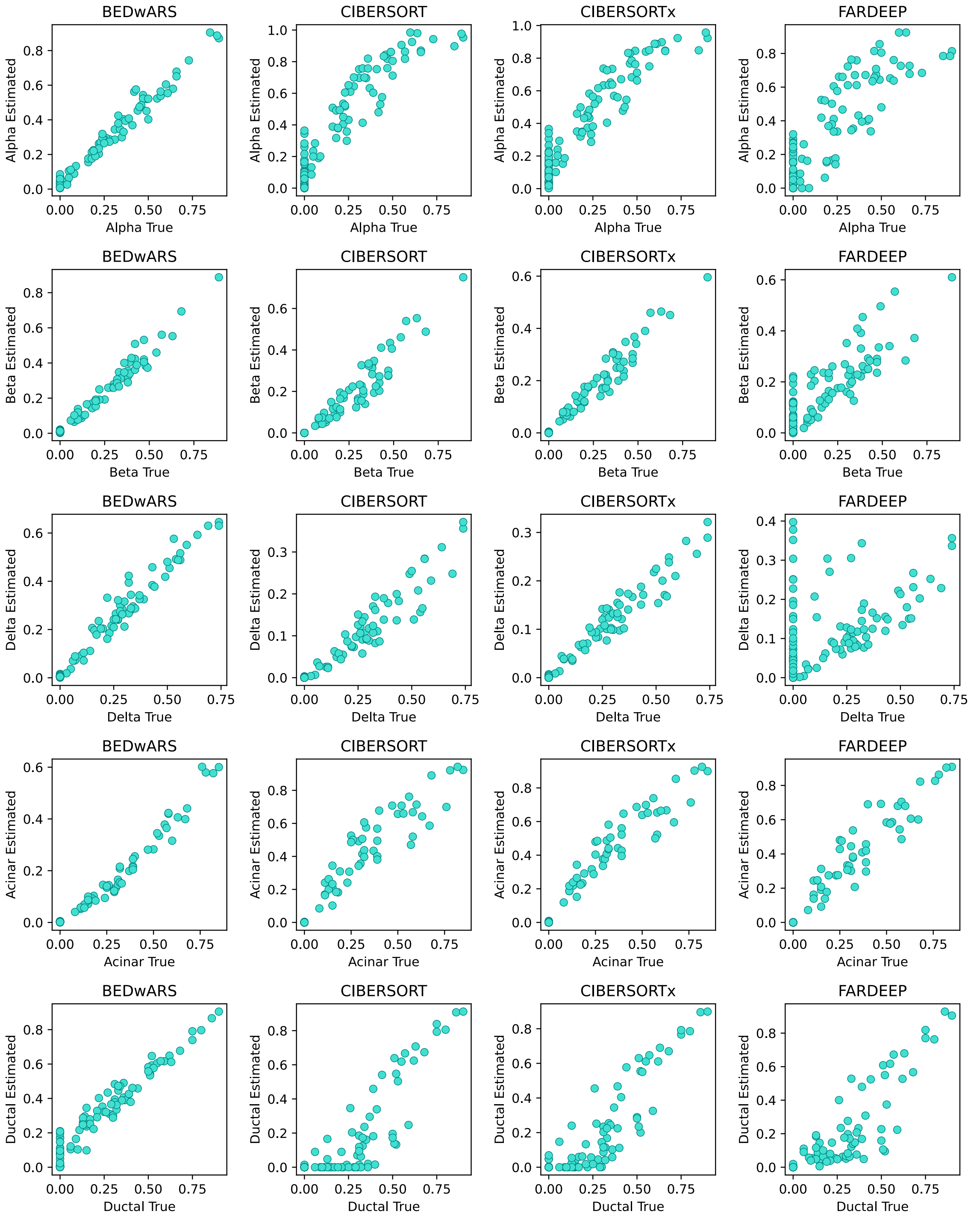


**Figure S10. Quality of cell type proportion inference using Baron signature in deconvolving pseudo-bulk samples from Enge-H.** The estimated and true proportions for each of 100 pseudo-bulk samples are compared, for all cell types and all methods (BEDwARS, CIBERSORT, CIBERSORTx, FARDEEP). BEDwARS estimations match the true values better than the other methods for all cell types. All methods except BEDwARS show a severe overestimation of zero-valued alpha proportions. CIBERSORT(x) underestimates delta proportions by 2 folds. FARDEEP significantly overestimates the zero-valued delta proportions and underestimates the highest delta proportions by 2 folds.

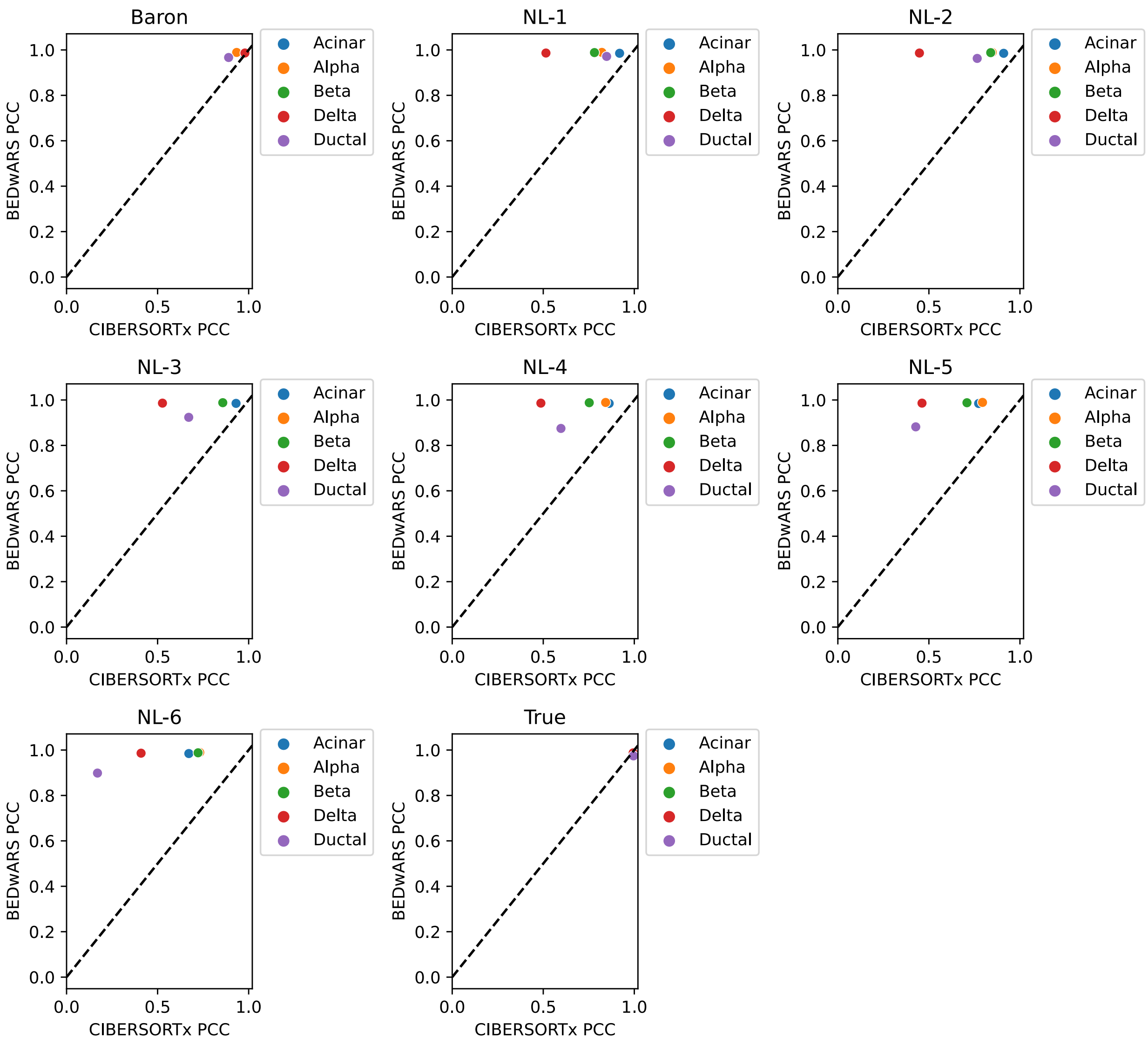


**Figure S11. Cell type level comparison of BEDwARS and CIBERSORTx for the task of proportion estimation by deconvolution of Enge-H pseudo-bulk samples.** Different panels correspond to different reference signatures used during deconvolution -- the Baron signature and its noisy variants NL-1, NL-2, … NL-6, as well as the true signature. Performance is measured by the Pearson Correlation Coefficient (PCC) between estimated and true proportions of a cell type, across the 100 pseudo-bulk samples. For evaluations with noisy signatures, the average over 11 noisy signatures at the same noise level is shown. The performance gap between BEDwARS and CIBERSORTx is large at all noise levels for most of the cell types. The performance gap is small in the case of the Baron signature, and absent when the true signature is used.

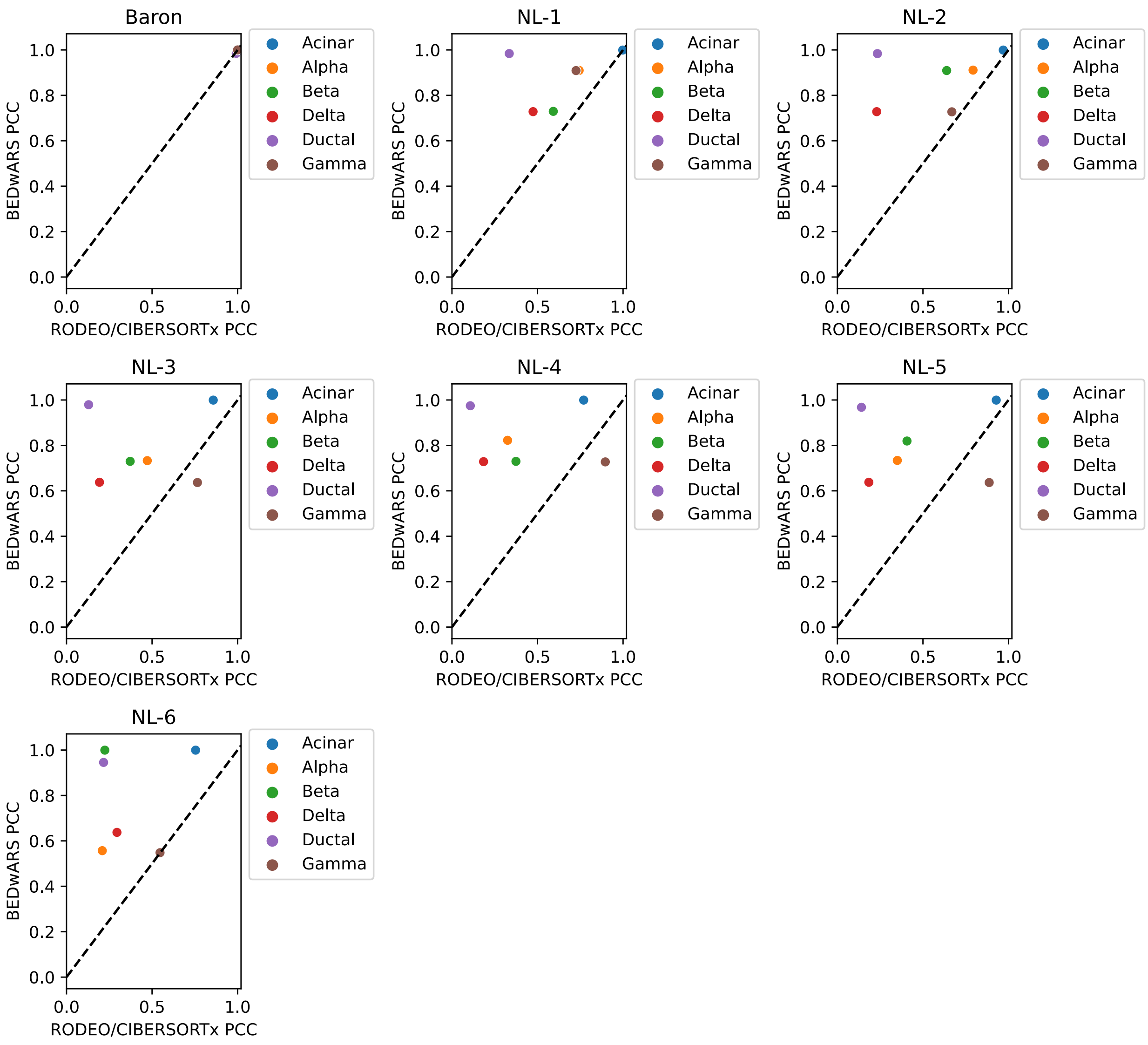


**Figure S12. Cell type level comparison of BEDwARS and RODEO/CIBERSORTx for the task of signature estimation by deconvolution of Segerstolpe-H pseudo-bulk samples.** Different panels correspond to different reference signatures used during deconvolution -- the Baron signature and its noisy variants NL-1, NL-2, … NL-6. Performance is measured by the Pearson Correlation Coefficient (PCC) between estimated and true signatures of a cell type. For evaluations with noisy signatures, the average over 11 noisy signatures at the same noise level is shown. BEDwARS performance is similar to or notably better than RODEO provided with CIBEROSRTx estimates of proportions at all noise levels except for the gamma cell type at noise levels NL-3, NL-4 and NL-5. The performance gap is absent in the case of the Baron signature.

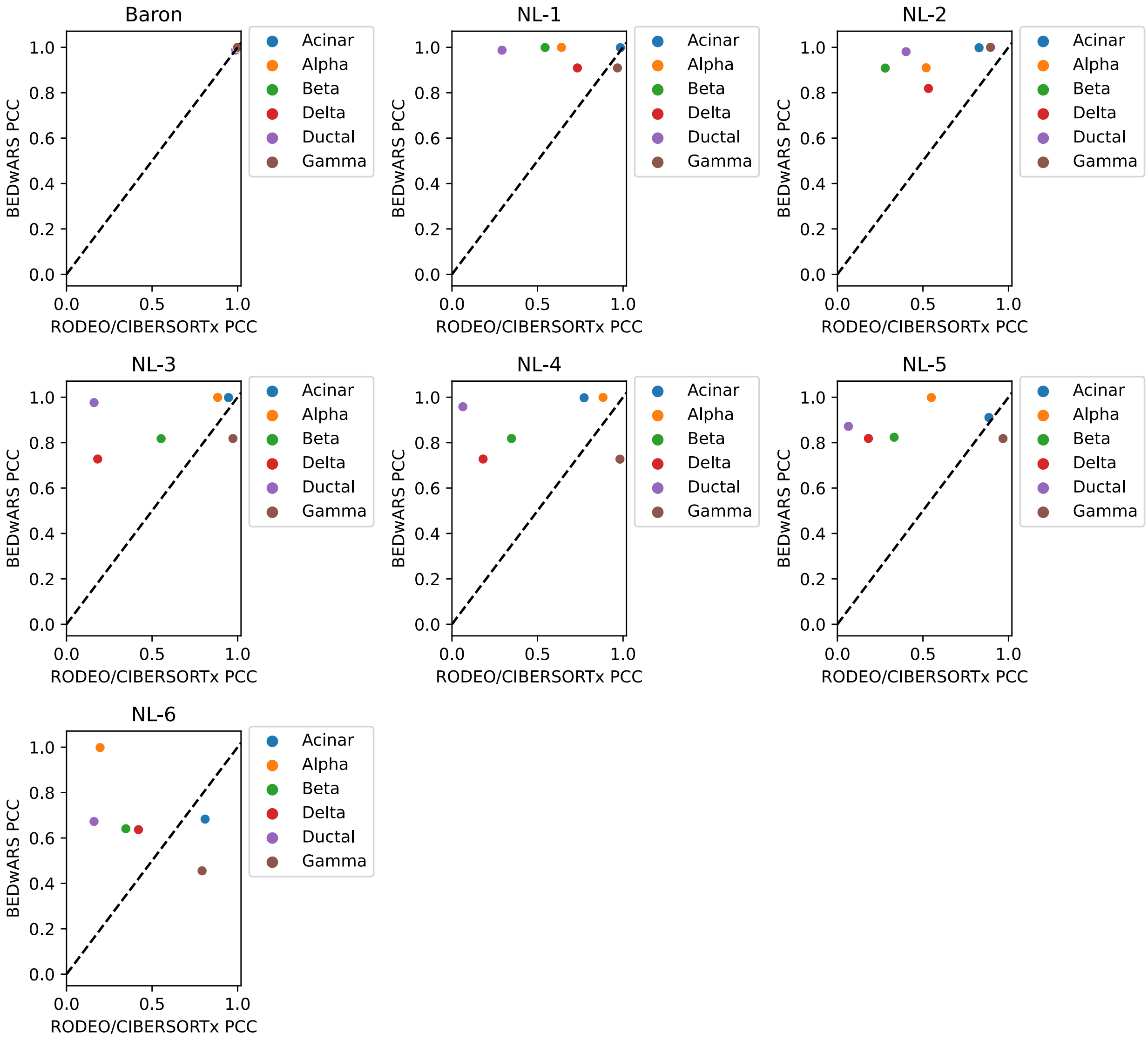


**Figure S13. Cell type level comparison of BEDwARS and RODEO/CIBERSORTx for the task of signature estimation by deconvolution of Segerstolpe-T2D pseudo-bulk samples.** Different panels correspond to different reference signatures used during deconvolution -- the Baron signature and its noisy variants NL-1, NL-2, … NL-6. Performance is measured by the Pearson Correlation Coefficient (PCC) between estimated and true signatures of a cell type. For evaluations with noisy signatures, the average over 11 noisy signatures at the same noise level is shown. BEDwARS performance is similar to or notably better than RODEO provided with CIBERSORTx estimates of proportions at all noise levels and cell types except for gamma cell type at noise levels NL-3, NL-4, NL-5 and NL-6, and acinar at the highest noise level. The performance gap is absent in the case of the Baron signature.

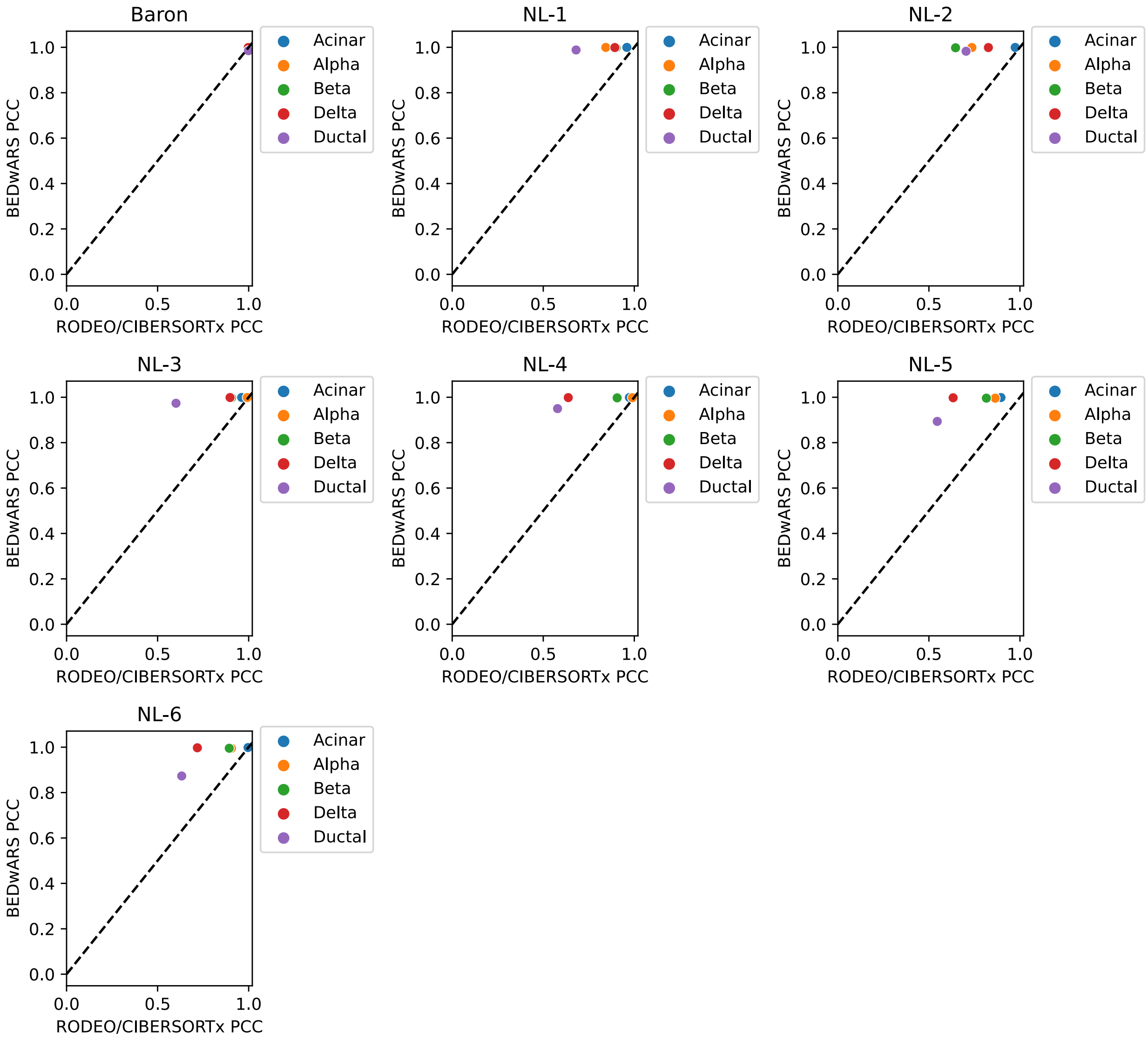


**Figure S14. Cell type level comparison of BEDwARS and RODEO/CIBERSORTx for the task of signature estimation by deconvolution of Enge-H pseudo-bulk samples.** Different panels correspond to different reference signatures used during deconvolution -- the Baron signature and its noisy variants NL-1, NL-2, … NL-6. Performance is measured by the Pearson Correlation Coefficient (PCC) between estimated and true signatures of a cell type. For evaluations with noisy signatures, the average over 11 noisy signatures at the same noise level is shown. BEDwARS outperforms RODEO provided with CIBERSORTx-estimated proportions at all noise levels. When the Baron signature is used (without added noise) both methods perform equally well.

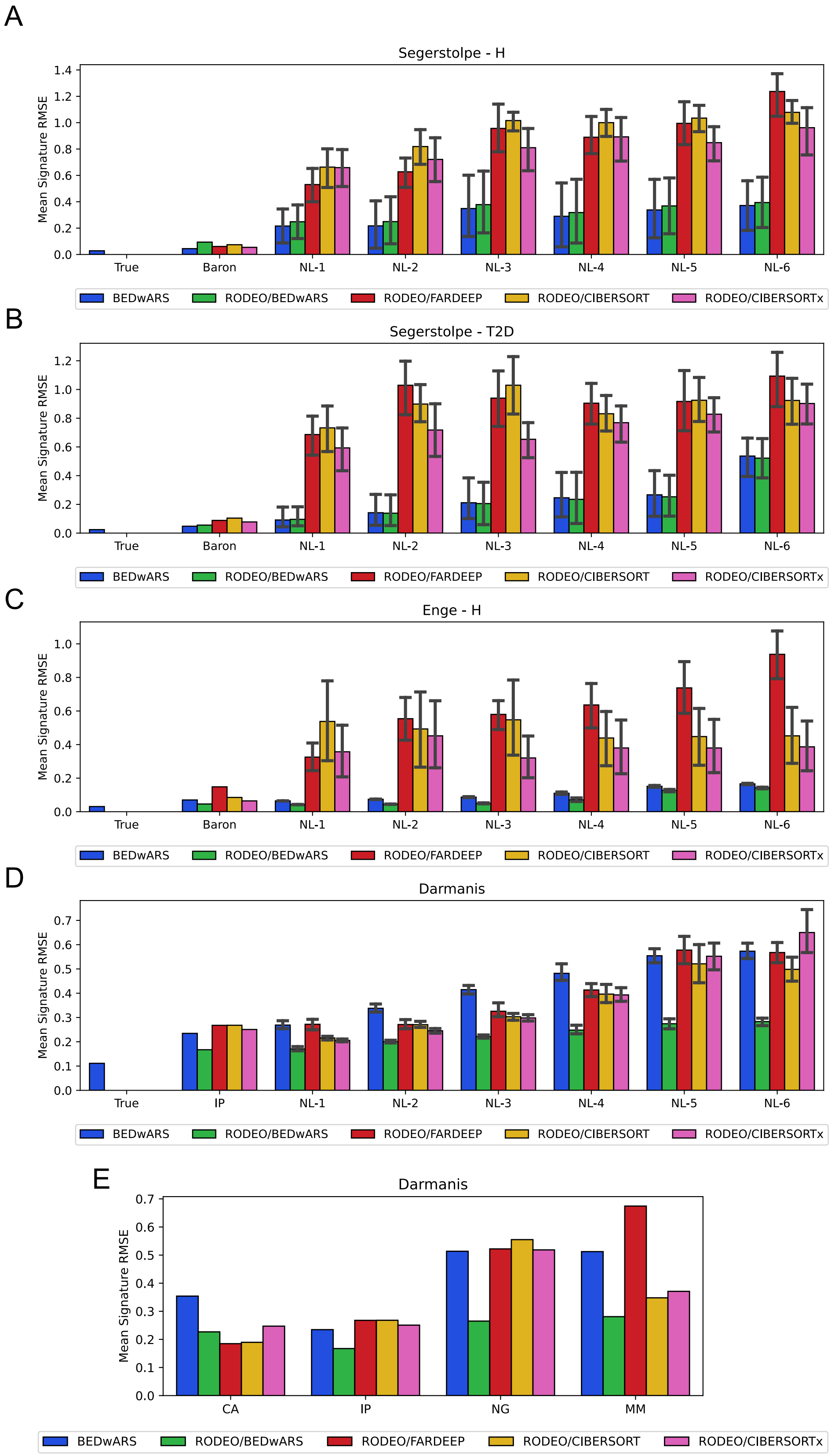

A
Segerstolpe - H
Mean Signature RMSE
True
Baron
NL-1
NL-2
NL-3
NL-4
NL-5
NL-6
BEDwARS
RODEO/BEDwARS
RODEO/FARDEEP
RODEO/CIBERSORT
RODEO/CIBERSORTx
B
Segerstolpe - T2D
C
Enge - H
D
Darmanis
IP
E
Darmanis
CA
IP
NG
MM

**Figure S15. Performance evaluation, by RMSE criterion, of methods for estimating cell type signatures by deconvolving bulk transcriptomic profiles with a given reference signature.** Shown is the RMSE between estimated and true gene expression values, averaged over the cell types, for all five methods tested by us. Four of the methods utilize RODEO after estimating cell type proportions using a deconvolution method – BEDwARS, FARDEEP, CIBERSORT or CIBERSORTx – these are named following the template “RODEO/M” where M is the method used for cell type proportion estimation. The fifth method evaluated is BEDwARS, since BEDwARS simultaneously estimates signatures as well as proportions. RMSE is computed between the inferred and true cell type signatures for the task of deconvolving 100 pseudo-bulk samples generated from Segerstolpe-H (**A**), Segerstolpe-T2D (**B**), Enge (**C**), and Darmanis (**D, E**) datasets. In **A, B,** BEDwARS has the least average RMSE at all noise levels (NL-X) as well as the Baron signature without added noise. Rodeo provided with BEDwARS-estimated proportions has similar performance. In **C**, there is a large performance gap between BEDwARS or RODEO/BEDwARS and other methods. In the deconvolution of brain transcriptomic profiles, RODEO provided with BEDwARS-estimated proportions has the least average RMSE using IP and its noisy versions (**D**) as well as other brain reference signatures except for CA (**E**).

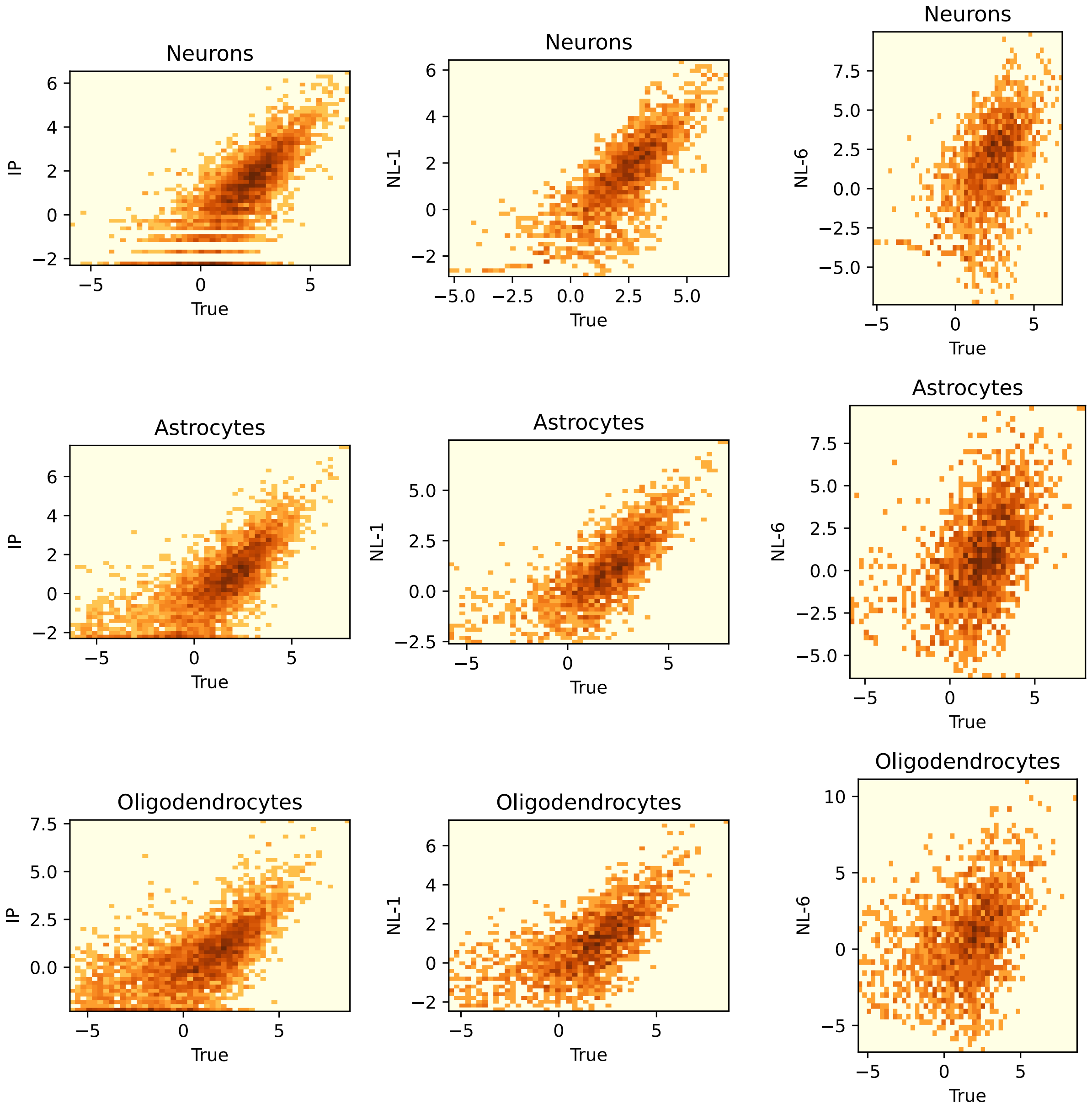


**Figure S16. Relationship of IP and its two noisy variants with the true Darmanis signature.** For each reference signature ("IP" and its two variants – "NL-1" and "NL-6", representing the lowest and highest levels of added noise respectively), each gene's expression value in that signature is shown on the y-axis and its expression in the true signature (Darmanis) is shown on the x-axis; both are in log scale. As the noise increases the signatures are less similar to the true signature. IP signature is highly diverged from the Darmanis signature.

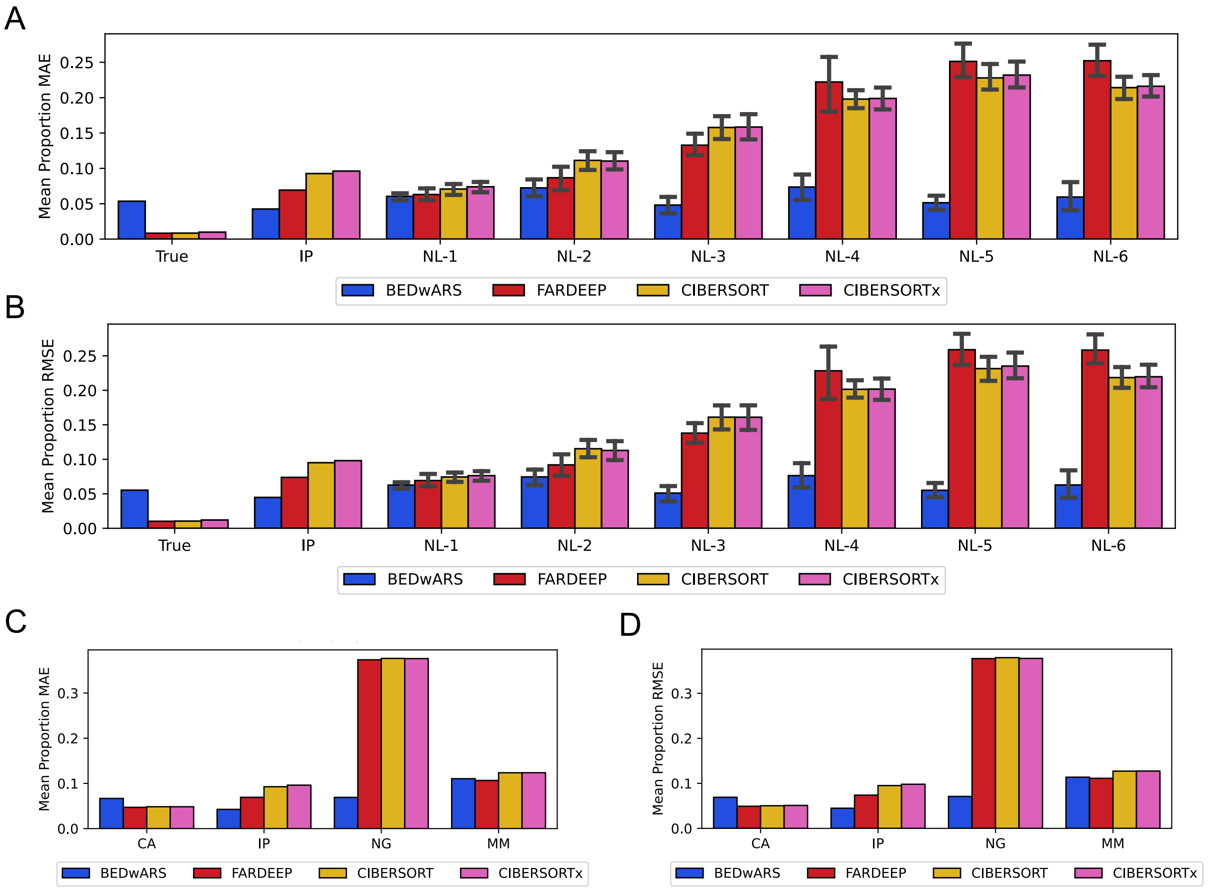


**Figure S17. Performance evaluation, by RMSE and MAE criteria, of methods for estimation of cell type proportions by deconvolving pancreatic and brain transcriptomic/pseudo-bulk profiles.** Shown are MAE (mean absolute error) (**A, C**) and RMSE (root mean squared error) (**B, D**) between estimated and true proportions, averaged over the cell types, for all four methods – BEDwARS (this work), FARDEEP [CITE], CIBERSORT [CITE], CIBERSORTx [CITE] – tested by us. RMSE and MAE are computed between the inferred and true cell type proportions of 100 pseudo-bulk samples derived from Darmanis dataset. Evaluations are shown with the IP reference signature as well as its noisy variants (NL-1, NL-2, … NL-6). BEDwARS has the least RMSE and MAE in IP group and all noise levels. Evaluations are also shown for the hypothetical case where the true underlying signature was available during deconvolution ("True" category in each panel), though this is not a common situation (**A, B**). Similarly, BEDwARS has the best performance by a large margin in both criteria using MM signature as the reference signature and it is close to other methods using CA and MM (**C, D**).

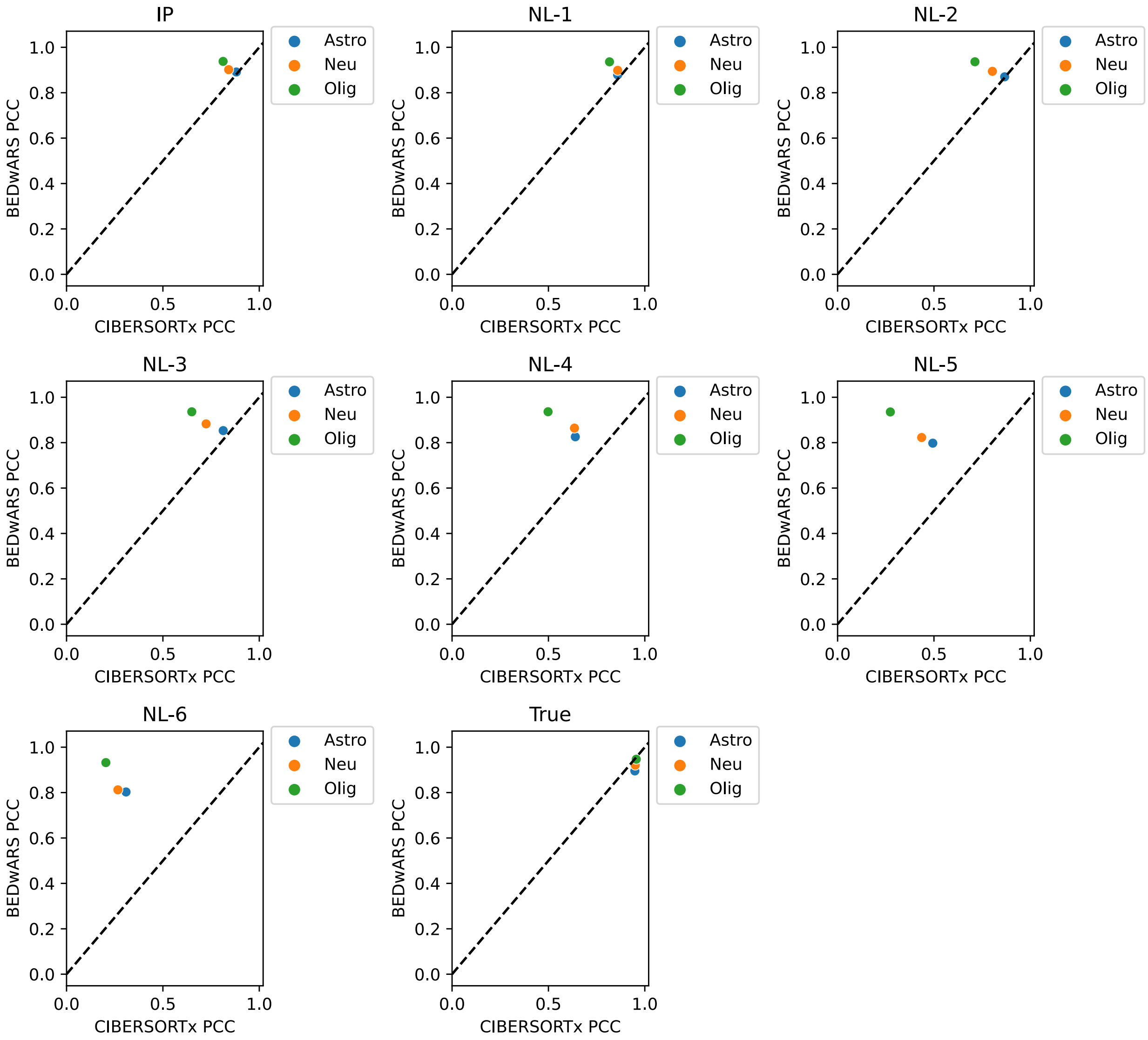


**Figure S18. Cell type level comparison of BEDwARS and CIBERSORTx for the task of proportion estimation by deconvolution of Darmanis pseudo-bulk samples.** Different panels correspond to different reference signatures used during deconvolution -- the IP signature and its noisy variants NL-1, NL-2, … NL-6, as well as the true signature. Performance is measured by the Pearson Correlation Coefficient (PCC) between estimated and true proportions of a cell type, across the 100 pseudo-bulk samples. For evaluations with noisy signatures, the average over 11 noisy signatures at the same noise level is shown. BEDwARS outperforms or performs similarly to CIBERSORTx in most of the cases.

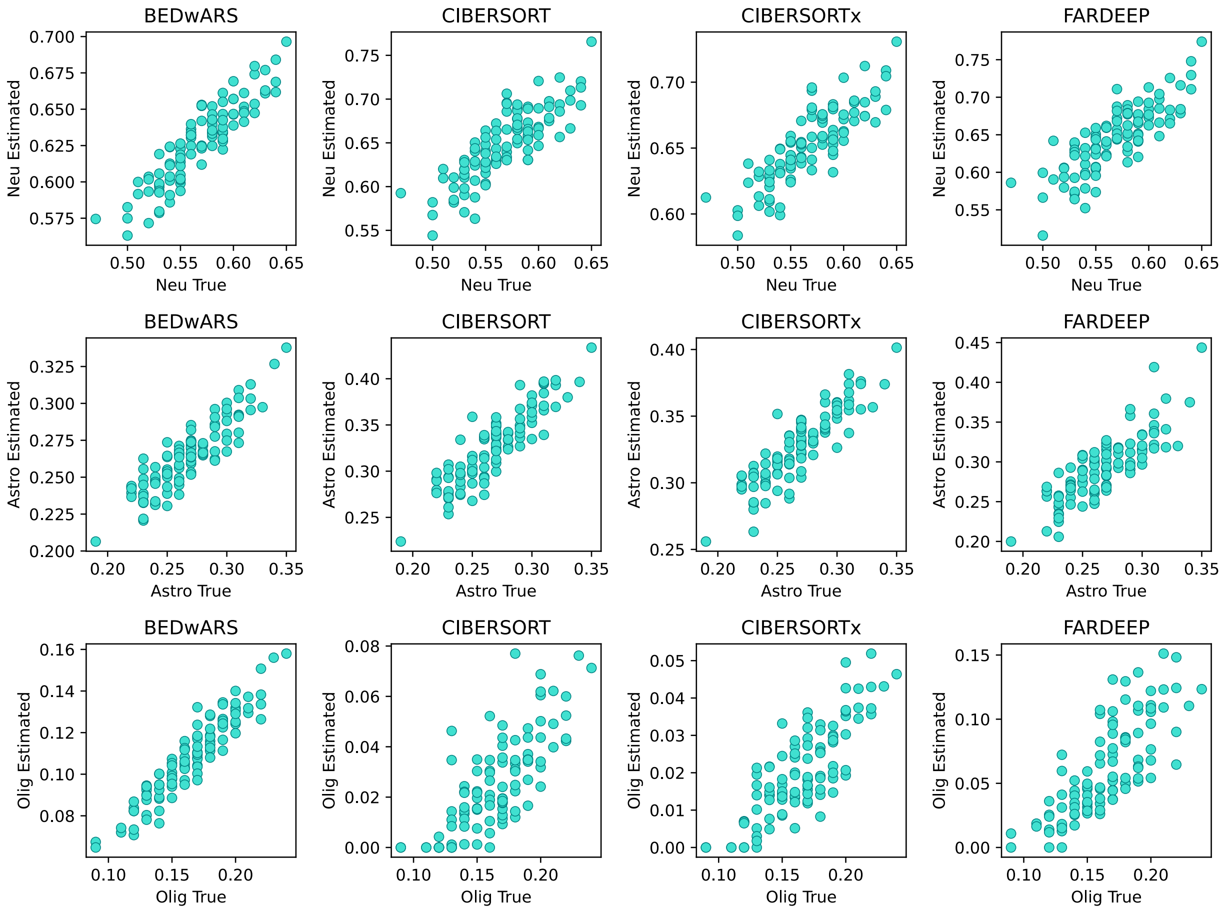


**Figure S19. Quality of cell type proportion inference using IP signature in deconvolving pseudo-bulk samples from Darmanis.** The estimated and true proportions for each of 100 pseudo-bulk samples are compared, for all cell types and all methods (BEDwARS, CIBERSORT, CIBERSORTx, FARDEEP). Overall, BEDwARS-estimated proportions have higher correlation with true proportions than other methods. The performance gap between BEDwARS and other methods is larger for oligodendrocytes, for which CIBERSORT(x) underestimates the highest proportions by nearly 10-fold. Neurons, astrocytes and oligodendrocytes are abbreviated as Neu, Astro, and Olig, respectively.

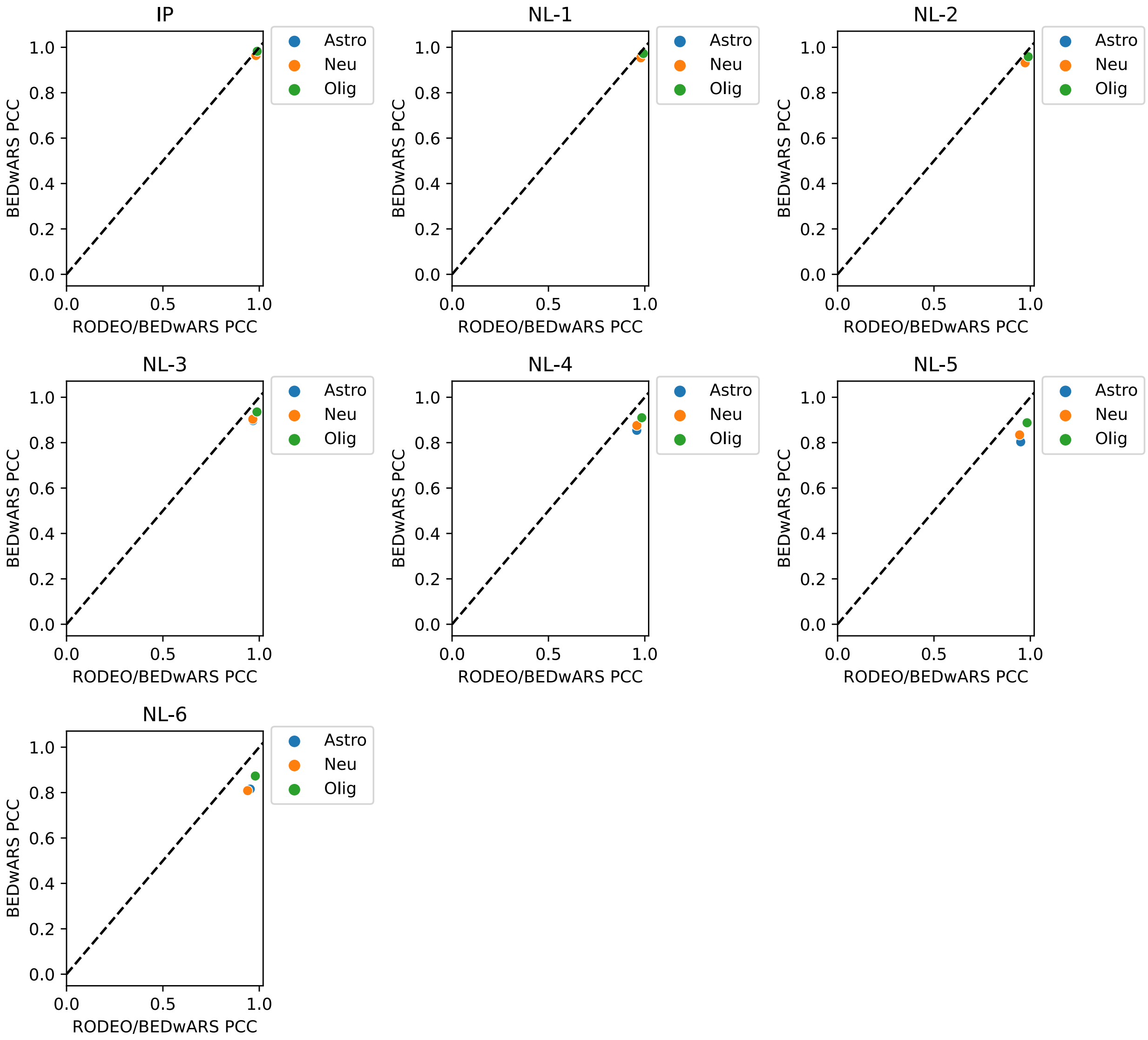


**Figure S20. Cell type level comparison of BEDwARS and RODEO/BEDwARS for the task of signature estimation by deconvolution of Darmanis pseudo-bulk samples.** Different panels correspond to different reference signatures used during deconvolution -- the IP signature and its noisy variants NL-1, NL-2, … NL-6. Performance is measured by the Pearson Correlation Coefficient (PCC) between estimated and true signatures of a cell type. For evaluations with noisy signatures, the average over 11 noisy signatures at the same noise level is shown. For the low noise levels and IP signature the cell type level performance of BEDwARS and RODEO/BEDwARS are similar. However, as the noise level increases RODEO/BEDwARS provides better estimation of cell type signatures.

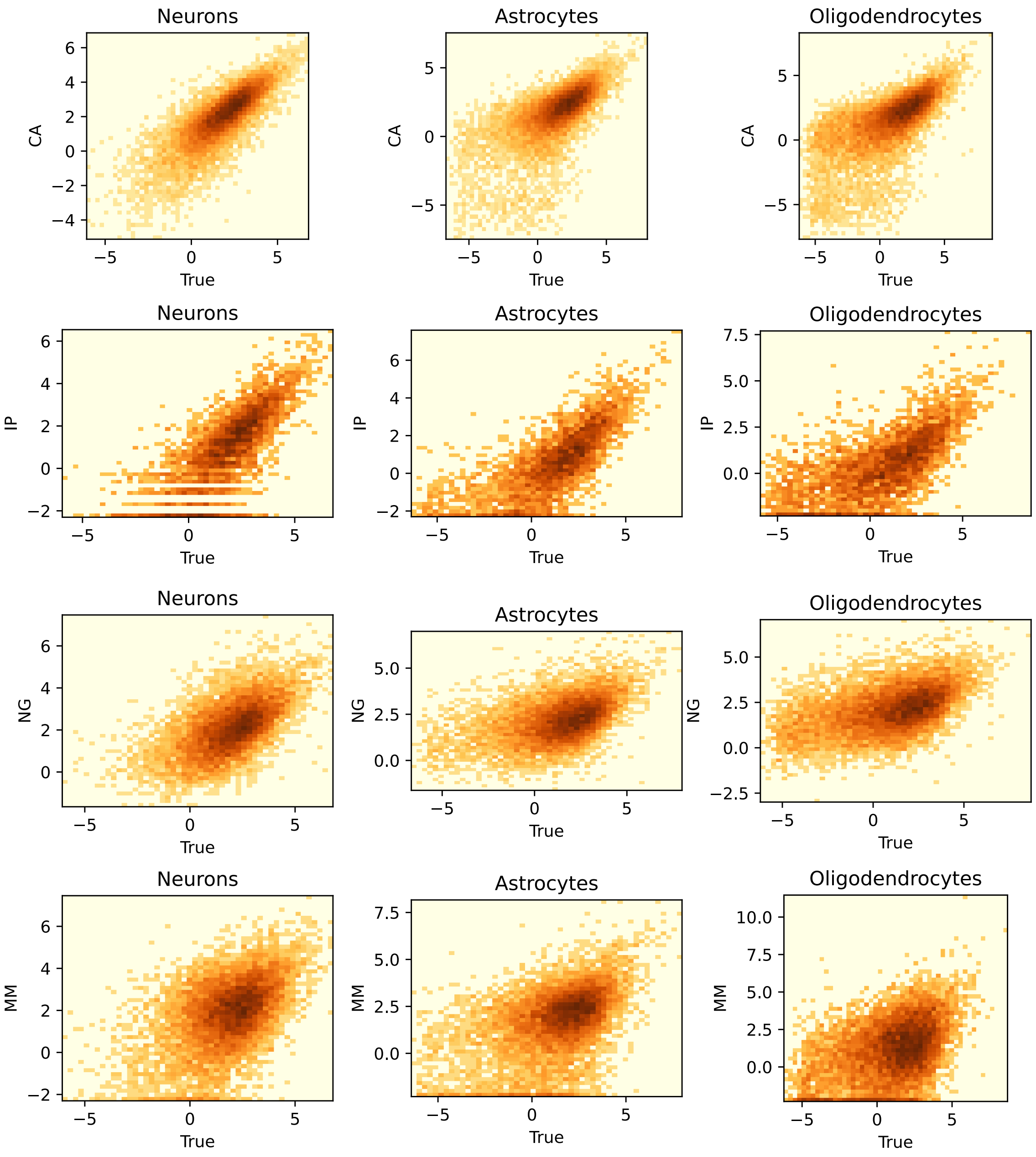


**Figure S21. Relationship of different brain reference signatures with the true Darmanis signature.** For each reference signature (CA, IP, NG, and MM), each gene's expression value in that signature is shown on the y-axis and its expression in the true signature (Darmanis) is shown on the x-axis; both are in log scale. CA is the most similar to the Darmanis signature whereas MM is the most dissimilar one. The MM signature is generated from mouse brain gene expression. NG signature is also highly diverged from the true signature, which may be attributed to it being derived from a different region (prefrontal cortex) of brain than the Darmanis (temporal middle gyrus) signature.

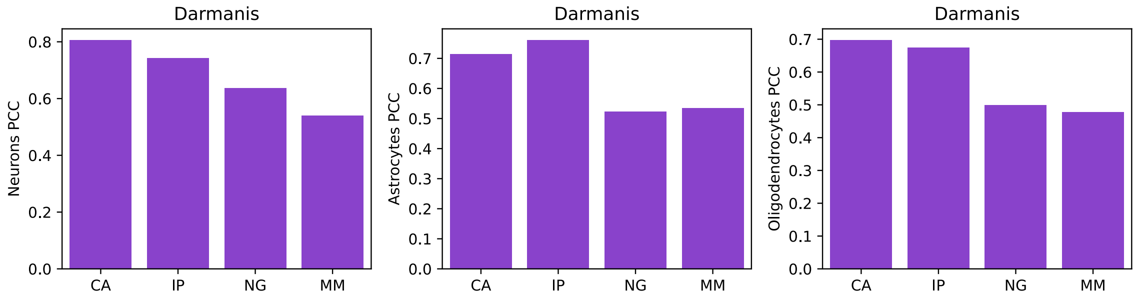


**Figure S22. Correlation between the true Darmanis signature and reference brain signatures from multiple sources.** For each cell type, Pearson correlation coefficient (PCC) is shown between the log-transformed reference and true signatures. Overall, CA and MM are the most similar and dissimilar respectively to the Darmanis signatures. CA: adult single-nuclease RNA-seq of human brain from Cell Atlas, IP: RNA-seq data on immune-purified cells from human brain, NG: single-nuclease RNA-seq of human brain (Nagy et al.), MM: RNA-seq data on immune-purified mouse brain.

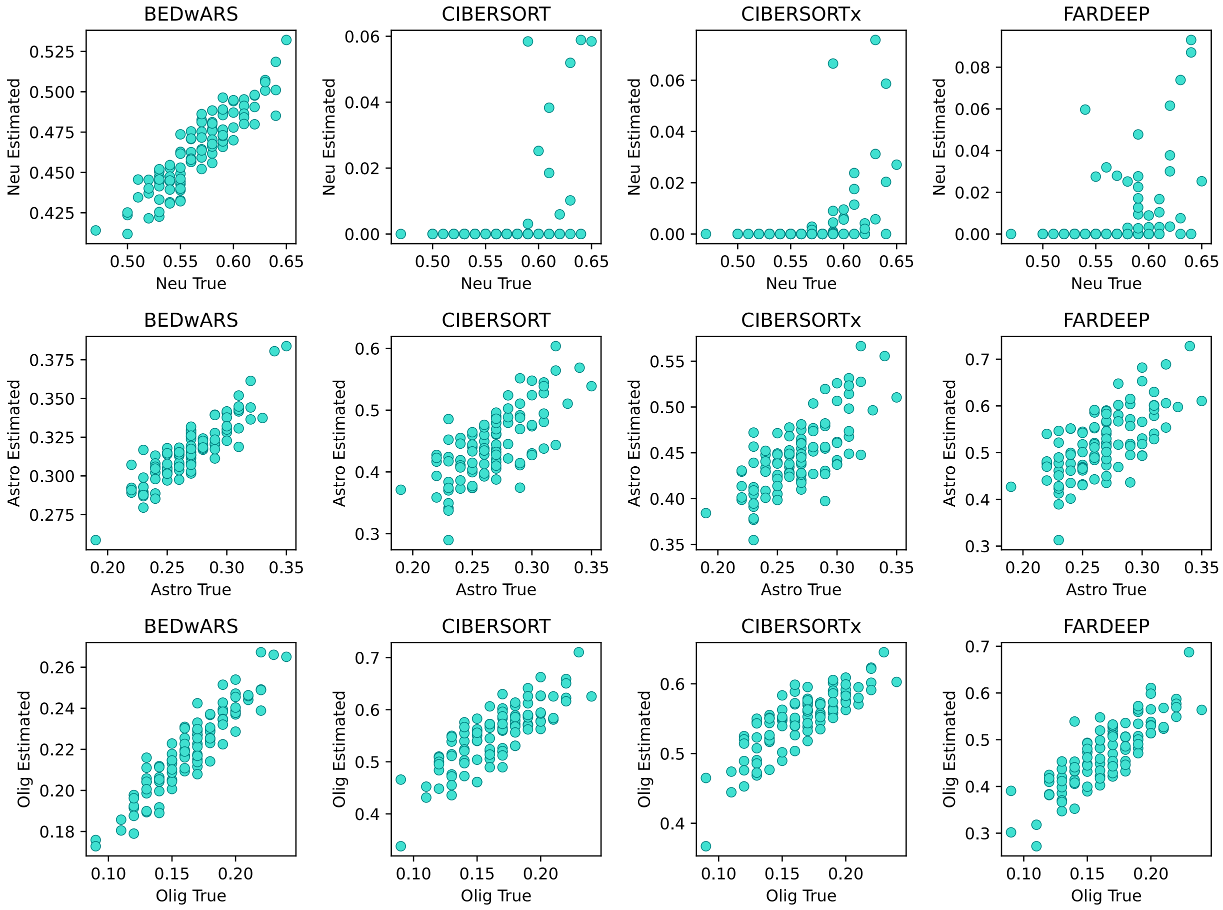


**Figure S23. Quality of cell type proportion inference using NG signature in deconvolving pseudo-bulk samples from Darmanis.** The estimated and true proportions for each of 100 pseudo-bulk samples are compared, for all cell types and all methods (BEDwARS, CIBERSORT, CIBERSORTx, FARDEEP). All methods except BEDwARS fail to predict neuron proportion for most of the samples. For astrocytes, the estimated values of the highest proportions are almost twice as large as true values, for all methods except BEDwARS. The same holds for oligodendrocytes, where other methods overestimate the highest proportions by ~ 3-fold. Note that the NG signatures are derived from a different region of brain than the Darmanis signatures.

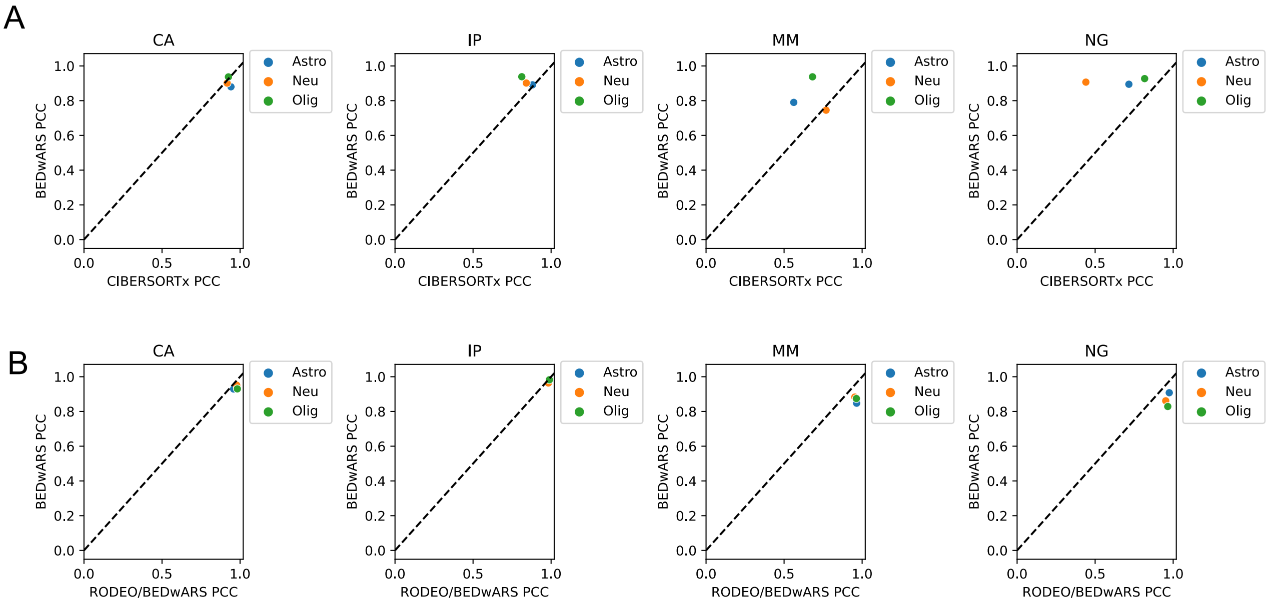


**Figure S24. Cell type level comparison of (BEDwARS, CIBERSORTx) and (BEDwARS, RODEO/BEDwARS) for proportion and signature estimation by deconvolution of Darmanis pseudo-bulk samples using multiple brain reference signatures.** Different panels correspond to different reference signatures used during deconvolution -- CA, IP, MM and NG. **A.** Performance is measured by the Pearson Correlation Coefficient (PCC) between estimated and true proportions of a cell type, across the 100 pseudo-bulk samples. Performance of BEDwARS and CIBERSORTx are similar in estimating cell type proportions when using reference signatures that are similar to the true signature (CA, IP). For the more deviated signatures (MM, NG) BEDwARS has clear advantage over CIBERSORTx. **B.** PCC is computed between estimated and true signatures of a cell type for signature estimation. Performance of RODEO provided with BEDwARS-estimated proportions is similar to BEDwARS (CA, IP) or slightly better than BEDwARS (MM, NG).

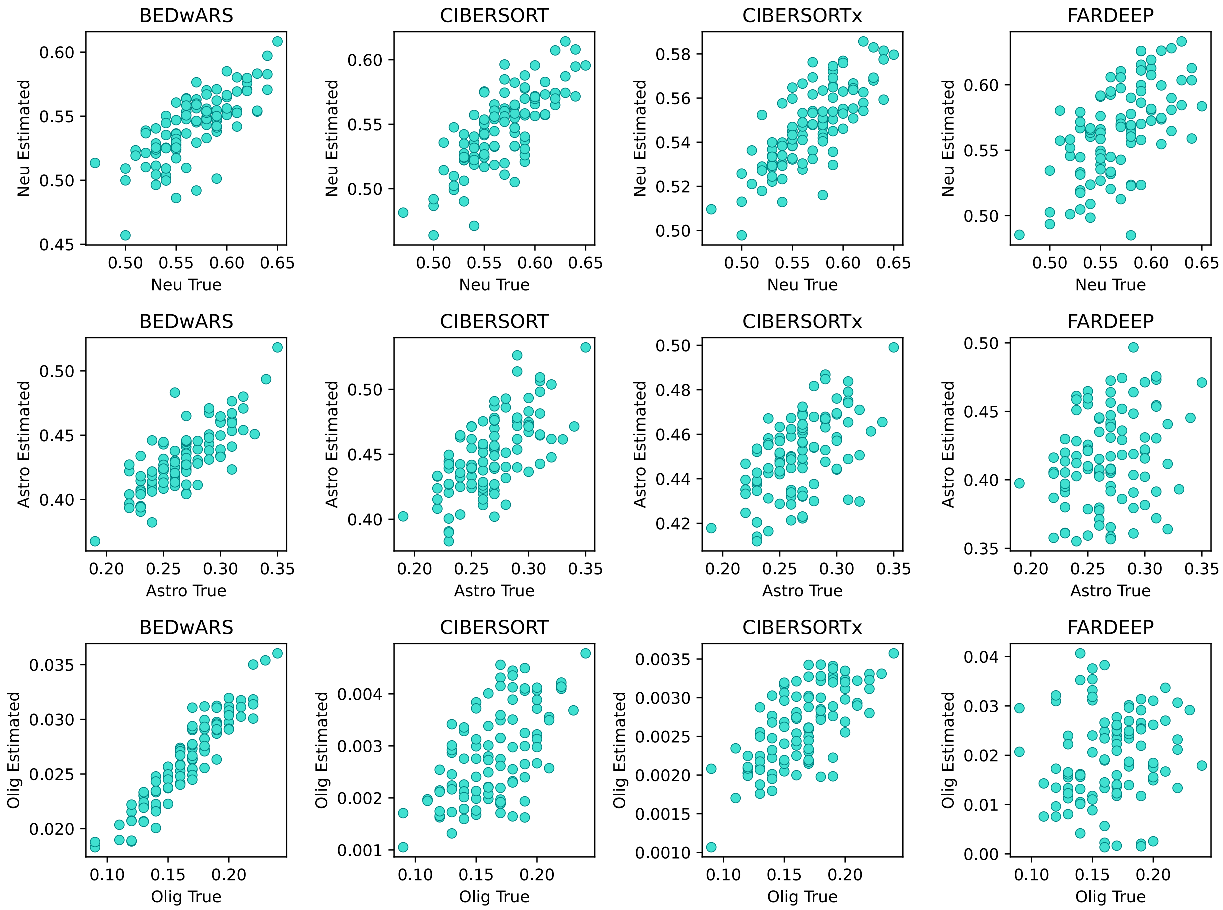


**Figure S25. Quality of cell type proportion inference using MM signature in deconvolving pseudo-bulk samples from Darmanis.** The estimated and true proportions for each of 100 pseudo-bulk samples are compared, for all cell types and all methods (BEDwARS, CIBERSORT, CIBERSORTx, FARDEEP). Overall BEDwARS-estimated proportions are best in capturing the trend of true proportions, especially for oligodendrocytes. While the maximum estimated proportion is ~10-fold (BEDwARS and FARDEEP) and ~100-fold (CIBERSORT(x)) less than the true proportion, BEDwARS estimates have significantly higher correlation for this cell type. A cell type signature estimation method such as RODEO does benefit from estimations highly correlated with true values even if the absolute values do not match perfectly.

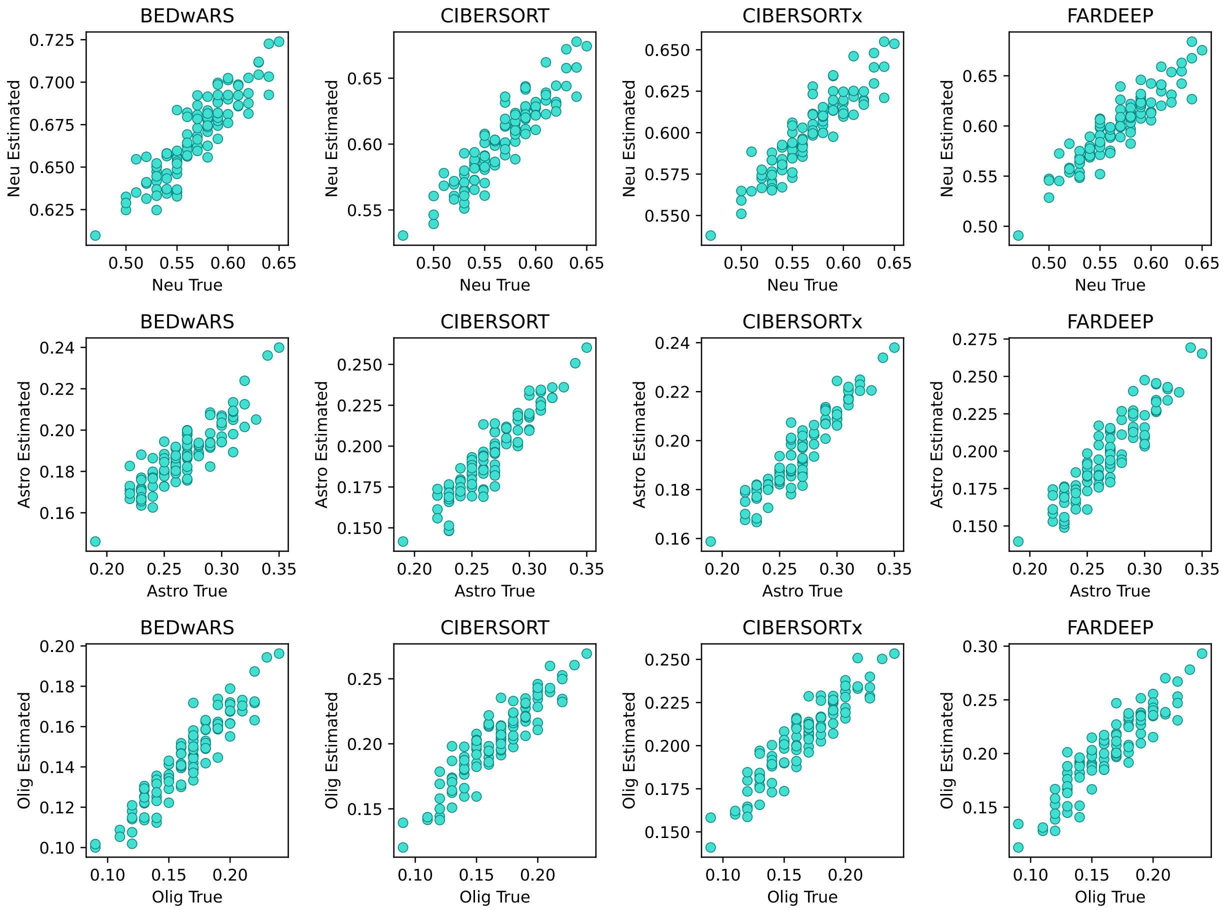


**Figure S26. Quality of cell type proportion inference using CA signature in deconvolving pseudo-bulk samples from Darmanis.** The estimated and true proportions for each of 100 pseudo-bulk samples are compared, for all cell types and all methods (BEDwARS, CIBERSORT, CIBERSORTx, FARDEEP). All methods perform almost equally well in estimation of proportions.

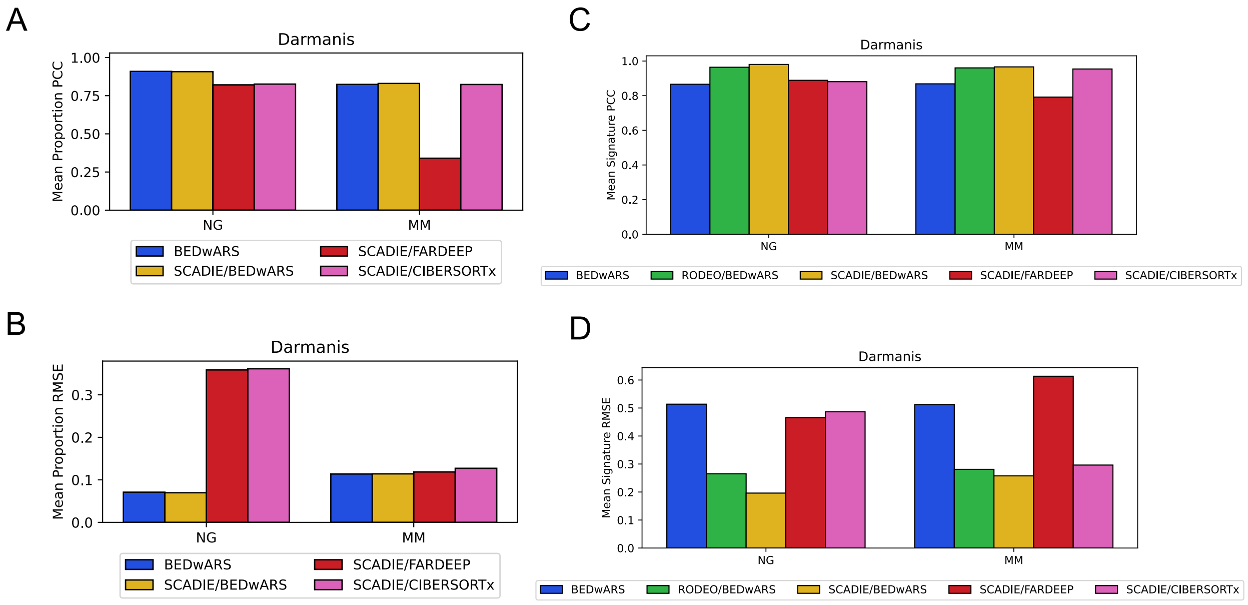


**Figure S27. Evaluation of cell type proportion and signature estimation by Pearson Correlation Coefficient and RMSE for SCADIE.** Pseudo-bulk profiles of Darmanis dataset were deconvolved with SCADIE using two reference signatures – NG and MM. Performance of SCADIE was evaluated with three different deconvolution methods – BEDwARS, FARDEEP and CIBERSORTx – in its first step, and compared with BEDwARS for cell type proportion (**A, B**) and with BEDwARS and RODEO/BEDwARS for cell type signature estimation (**C, D**). For the estimation of cell type proportion with NG signatures, SCADIE/BEDwARS is competitive with BEDwARS but SCADIE/FARDEEP and SCADIE/CIBERSORTx are not competitive. For the estimation of cell type signatures starting with NG signatures, SCADIE/BEDwARS is competitive with RODEO/BEDwARS but SCADIE/FARDEEP and SCADIE/CIBERSORTx are not. SCADIE's performance in the inference of cell type proportions and signatures is strongly dependent on the deconvolution method used for the initialization of cell type proportions in its first step. For example, FARDEEP shows the lowest quality of proportion estimates by PCC criterion using MM signature (**Figure 4C)** and SCADIE/FARDEEP (i.e., SCADIE with FARDEEP deconvolution in its first step) likewise has the lowest average correlation for the estimation of cell type proportions (A). Similarly, BEDwARS has the highest quality of cell type proportion estimates by both PCC and RMSE criterion using NG signature, and SCADIE initialized with BEDwARS-estimated proportions has similar performance to BEDwARS (A).

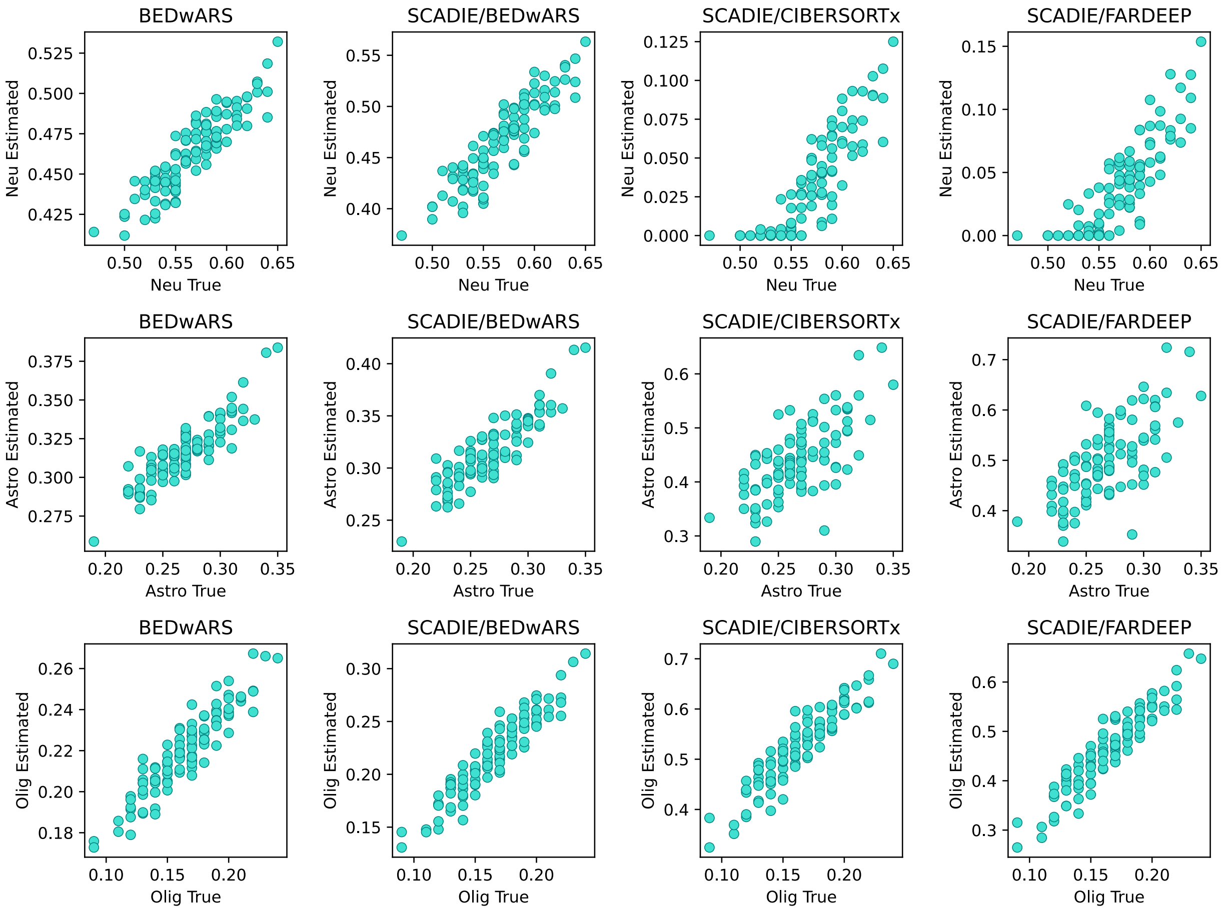


**Figure S28. Quality of cell type proportion inference using NG signature in deconvolving pseudo-bulk samples from Darmanis for BEDwARS and SCADIE.** SCADIE was evaluated with three different deconvolution methods – BEDwARS, CIBERSORTx, FARDEEP – for its first step (denoted by SCADIE/BEDwARS, SCADIE/CIBERSORTx, SCADIE/FARDEEP). The estimated and true proportions for each of 100 pseudo-bulk samples are compared, for all cell types and methods. Quality of estimated proportions by SCADIE are affected by the deconvolution method used for the initialization of cell type proportions. In astrocytes and oligodendrocytes SCADIE estimated proportions are almost two and three times larger than the true proportions when initialized with CIBERSORTx and FARDEEP. Neuron proportions are extremely underestimated as well. Similar patterns exist in **Figure S23** for CIBERSORTx and FARDEEP. BEDwARS and SCADIE/BEDwARS have similar quality of estimation and show the most accurate estimates for all cell types.

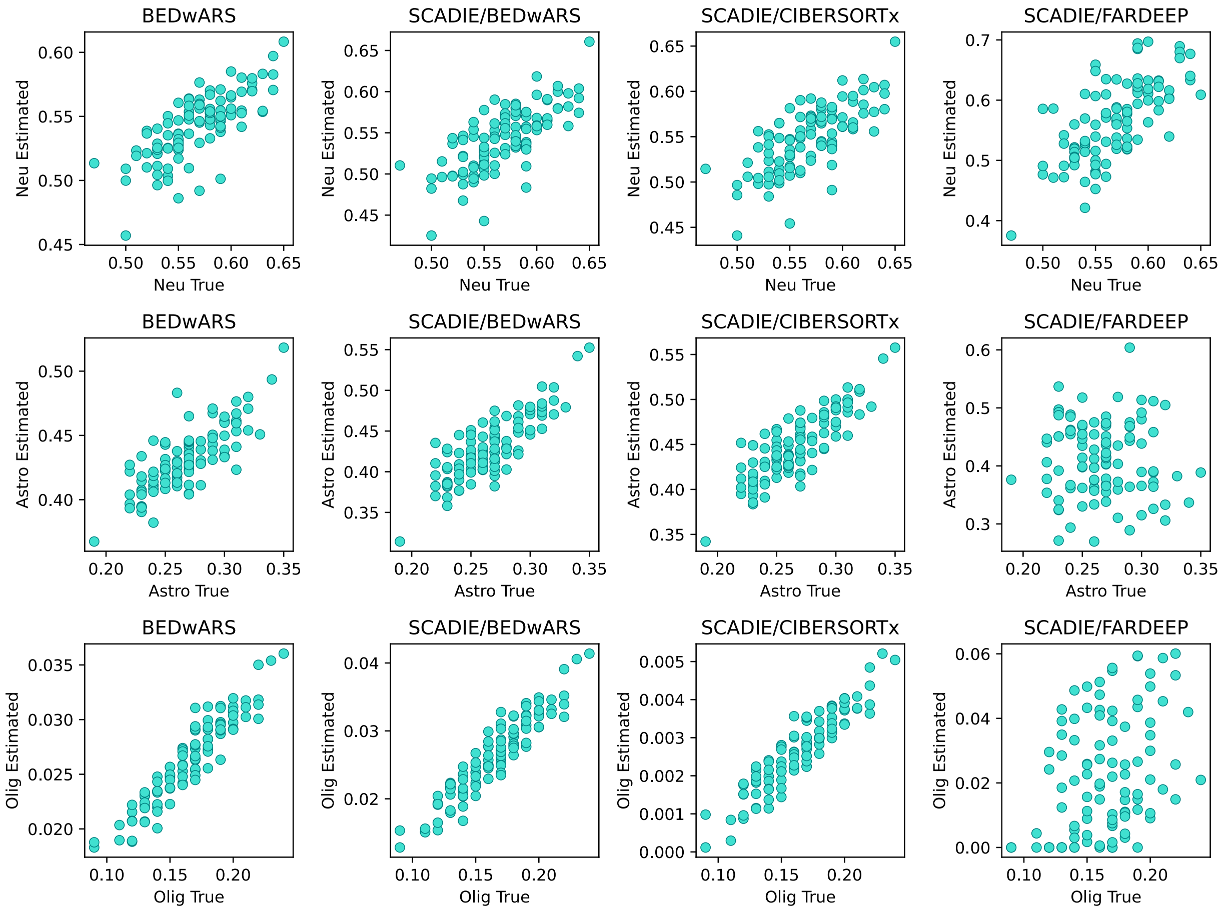


**Figure S29. Quality of cell type proportion inference using MM signature in deconvolving pseudo-bulk samples from Darmanis for BEDwARS and SCADIE.** The estimated and true proportions for each of 100 pseudo-bulk samples are compared, for all cell types and variants of SCADIE as well as BEDwARS. Quality of estimated proportions by SCADIE are affected by the deconvolution method used for the initialization of cell type proportions. For oligodendrocytes and astrocytes, the estimated proportions are poorly correlated with true correlations when FARDEEP is used for initialization. When initialized with CIBEROSRTx-estimated proportions the final proportions of oligodendrocytes reported by SCADIE are 100 times less than the true proportions even though they are highly correlated with each other. These observations are similar to the ones in Figure **S25** which confirms that SCADIE's performance is highly dependent on its initialization with other deconvolution methods. SCADIE initialized with BEDwARS has similar quality of estimates to BEDwARS and is more accurate than its other variants.

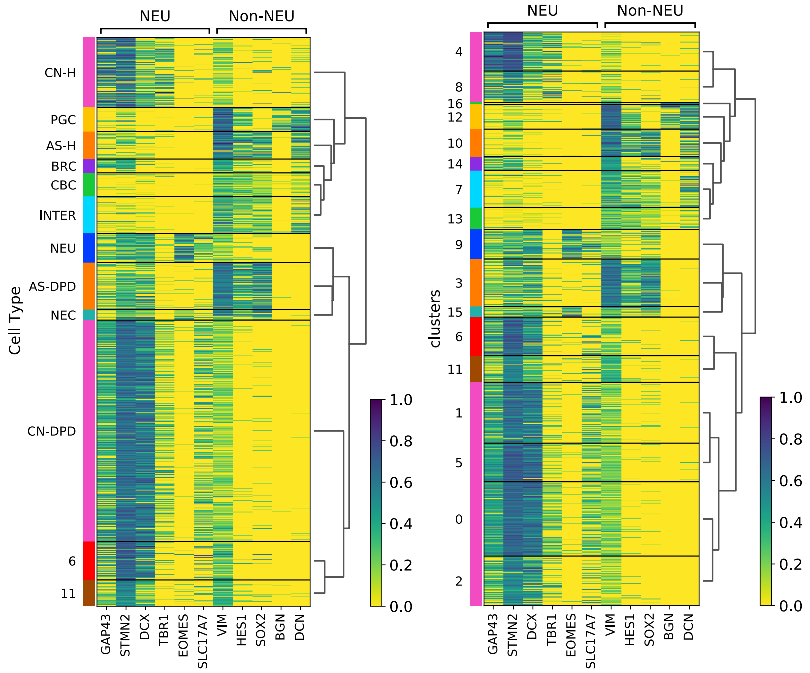


**Figure S30. Hierarchical clustering of the cells from DPD-deficient patient and the non-affected individual**. The standardized/scaled expression (divided by max in each dimension to fall in (0, 1) interval) of neuronal (GAP43, STMN2, DCX, TBR1, EOMES, SLC17A7) and non-neuronal (VIM, HES1, SOX2, BGN, DCN) gene markers are shown for the cells of DPD-deficient patient and non-affected individual. Hierarchical clustering was performed on the cells grouped by their assigned cell type (**left**) or their cluster index (**right**). Each cell type and all the cluster indices assigned to it are represented with the same color in both heatmaps. Cells from both individuals were pooled into a single set before clustering.

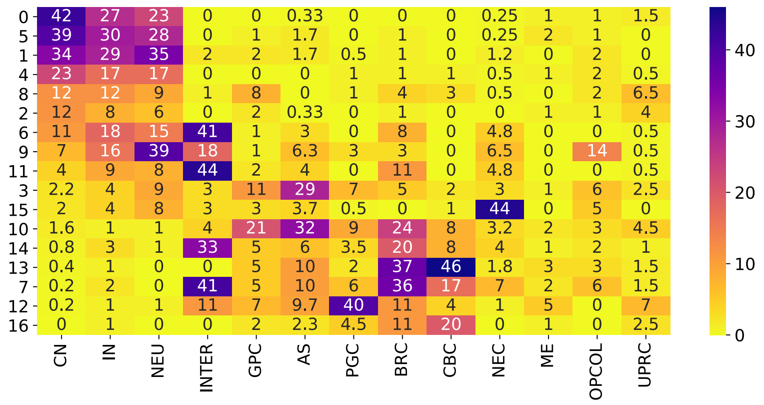


**Figure S31. Overlap of statistically derived markers between DPD deficiency dataset and Tanaka et al. (25) study.** For each cluster of cells (rows), shown is the overlap of its markers with the markers of a cell type from Tanaka et al. (columns). In cases where a cell type from Tanaka et al. has multiple subtypes such as CN1, CN2, …, CN5, the average overlap over the subtypes was computed.

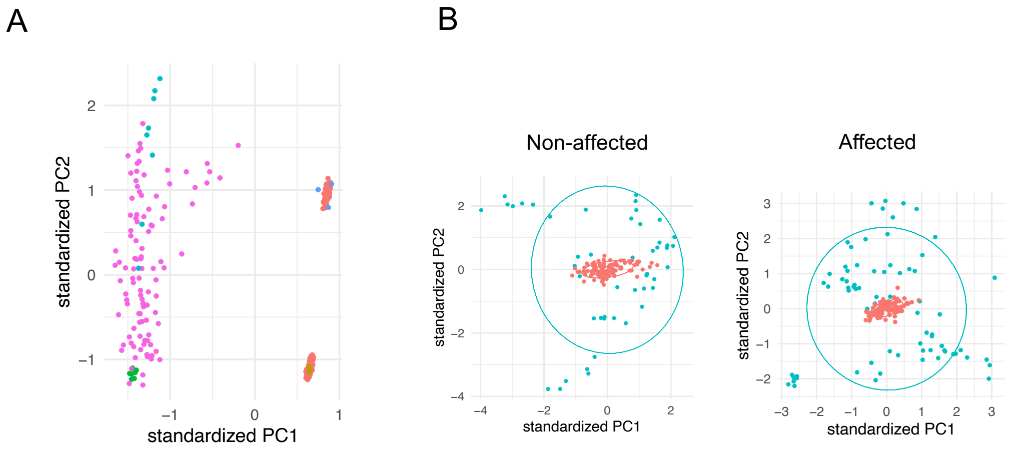


**Figure S32. Batch correction for bulk RNA-seq samples of affected and non-affected groups of individuals. A.** PCA plot of bulk RNA-seq and pseudo-bulk samples used in the study of DPD deficiency. Points on the left (pink, teal and green) are the bulk RNA-seq samples from non-affected (48) and affected individuals (72). The teal and green points represent the eight semi-matched bulk samples of the non-affected and affected groups. Points on the right are the pseudo-bulk samples for each group. Red points represent pseudo-bulk samples generated by bootstrapping from single cell data for non-affected (top) and affected (bottom) groups. The blue and brown points are the pseudo-bulk samples generated by summing the expression of cells within each organoid. **B.** The bulk samples (teal) of non-affected and affected groups are batch corrected to the bootstrapped pseudo-bulk samples (red) of non-affected and affected groups separately.

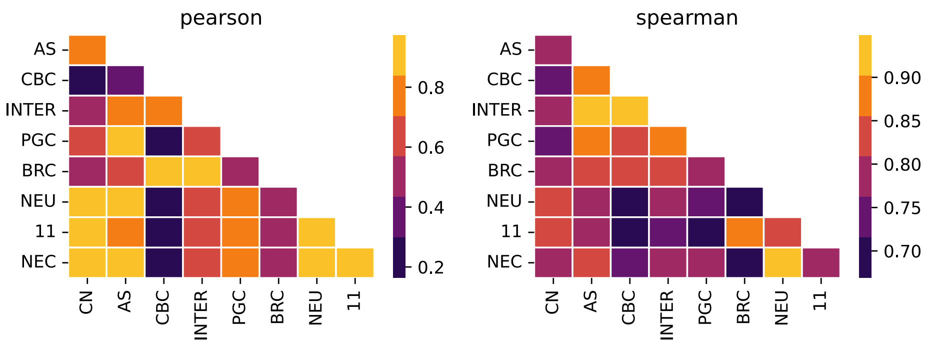


**Figure S33. Pairwise similarities of cell type signatures used for deconvolving affected and non-affected bulk samples by DPD deficiency.** Pearson (left) and Spearman (right) correlation coefficients are shown for each pair of cell types. CBC has the highest correlation with INTER by both statistics. Similarly, CN is highly correlated with neuron (NEU), NEC, and cluster 11 by both criteria.

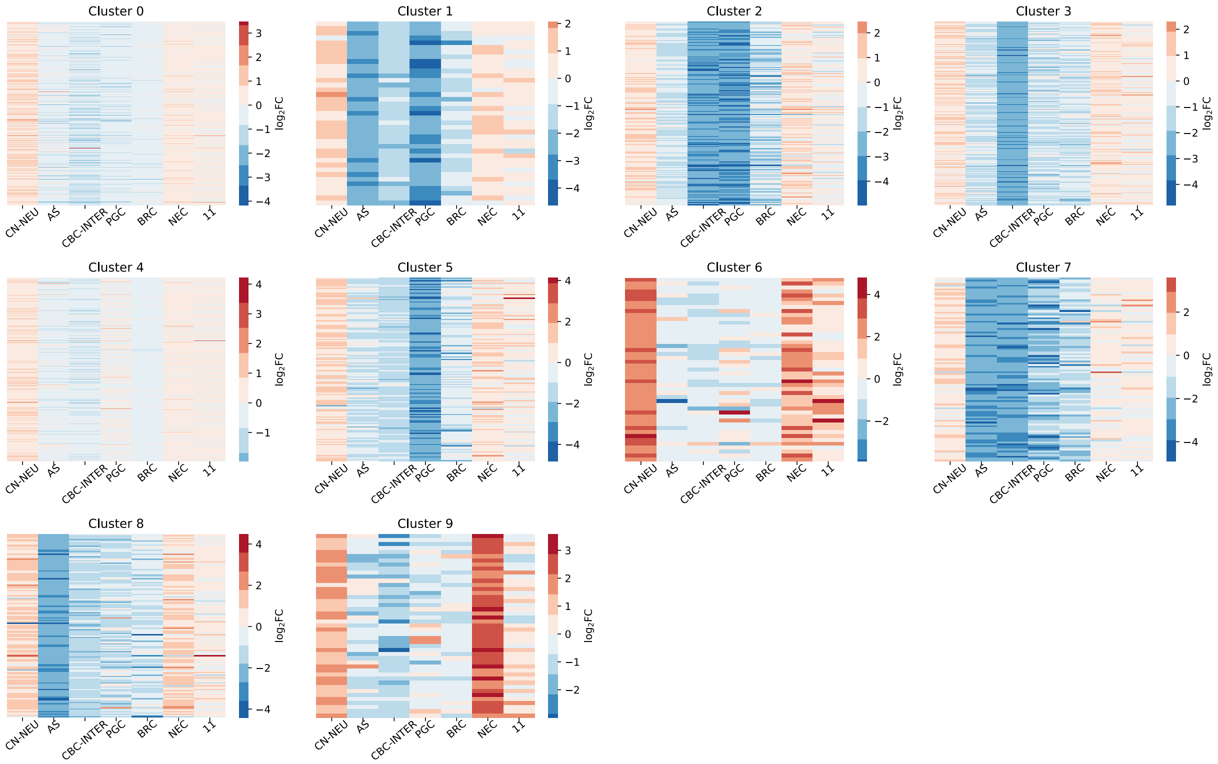


**Figure S34. Kmeans clustering of differential expression patterns across cell types.** Each gene is assigned a cell type-specific expression fold change between affected and non-affected individuals based on results from BEDwARS deconvolution of bulk RNA-seq data. A gene's (log) fold-change in each of 7 cell types (two of these represent pairs of cell types) is its differential expression pattern. Genes' expression patterns are clustered into ten clusters using K-means. Shown are the differential expression patterns of genes in each resulting cluster.

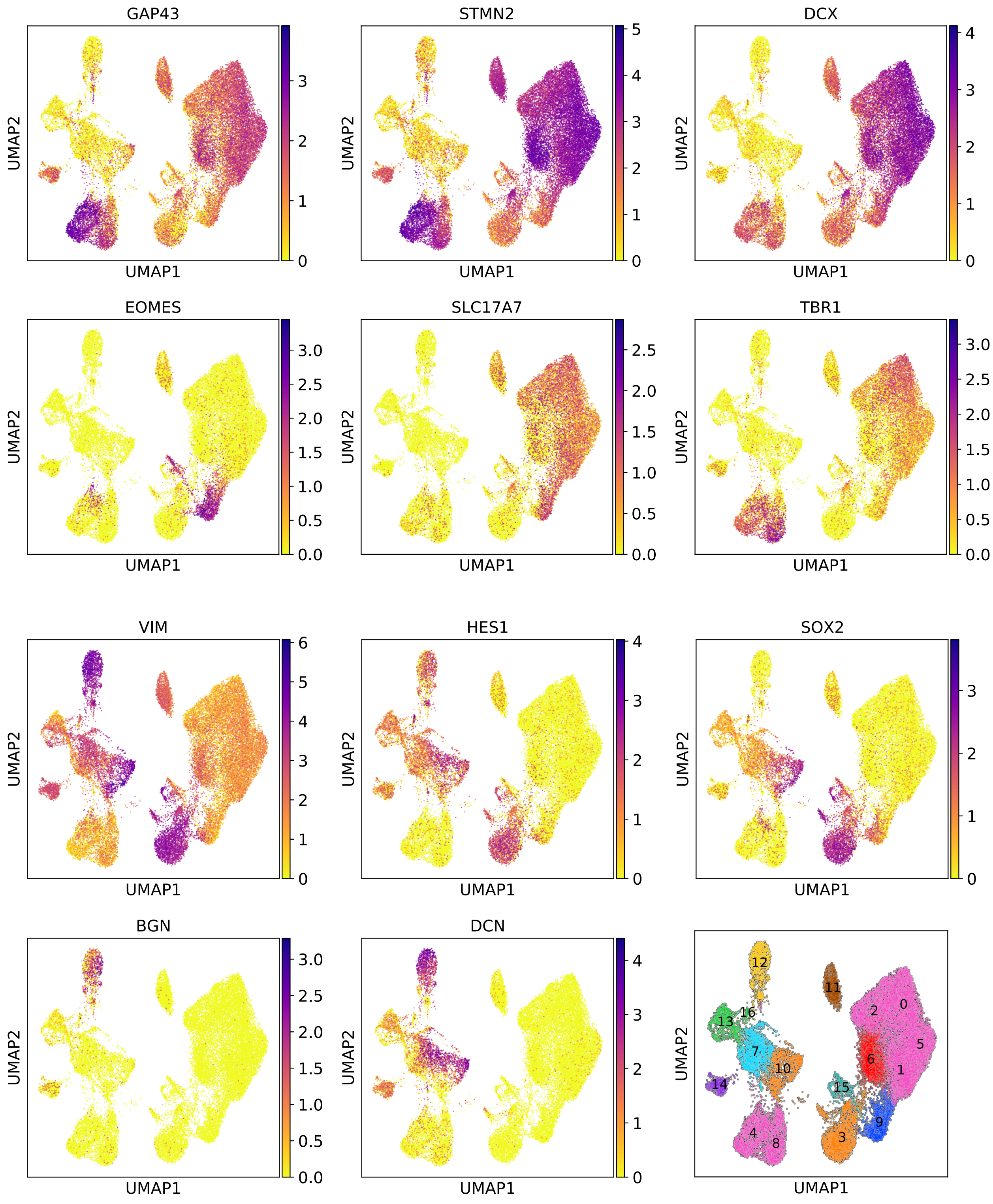


**Figure S35. Distribution of the expression of marker genes over the cells.** The normalized and log-transformed expression of neuronal (GAP43, STMN2, DCX, TBR1, EOMES, SLC17A7) and non-neuronal (VIM, HES1, SOX2, BGN, DCN) marker genes represented in the UMAP plots of the DPD-deficient patient and non-affected individual. The last panel shows the cluster indices on cell clusters. Clusters with the same color were assigned to the same cell type.

| Cluster Index | Expressed Markers | Marker Overlap | GO Enrichment | Cluster type | Assigned Cell Type |
|---|---|---|---|---|---|
| 0 | CN | CN | - | Affected | CN |
| 1 | CN | CN | - | Affected | CN |
| 2 | CN | CN | - | Affected | CN |
| 3 | Non-neuronal | AS | AS | Affected | AS |
| 4 | CN | CN | - | Non-Affected | CN |
| 5 | CN | CN | - | Affected | CN |
| 6 | CN | INTER | - | Affected | unassigned |
| 7 | Non-neuronal | INTER/BRC | - | Non-Affected | INTER |
| 8 | CN | CN | - | Non-Affected | CN |
| 9 | Neuron | Neuron | - | Affected | Neuron |
| 10 | Non-neuronal | AS | AS | Non-Affected | AS |
| 11 | Inconclusive | INTER | - | Non-Affected | unassigned |
| 12 | PGC | PGC | - | Non-Affected | PGC |
| 13 | Non-neuronal | CBC | CBC | Non-Affected | CBC |
| 14 | Non-neuronal | INTER/BRC | - | Non-Affected | BRC |
| 15 | Non-neuronal | NEC | NEC | Affected | NEC |
| 16 | Non-neuronal | CBC | CBC | Non-Affected | CBC |

**Table S1.** Cell type assignment to cell clusters for the DPD deficient patient and the non-affected individual for whom scRNA-seq data were available. Each cluster of cells is assigned to a cell type based on the agreement of at least two out of three criteria used. First criterion (Expressed Markers) is the average expression of neuronal and non-neuronal genes used by Tanaka et al., second criterion (Marker Overlap) is the overlap of the statistically derived markers of our clusters and the cell types of Tanaka et al., and the third criterion (GO Enrichment) is the enrichment of statistically derived markers in relevant GO terms. Cluster type specifies whether most cells belong to the affected individual or non-affected individual.

**Table S2. Differential gene expression analysis for DPD deficiency.** The summary of differential gene expression analysis using Limma package for bulk, pseudo-bulk, and deconvolved cell type or pairs of cell types (CN-NEU, CBC-INTER) expression profiles (**"DGE"**). Top 200 DE genes per cell type or pairs of cell types are listed in **"Top 200 DE genes per cell type"** sheet**.**

| | |
|---|---|
| GO:0000785 | chromatin |
| GO:0005576 | extracellular region |
| GO:0005615 | extracellular space |
| GO:0005683 | U7 snRNP |
| GO:0005739 | mitochondrion |
| GO:0005743 | mitochondrial inner membrane |
| GO:0005747 | mitochondrial respiratory chain complex I |
| GO:0005758 | mitochondrial intermembrane space |
| GO:0005783 | endoplasmic reticulum |
| GO:0005788 | endoplasmic reticulum lumen |
| GO:0005794 | Golgi apparatus |
| GO:0005852 | eukaryotic translation initiation factor 3 complex |
| GO:0016282 | eukaryotic 43S preinitiation complex |
| GO:0016607 | nuclear speck |
| GO:0032543 | mitochondrial translation |
| GO:0032870 | cellular response to hormone stimulus |
| GO:0034663 | endoplasmic reticulum chaperone complex |
| GO:0035976 | transcription factor AP-1 complex |
| GO:0042612 | MHC class I protein complex |
| GO:0042824 | MHC class I peptide loading complex |
| GO:0051082 | unfolded protein binding |
| GO:0098869 | cellular oxidant detoxification |
| GO:1990837 | sequence-specific double-stranded DNA binding |

**Table S3. GO term names for the GO IDs in Figure 5E.**

**Table S4. Summary of David gene set characterization performed on top 200 DE genes identified by DGE analysis for bulk, bootstrapped pseudo-bulk and deconvolved cell type expression profiles for DPD deficiency.** In each annotation cluster the first GO with significant FDR (FDR < 0.05), highlighted with yellow, was considered.

**Table S5. Cluster of genes identified by Kmeans clustering based on the pattern of genes' differential expression across the cell types for DPD deficiency.** Each column represents the genes belonging to the same cluster.

**Table S6. Summary of David gene set characterization for cluster of genes identified by Kmeans algorithm based on the pattern of genes' differential expression across the cell types for DPD deficiency.** The David results for each cluster are included in a sheet named with the cluster name (Cluster X). Clusters zero and four were excluded as they had more than 2000 genes. David results for top 200 DE genes identified by DGE analysis on bulk and bootstrapped pseudo-bulk are included for convenient comparison.

**Table S7. Top 200 markers per cluster of non-affected and affected cells for DPD deficiency.** Each column contains the top 200 filtered marker genes for a cluster of cells. These markers were used for assigning cell types to the clusters.

| Cluster Index | GO Term | $-\log_{10}(\text{pvalue})$ |
|---|---|---|
| 3 | astrocyte differentiation | 2.7 |
| 10 | astrocyte differentiation | 4.5 |
| 15 | mitotic cell cycle | 17.7 |
| | mitotic chromosome condensation | 10.6 |
| | regulation of mitotic cell cycle | 5.8 |
| | regulation of mitotic nuclear division | 2.8 |
| 13 | motile cilium | 4 |
| | epithelial cilium movement | 3.6 |
| 16 | cilium movement | 9.7 |
| | motile cilium | 7.6 |
| | cilium assembly | 4.7 |
| | motile cilium assembly | 4.3 |

**Table S8. Enrichment of statistically derived cell cluster markers for gene ontology terms used by Tanaka et al. in cell type assignment.** Significance of the GO terms enriched in top 200 statistically derived marker genes for a subset of cell clusters from affected and non-affected individuals. The enrichment of these GO terms was used in assigning cell type to clusters.

**Table S9. KnowEng gene set characterization performed on the markers of a subset of cell clusters for DPD deficiency.** These results were used to assign cell types to cluster of non-affected and affected cells. The GO terms that were enriched and used for cell type assignment are summarized in **Table S8** for easier lookup.
